# Lumen charge governs gated ion transport in β-barrel nanopores

**DOI:** 10.1101/2024.09.26.615172

**Authors:** Simon Finn Mayer, Marianna Fanouria Mitsioni, Paul Robin, Lukas van den Heuvel, Nathan Ronceray, Maria Jose Marcaida, Luciano A. Abriata, Lucien F. Krapp, Jana S. Anton, Sarah Soussou, Justin Jeanneret-Grosjean, Alessandro Fulciniti, Alexia Möller, Sarah Vacle, Lely Feletti, Henry Brinkerhoff, Andrew H. Laszlo, Jens H. Gundlach, Theo Emmerich, Matteo Dal Peraro, Aleksandra Radenovic

## Abstract

β-barrel nanopores are involved in crucial biological processes, from ATP export in mitochondria to bacterial resistance, and represent a promising platform for emerging sequencing technologies. However, in contrast to ion channels, the understanding of the fundamental principles governing ion transport through these nanopores remains in its early stages. In this study, we integrate experimental, numerical, and theoretical approaches to elucidate ion transport mechanisms in these biological nanopores. We identify and characterise two distinct nonlinear phenomena: open-pore rectification and gating. Through extensive mutation analysis of aerolysin nanopores, we demonstrate that open-pore rectification is caused by ionic accumulation driven by the distribution of lumen charges. Additionally, we provide converging evidence suggesting that gating is controlled by electric fields dissociating counterions from lumen charges, promoting local structural deformations. Our findings establish a rigorous framework for the characterisation and understanding of biological ion transport processes, enabling the design of adaptable biosensors. We illustrate this by optimizing an aerolysin mutant for computing applications, paving the way for novel nanofluidic technologies.

## Introduction

Pore-forming proteins (PFPs) are biological nanofluidic channels enabling the regulated or free passage of molecules and ions across membranes. They are essential for information transmission, import and export, as well as attack and defence of cells and organisms^1^. Out of their biological context, some PFPs have been repurposed as sensors for probing a diverse range of molecules^2–9^. These sensors are known as biological nanopores and have been commercialised for low-cost, long-read DNA sequencing^8^, a first demonstration of their potential for societal impact. Despite evident technological interest, ionic motion through biological nanopores still awaits proper rationalisation. It is well known that ion transport through such biological channels is highly nonlinear^10,11^. In particular, the ionic current through such pores can be polarity-dependent, an effect known as rectification^12–14^, and it can abruptly decrease, a phenomenon referred to as gating^11,15–23^. The mechanisms underlying such nonlinearities, particularly the role of lumen charge, remain poorly understood, hindering further progress in biological nanopore sensing technologies. This contrasts with solid-state nanofluidics, where advances in nanofabrication and theoretical modelling have enabled in-depth rationalisation of ion transport in confinement^24^ and with ion channels, where decades of advances in molecular biology have shed light on a rich mechanistic phenomenology^10^. Biological nanopores represent a grey area, where such characterization is more challenging due to their heterogeneous surface charge, and their intermediate size around 1 nm, that precludes a strictly molecular or continuous understanding. Rationalizing ion transport in these pores is nonetheless of paramount importance due to their pivotal role in living organisms as well as their potential for disruptive technology.

In this work, we aim at rationalising ion transport in biological pores taking into account the complexity of their lumen charge as well as their mechanical properties. We then harness this knowledge to uncover the physical origin of biological nanopore gating. We focus on multimeric β-barrel-forming nanopores widely used in the nanopore field (SI Figure 3) such as the toxins aerolysin and α-hemolysin (α-HL) as well as *Mycobacterium smegmatis* porin A (MspA). They are to be distinguished from smaller highly specific ion channels that often undergo conformational changes to enable or disable transport of a specific species of ion^10^. To capture the complexity of the lumen charge, we study the dependence of gating on 26, 6 and 5 different mutants of aerolysin, α-HL and MspA, respectively. Such pores have constriction diameters between around 1 and 1.4 nm and a length of roughly 10 nm^9,25^. Our study combines single-pore and ensemble transmembrane ion transport measurements, biophysical modelling, atomistic molecular dynamics (MD) simulations and cryo-electron microscopy (cryo-EM) characterization. We show how the charge landscape of the pore lumen controls ion transport across the open pore. We then present evidence that this landscape impacts voltage-induced mechanical deformations resulting in gating, and build a model of nanopore gating based on these observations, using electrostatics and statistical mechanics. Finally, we harness this knowledge to mimic synaptic potentiation using a carefully selected mutant of aerolysin.

### Non-linear responses of biological nanopores: rectification and gating

To show the non-linear responses of biological nanopores, we start by studying the response of single aerolysin wild type (wt) pores to a constant applied potential. Figure 1a illustrates our experimental setup, where a single nanopore is embedded into a lipid membrane separating two electrolytic solutions containing two silver/silver-chloride electrodes. A voltage bias is applied on the electrodes and the resulting ionic current across the nanopore is measured. The membranes are freshly folded^15^ or painted^26^ and, by fine tuning the nanopore concentration, we have control over the number of nanopores inserted in the membrane.

**Figure 1.**
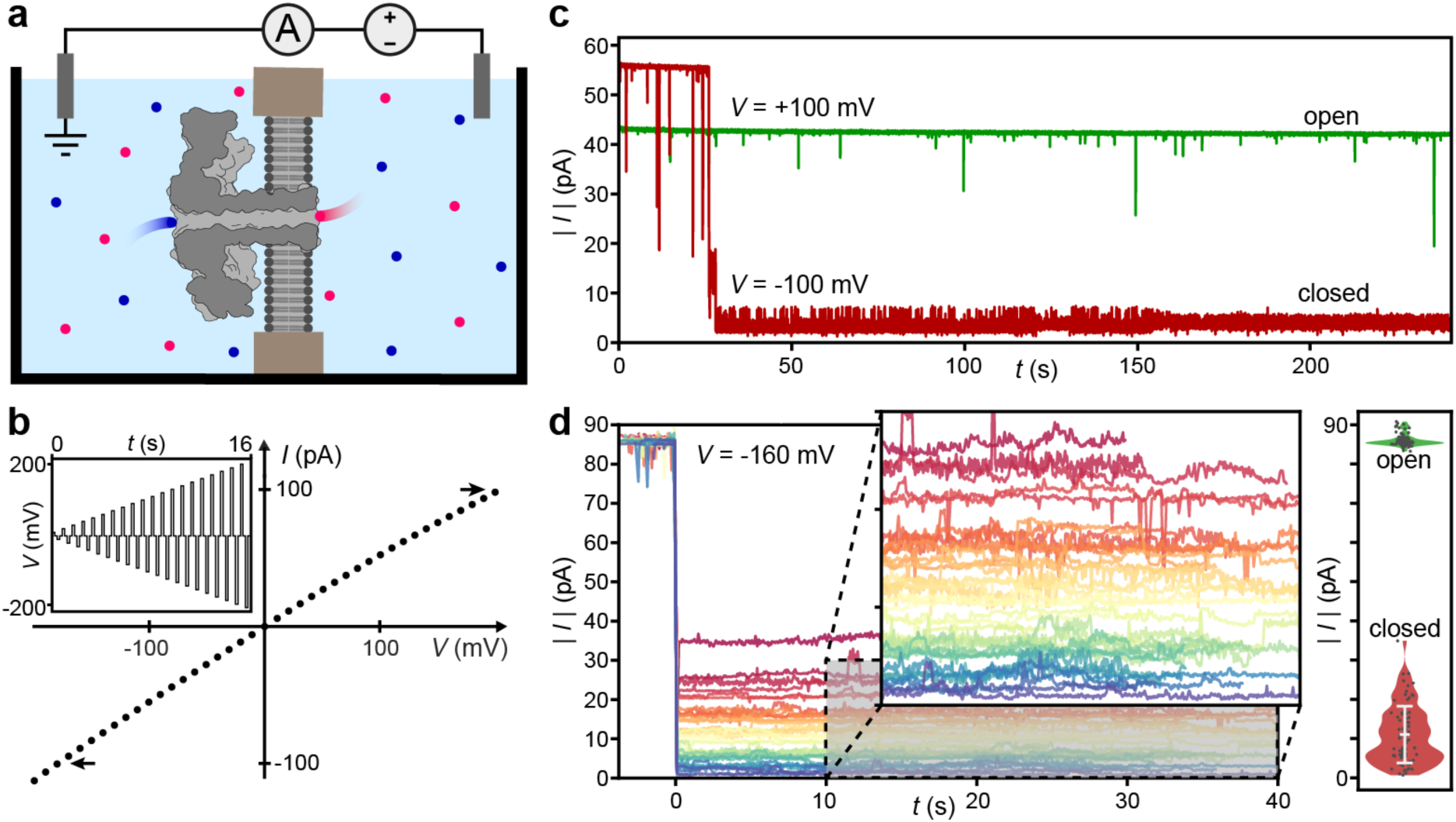
Complexity and diversity of ion transport in single aerolysin pores. **a.** Not-to-scale schematic of the experimental setup. A lipidic membrane separating two reservoirs filled with electrolyte (cations are shown in blue, anions are shown in red) with a single aerolysin wild type (wt) pore incorporated, the current through the pore is measured via Ag/AgCl electrodes. **b.** Typical single pore *IV* curve where the current of the open pore is measured at different voltages consecutively. A slight rectification is visible and indicated through arrows at ±100 pA. The inset shows the voltage trace that can be used to acquire such data. **c.** Absolute current over time traces at constant potential of a single aerolysin wt nanopore. The same pore is measured first at 100 mV (green) and then at-100 mV (red) for 240 seconds. Rectification is clearly visible as the absolute current is higher at-100 mV. At 100 mV the measured current is constant while at-100 mV the current decreases spontaneously, this behaviour is referred to as gating. **d. (left)** 39 gating traces recorded for a single aerolysin wt pore at-160 mV, coloured by current, showing the diversity of gating states associated with different current levels. The overlay is a zoom-in of the area shaded in grey **(right)** Violin plot of gating (red) and open-pore (green) current levels measured in 4 different single pores at 160 mV. Each randomly offset point in the violin plot represents the average of the gating or open-pore current of one gating trace. The white bar denotes the average and standard deviation of the average closed-state currents. The large closed-state current variation relative to the open-pore current variation points to a stochastic phenomenon. All experiments are done in 1 M KCl buffered to pH 6.2 with 10 mM phosphate.

When probed at different voltages for a few seconds at a time in order to obtain the *IV* curve, a small rectification is observed (Figure 1b), meaning that the pore conductance at +100 mV differs from that at-100 mV. We then applied a constant potential of both opposite signs for a few minutes in order to monitor the pore stability (Figure 1c). We observed that after a few tens of seconds, the negative current trace suddenly reduces by more than 90%. This abrupt decrease is the signature of gating^11^ and it hinders analyte sensing applications where a stable baseline current trace is required^9,27^. Over the last decades, a number of experimental studies aimed at characterising and understanding the gating phenomenon in β-barrel biological nanopores have been conducted and several hypotheses have been proposed, most convincingly conformational changes, among others^15–18,11,19–23^. However, no compelling evidence has been presented yet. Aerolysin, as well as other biological pores, thus exhibit two distinct nonlinearities. The first takes the form of an open-pore rectification, while the second is a delayed, sharp, and polarity-dependent decrease in current, referred to as gating.

To investigate this behaviour further we recorded the same single aerolysin wt nanopore over several hours and found that despite the fact that the current reduces to a discrete state or switches between several discrete states, these states are diverse in their average current. We proceeded as follows: we applied a voltage of-160 mV until the pore gated, then removed the applied voltage after a few tens of seconds, before reapplying the same-160 mV bias. As shown in Figure 1d it is rare that the same state is observed in different gating events. Overall, we recorded 4 different pores – no clearly defined separated states emerged from this large sample (Figure 1d). The low throughput of single-pore direct current (DC) measurements combined with the stochastic and diverse nature of gating makes this approach unsuitable for extensive characterization of the phenomenon. This calls for a novel approach to understand this behaviour.

Biological pores exhibit history-dependent conductivity as they switch from the open to the closed state after approximately 30 seconds in the displayed trace (Figure 1c). In other words they behave as resistors with an internal variable sensitive to the applied voltage. Such devices are called memristors (resistors with memory). Intensively studied in electronics^28–30^, memristive behaviour has recently been reported in solid-state^31–34^ and biological nanofluidic devices^20,21^. As is standard in memristor studies, we quantified the similarly nonlinear dynamics of our biological nanopores through alternating current (AC) measurements of the *IV* curves.

### Open-pore rectification and gating characterised with time-varying voltage

We used a periodic potential bias to measure and quantify both the open-pore rectification and the gating in a robust and rapid manner. For the open-pore rectification, triangular forcing was applied across a single pore at a frequency high enough for gating-free cycles to be observed (∼Hz) (Figure 2a). The rectification of a given pore is then quantified as a unitless value between-1 and 1 by the rectification factor *β* = (*I_+_*-|*I_-_*|)/(*I_+_* +|*I_-_*|) where *I_±_* = *I*(*V* = ±100 mV). To investigate gating, we chose a low-frequency sinusoidal span to maximise the time spent at higher voltages and therefore the closed-state probability.

**Figure 2.**
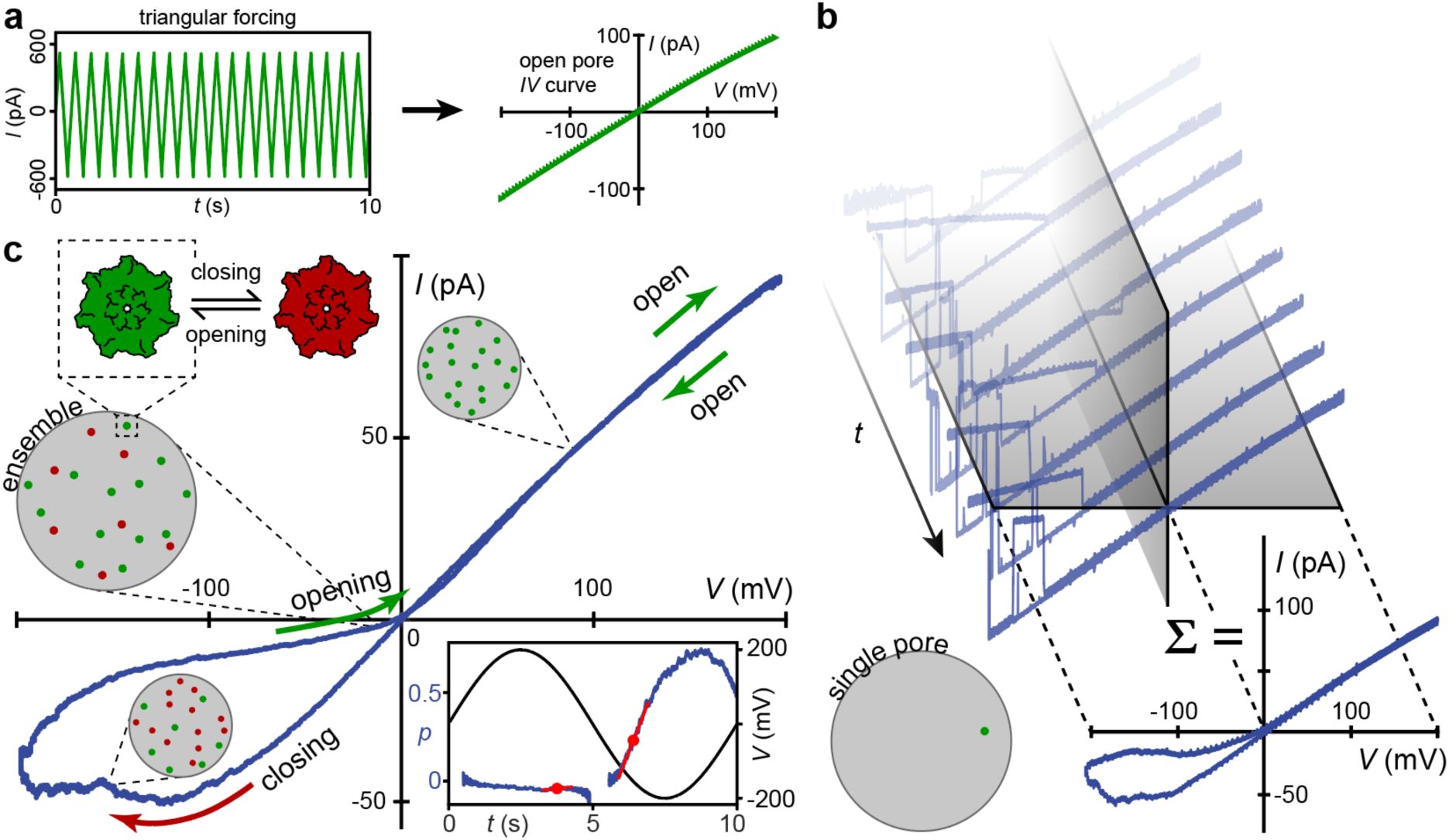
Methodology for the characterization of nonlinearities using time-varying voltage drop across single and multi-pore membranes. **a.** Triangular forcing at 2 Hz, 200 mV, with six aerolysin wt nanopores used to extract the single open-pore *IV* curve. b. *IV* characteristics of a single aerolysin wt nanopore in the lipid membrane measured with sinusoidal potential (0.1 Hz). 50 consecutive cycles are summed to retrieve the smooth hysteresis loop. **c.** normalised *IV* characteristics of 26 pores in the same membrane showing the smooth hysteresis loop (0.1 Hz). Pores in the membrane are closing and opening in a voltage-dependent way, giving rise to the hysteresis loop. Arrows indicate the direction of the sweep. The inset shows how we quantify gating by dividing the measured current by the single open-pore *IV* to retrieve the closed-state probability *p*, the solid black line shows the sinusoidal voltage. The calculation of *p* is ill-defined around the origin, resulting in a discontinuous line in the probability plot which did not affect our conclusions (see methods Section 4). The maximum slope indicated by the red lines and dots corresponds to *k_X_*/*ω* for both polarities where *k_X_* is the maximum closing rate and ω the forcing frequency. All experiments were conducted in 1 M KCl buffered with 10 mM phosphate to pH 6.2.

**Figure 3.**
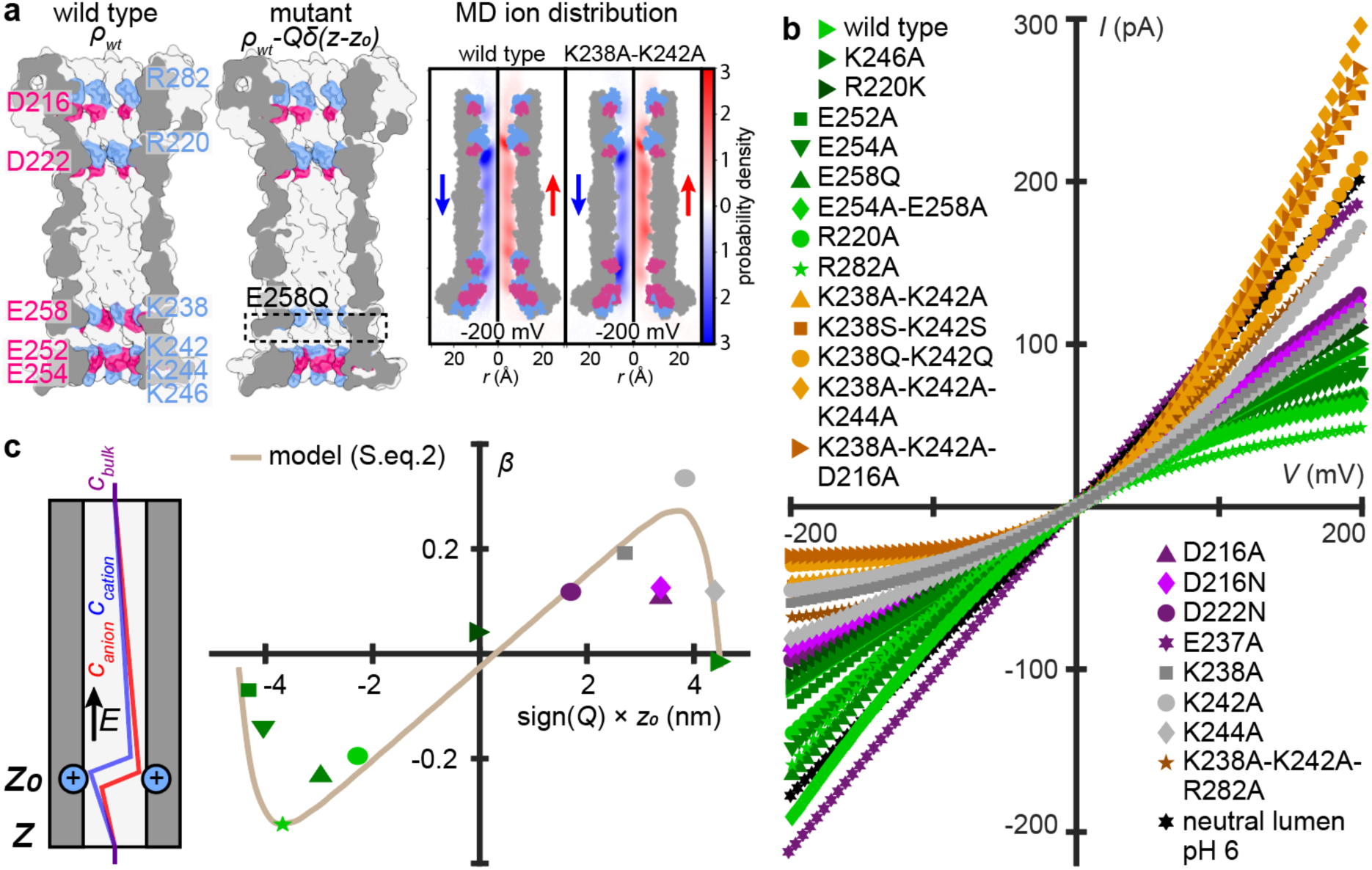
Characterization of open-pore current rectification using a wide library of aerolysin mutants. a. (left) Sketches of aerolysin wt and mutant E258 showing the deletion of a charged residue; Positive charges are drawn in blue and negative charges are drawn in pink. **(right)** ion radial probability distributions (blue for K^+^, red for Cl^-^) measured on the MD simulations of aerolysin wt and mutant K238A-K242A at-200 mV. **b.** Open-pore *IV* curves of several mutants of aerolysin showcasing the different degrees of rectification tuned by the lumen charge (1 M KCl, 10 mM phosphate pH 6.2, 1 Hz). **c. (left)** Schematic representation of mutant E258Q in the rectification model. The deletion of E258 which is negatively charged is represented as the addition of a fixed positive charge in the pore’s energy landscape, leading to ion accumulation downstream of the fixed charge. **(right)** Rectification factor *β* as function of the sign and position of the charge deleted after mutation. Solid curve: theoretical model (SI Equation 2)

When this low-frequency voltage was applied, the single aerolysin wt pore switched between open and closed states in the previously described stochastic, voltage-dependent way, the pore remained in the open state at positive biases and closed at negative applied biases. The stochasticity resulted in significant variations of the *IV* curve at negative voltage between cycles (Figure 2b). When 50 consecutive cycles of the same pore are averaged the characteristic memristive hysteresis becomes clearly visible, in the form of a loop in the *IV* curve (Figure 2b). Analysing this stochastic phenomenon thus requires a high number of repetitions in order to extract its average behaviour. We found that measuring a single sweep of multiple pores in the membrane at the same time (ensemble average) (Figure 2c) results in the same response as the time average performed in Figure 2b. Contrary to previous interpretations^35^, this proves that gating is ergodic. Furthermore, using phenomenological modelling, we were able to capture the transition from single pore to collective behaviour. The conductance of the ensemble membrane can be expressed from single pore gating dynamics through the following equation:

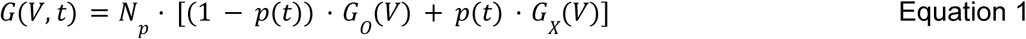

where *N_p_* is the number of pores, *p(t)* is the time-dependent probability for the pores to be closed and *G_O/X_(V)* refers to the single-pore voltage-dependent conductance in the open/closed state, respectively. Ensemble measurement *IV* curves were normalised by the single pore *IV* curve in a voltage window where no gating takes place which yields *N_p_* (details in SI Section 4). We could reliably conduct single-pore measurements where the pore remains open for a full cycle (Figure 2b), yielding *I_O_ = G_O_(V(t))*·*V(t)*. Thus, relying on the observation from single-pore measurements that the gated state conducts 14% of the open-pore current on average (*G_x_* ≃ 0.14 · *G_O_*, Figure 1d) we can invert Equation 1 to obtain *p(t)*. The measured probability *p(t)* is governed by the following rate equation:

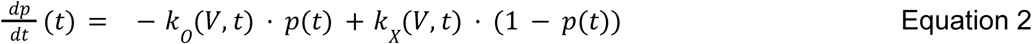

By using constant voltage pulses, similar to the methodology used by Rappaport et al.^36^, we were able to extract the opening rate *k_O_* and closing rate *k_X_*at different voltage biases and accurately fit the ensemble memristive hysteresis across various frequencies (SI Figure 4).

**Figure 4.**
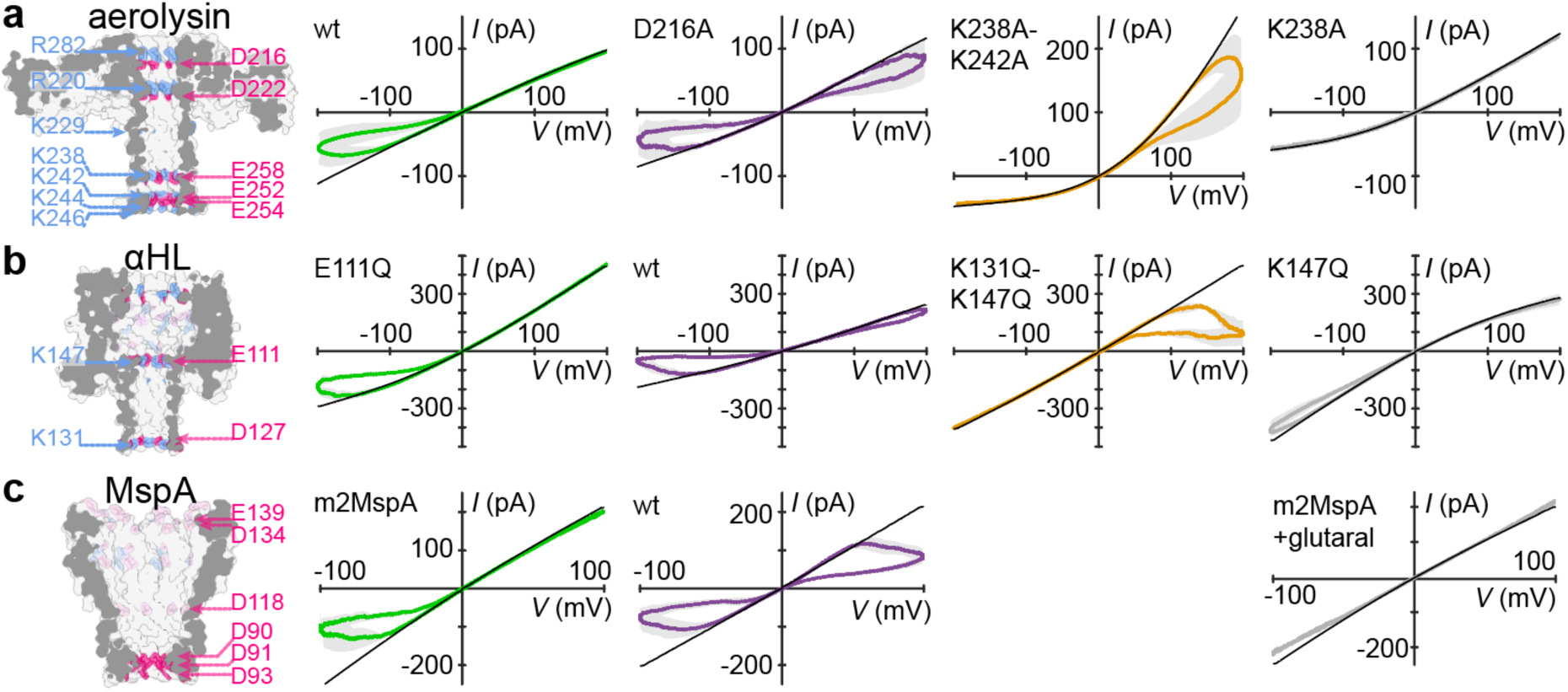
Gating behaviour measured with different aerolysin (a), α-HL (b) and MspA (c) mutants, showcasing the great tunability of gating by changing lumen charge. Mutants gating at negative potentials displayed in green, at positive potentials in orange, mutants showing low levels of gating in grey and mutants that gate at both polarities in purple. Each plot displays the open-pore *IV* curve in black, displaying the differences in rectification.

Following this analysis, we combine ensemble and single-pore measurements in order to quantify gating in various different experimental conditions. SI Figure 4 shows that when the pores start closing above a critical voltage *V_c_*, the opening rate is zero. This means that for *|V| > V_c_* the first term in equation 2 is dropped. Therefore the maximum slope of *p(t)* is simply the rate of closing *k_X_* (Figure 2c, inset), that we chose to quantify gating.

Overall, ensemble measurements present three major advantages over single pore measurements. They offer a high throughput and a large signal to noise ratio, and have a higher success rate since it is much easier to incorporate many pores in a membrane than one. Thus, relying on the ergodicity of the stochastic ion transport through aerolysin pores, ensemble measurements enable the high-throughput quantification of gating. Our measurements inspired by the memristor literature also allow us to quantify both open-pore rectification and gating independently and reliably by varying the voltage frequency. We now harness this technique to understand the role of pore lumen charge on ion transport in biological pores. To achieve this, we tuned the lumen charge at specific locations inside the different pores by creating a wide mutant library. We focus mainly on aerolysin because of its constant lumen radius even though our conclusion may be extended to other pores with variable radii.

### Open-pore rectification of aerolysin is directly driven by lumen charge

Mutations allow us to easily add, delete or change lumen charges at a resolution equal to the multimericity of the pore. In the case of the heptameric aerolysin, when we mutate for instance a negative amino acid to a neutral one, the resulting assembly will have lost seven negative charges compared to the wild type (wt) pore. Aerolysin wt has 13 charged amino acid positions in the β-barrel region, thus corresponding to 13 rings of seven charged amino acids at given z-positions along the β-barrel (Figure 3a). Two of these rings have amino acids pointing outwards towards the electrolyte and membrane (K229 and E237, although E237 was revealed by cryo-EM to be in a protonated form^37^), leaving 11 charged rings with amino acids pointing inside the pore lumen. The upper 4 charged positions come in clear pairs where a negative ring is right above or below a positive one (R282 and D216; R220 and D222), and create the first known constriction of the pore^14,38,39^. The remaining 7 charged positions are located in the transmembrane region of the β-barrel: K238 and E258 are also right next to each other, followed by K242, E254, E252, K244 and K246 (Figure 3a), which form the second sensing constriction leaving the pore with an overall positive theoretical charge.

We mutated different combinations of these charged amino acids to change distribution of the pore lumen and thus measured a library of 26 different aerolysin mutants (SI Figure 5). In what follows, we first focus on the consequences of mutations over the open-pore ionic transport by using high frequency sweeps at low pore numbers. We then study the effect of mutations on the gating behaviour. We report in Figure 3b, *IV* curves measured with high frequency sweeps for all our aerolysin mutants (separated curves can be found in SI Figure 5). A first observation is that modifying lumen charges has dramatic consequences over the shape of the open-pore *IV* curve. Convex and concave rectification can be observed, as well as linear behaviour. Further, we find that the conductance of all mutants is below the bulk theoretical conductance of 2 nS by a factor of 2-8 (SI Table 1). We attribute the reduced and nonlinear ionic conduction to the presence of energy barriers arising from the combination of lumen charge and nanometric confinement. Altogether, these results show that ion transport through aerolysin, like α-HL^12,40^ is controlled by lumen charges and highly sensitive to its modifications which are easily achieved via well-established bioengineering processes.

**Figure 5.**
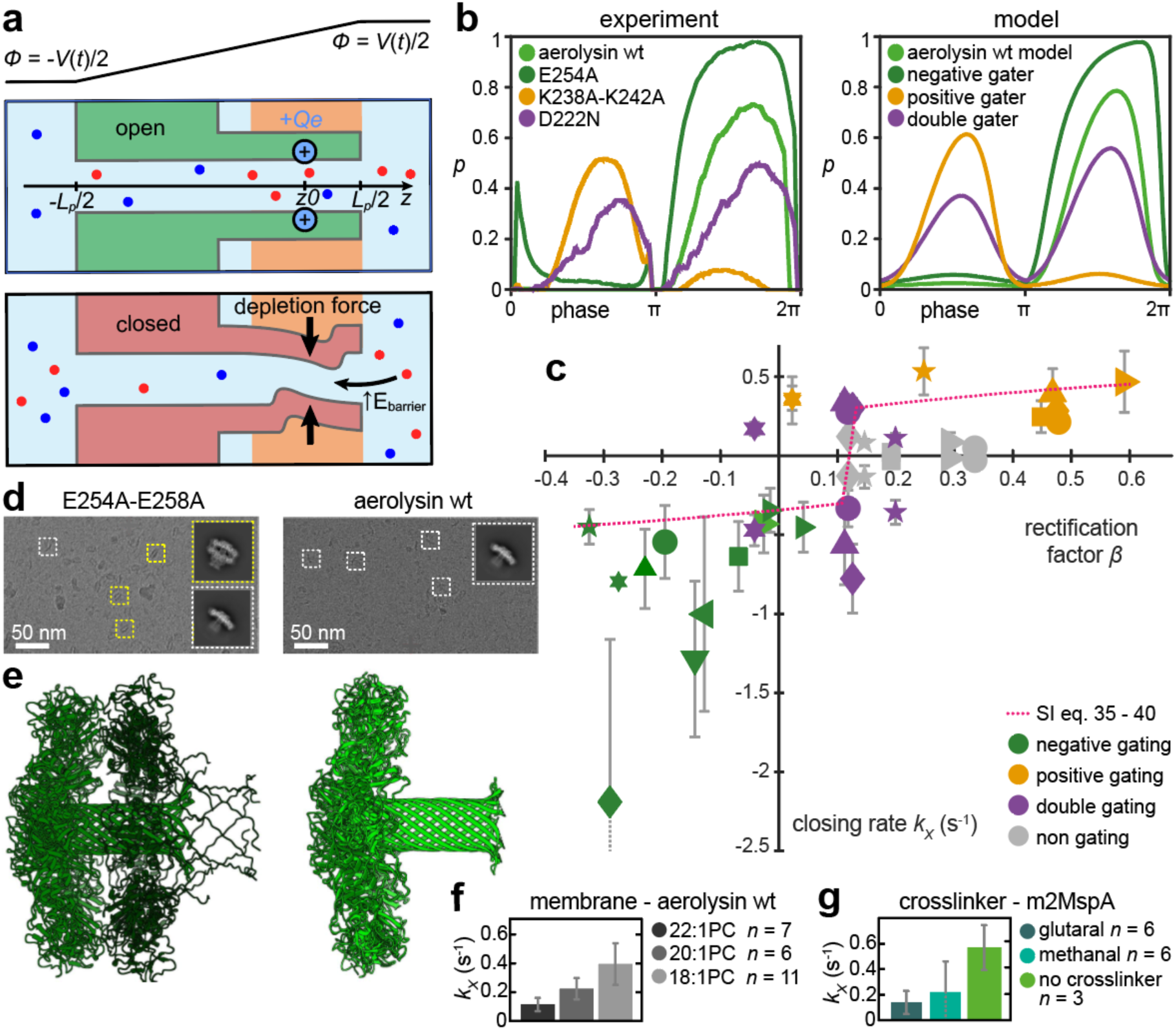
Evidence in support of a structural deformation. a. Sketch illustrating both states of the model. The pore shows mechanical bistability induced by depletion forces: when the pore diameter fluctuates below a certain threshold, ions are expelled from the pore, which partially collapses. This effect is promoted by dissociation of counterions from lumen charges under an applied voltage. **b.** Closed-state probability plotted against the phase of the voltage for aerolysin wt and four representative mutants obtained from experiments **(left)** and biophysical model **(right)** (see SI Section 2 for details). The model accurately predicts if the given mutant is a positive, a negative or a double gater. It also predicts that the E254A mutant gates stronger than aerolysin wt. **c.** Closing rates of aerolysin wt for different membrane thickness and of m2MspA and m2MspA cross-linked with glutaral and methanal (1 M KCl, 10 mM phosphate, pH 6.2, 0.1 Hz). **d.** Scatter plot of the pH response and all 26 aerolysin mutants’ closing rates *versus* rectification factor *β*. Pores that show low gating or gating at both polarities are represented twice. For pores that show gating only in the negative or positive quadrant only one value is shown. The error bars show the standard deviation of closing rates of different repeat experiments. Each symbol represents a different mutant, see SI Figure 12. Dotted line: theoretical prediction based on models of rectification and gating (SI Section 2).

The effect of ionic rectification, also known as nanofluidic diode, has been the study of many experimental and theoretical studies in biological channels^41^ as well as solid-state nanofluidics^42–44^. However, while recent advances have enabled the fabrication of sub-nanometer pores in 2D materials and angstrom slits^45^, solid state nanofluidics has predominantly focused on relatively large channels (∼10-100 nm in diameter) with spatially-extended surface charges and geometrical asymmetries while smaller biological nanopores are explored mainly through simulations. Here, we developed a theoretical framework to tackle the effect of individual, localised charge in a biological pore with sub-nanometric radius. To rationalise this, we derived an analytical theory (detailed in SI Section 1) accounting for rectification and linking its magnitude to the distribution of charge in the pore. Briefly, we modelled aerolysin as a cylindrical pore with a distribution of surface charges *ρ* corresponding to charged residues (Figure 3a). These charges define the potential energy landscape experienced by diffusing ions. When an electric field is applied, ions accumulate downstream of potential wells and upstream of potential barriers, and are depleted in opposite configurations (Figure 3a and SI Figure 18). Depending on the distribution of lumen charges, which act as potential wells for counterions, this effect leads to an overall increase or decrease in the number of ions inside the pore, and likewise in conductance. We observe an excellent agreement between the model and experimental data (Figure 3c). In particular, our model correctly predicts that the charged groups located near, but not at the mouth of the pore play a key role in ionic conduction. This is because a charge that is placed at the mouth of the pore, will modify counterion distributions mostly in the reservoir, rather than inside of the pore.

When removing all above-mentioned charged amino acids present in aerolysin wt, the resulting *IV* curve is almost perfectly ohmic (SI Figure 6). We can thus deterministically control the degree of rectification as well as completely suppressing it, relying on the accurate mutation. We also experimentally confirmed that variations in ionic transport are mainly controlled by charge modification rather than a change in pore diameter by observing that mutating lysines (K) (168.6 Å^3^) to alanine (A) (88.6 Å^3^), serine (S) (89.0 Å^3^) or the much larger glutamine (Q) (143.8 Å^3^)^46^ is nearly equivalent (SI Figure 5). Having fully rationalised the relationship between lumen charge and open-pore ionic conduction, we now study the influence of lumen charges and their modification on the gating.

**Figure 6.**
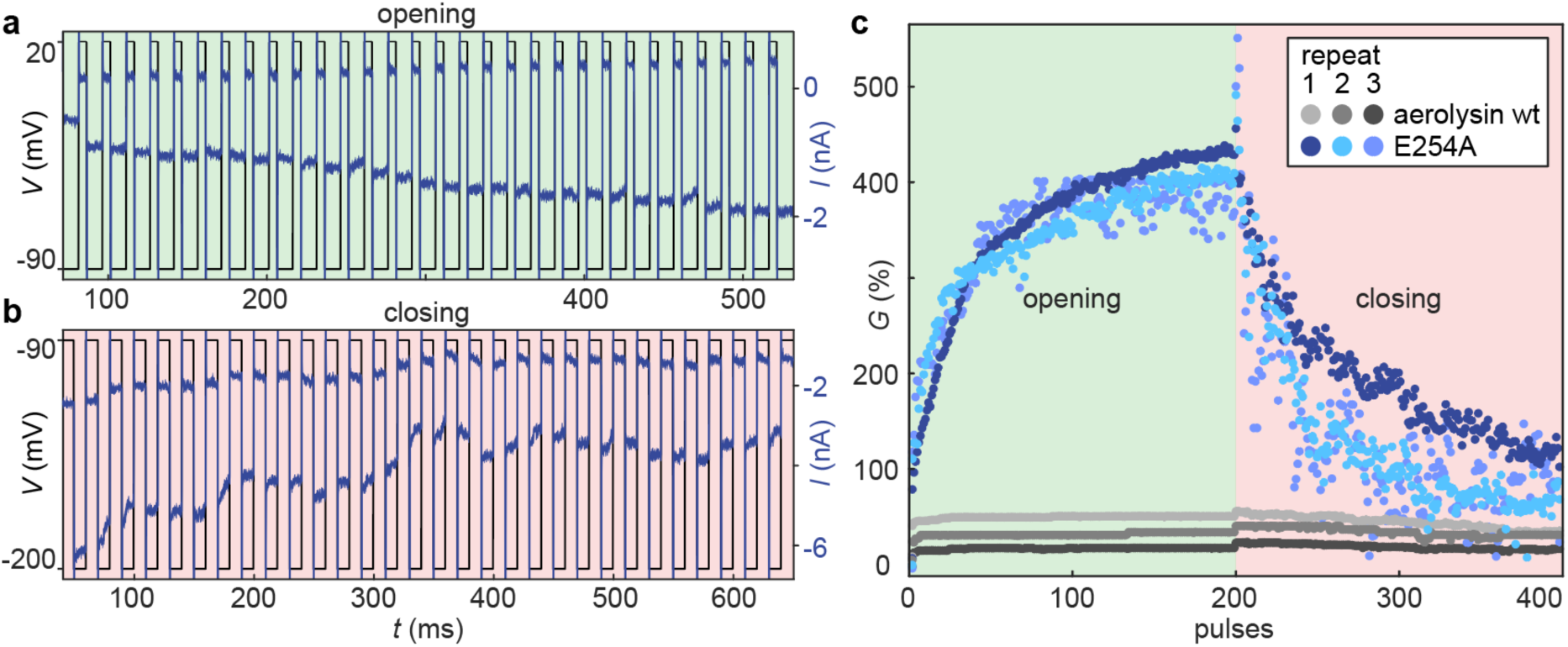
Synaptic dynamics with biological nanopores. **a.** Synaptic potentiation, through repeated voltage pulses, is used to open the ensemble of nanopores in the membrane. 110 mV pulses with duration of 5 ms are used. **b.** Synaptic depression through repeated voltage pulses, used to close the ensemble of nanopores in the membrane.-110 mV pulses with a duration of 10 ms are used. For both **a** and **b,** the waiting time between pulses was 10 ms and the first 30 pulses are shown. The baseline was fixed at-90 mV in order to minimize closing or opening of the pores. **c.** Conductance evolution during potentiation and depression. The conductance normalised by the closed state current is shown for three repeat experiments of aerolysin wt and E254A. The overshooting when switching from opening to closing is due to the rectification.

Further, each plot shows the standard deviation and number of experiments as a grey area. All experiments were conducted in 1 M KCl buffered to pH 6.2 with 10 mM phosphate. Sinusoidal forcing was applied at 0.1 Hz and 200 mV amplitude (100 mV for MspA wt and m2MspA).

### Gating behavior is controlled by pore charge

We now examine the gating behaviour of our mutant library. We display in Figure 4a-c the *IV* curve obtained for several mutants of aerolysin, α-HL and MspA using low-frequency sinusoidal sweeps. We can observe that, depending on the mutations, gating occurs at positive, negative or both polarities, or is largely absent. Gating appears to be additive: for example, aerolysin wt shows negative gating, the two mutations K238A and K242A individually lead to a suppression of the gating and collectively (K238A-K242A) change the gating all the way to positive gating. This finding suggests that the underlying gating mechanism remains the same for all mutants, and that the lumen charges individually contribute to the stability of the closed state under a given voltage, tuning the kinetics (e.g. closing rate) of gating. Taken together with previous observations^3,25,47–49^, this demonstrates both the universality of this phenomenon in β-barrel forming nanopores and its sensitivity to local charge variations. To further back the phenomenon’s universality, we confirmed the previously observed increase in gating with decreasing charge carrier concentration^19,50^, pH^18,50–52^, and temperature^53,54^ (SI Figure 15). As for the rectification, we verified that gating is dependent on the charge but not on the size or hydrophilicity of the exchanged amino acids. Indeed, the gating behaviours of K238A-K242A, K238S-K242S and K238Q-K242Q as well as D216A and D216N are highly similar (SI Figure 9, SI Table 1). Since A, S, Q and N are all non charged but differ in size and hydrophobicity, this leads us to conclude that the change of charge is the determining factor of gating. Gating, like rectification, is thus highly sensitive to the charge distribution in the pore lumen. Since both phenomena share a common dependency, we explore their correlation.

For aerolysin, we observed a correlation between closing rate, obtained from low frequency sweeps and rectification factor *β*, obtained from high frequency sweeps (Figure 5c). This means that the gating is most likely to occur at the polarity where the open-pore current is highest as can be observed in Figure 1c and Figure 5c. The correlation between the closing rate and *β*, which our rectification model previously showed to be solely dependent on lumen charge, clearly indicates the central role of lumen charge in the gating phenomenon.

While tuning of the lumen charge affects gating behaviour very predictably in aerolysin, α-HL and MspA (Figure 4a-c), the aerolysin mutant with a neutral lumen exhibits polarity-dependent gating (SI Figure 9). This strongly indicates that additional charges such as the ones placed outside of the lumen (e.g. K229) also play a role in gating. A non charge-related mechanism is not likely precisely because of the polarity dependence of the gating of the neutral lumen, and a neutral lumen is likely because of the ohmic conductance of that mutant. Overall, this suggests that the pore exhibits some mechanical bistability even without lumen charges, but the latter strongly controls this effect.

### Gating can be explained by a voltage-induced conformational change

A reversible change of lumen conformation, triggered by voltage and controlled by lumen charge as well as mechanical properties has been previously hypothesised for the studied pore-forming proteins^20,22^ and was demonstrated for structurally different voltage-gated ion channels^10,55^. In some cases, gating was attributed to the motion of flexible and charged elements within the pore^49^; however gating could still be observed when these groups are deleted^56^. This suggests that gating could emerge from the structure and dynamics of the β-barrel itself. While obtaining direct experimental proof of a conformational change at these length scales is challenging, our dataset combines several clues pointing in that direction.

Based on all experimental findings thus far, we build an analytical biophysical model for gating in aerolysin, detailed in SI Section 2. The core idea of the model is that the β-barrel is partially flexible, allowing for some fluctuations in the pore’s radius. However, ions face an electrostatic energy barrier to enter confined systems: a radius variation of around 30% leads to an increase of this energy barrier by around 5 *k_B_T*. Then, it becomes energetically favourable to expel ions from the pore by partially collapsing its structure (Figure 5a). This process, known as depletion interactions in colloid science, induces an inherent bistability of the pore between a conductive open state filled with ions and a resistive depleted closed state. Indicated by the diversity of residual current (Figure 1d), the collapse of the β-barrel is likely structurally diverse in that the barrel may deform at different locations to different degrees.

We then ask how an external voltage might favour depletion interactions, and how the effect of voltage polarity and lumen charges could come into play. Counterintuitively, external electric fields are not expected to exert any net mechanical force on fixed lumen charges; this is because the electrostatic force 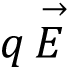 is cancelled out by electro-osmotic forces (originating from the same electric field acting on counterions, inducing a water flow and a friction force 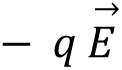 on the lumen walls). However, we find that the extreme confinement in the closed state can induce a partial breakdown of electroneutrality: briefly, an external field can induce a separation of a lumen charge/counterion pair. In that case, electro-osmosis will no longer compensate electrostatic forces.

Quantitatively, we build a statistical mechanics toy model for the process outlined above. We find that the pore oscillates between two states, one open and one partially collapsed, due to depletion forces. When an external field is applied, the energy of the open state is unchanged, but the closed state is stabilized by the charge separation effect depending on the position of the lumen charges and the voltage polarity, promoting gating. This model recapitulates all the key features of gating. In particular, both in the model and in experiments, the open state is maximally favoured at zero voltage, when charge separation is least likely to occur. While gating is maximal for a voltage of a certain polarity, a weak gating effect still exists under a strong enough voltage of the opposite polarity. Overall, we model each aerolysin pore (wt or mutant) by an effective lumen charge placed at some position *z_0_* along the pore (Figure 5a). Doing so, we accurately reproduce the observed dependency of gating with charge distribution (Figure 5b). The model also accounts for the effect of ionic strength — a higher concentration leads to stronger energy barrier between the open and the closed states due to depletion forces, reducing the gating rate (SI Figure 2c and 15); and of temperature — a higher temperature speeds up the entire dynamics, leading to a greater gating rate (SI Figure 12k and 15). Our model does not account for the effect of ionic species. However, the proposed mechanism is reminiscent of Coulomb blockade, which is known to depend on e.g. ion valence^57^. Similar effects could explain the phenomenology observed here (SI Figure 15).

Lastly, combining our models of rectification and gating, we are able to rationalize the correlation observed between these two effects. Both effects are stronger when the electric field is directed from lumen charges towards the closest entrance of the pore, therefore dragging counterions over a substantial portion of the channel. Plotting the theoretical closing rate *k_x_* versus the rectification factor *β*, we find good overall agreement with experimental data (Figure 5c). The graph features a vertical line, where gating and rectification seem to decouple: this corresponds for the most part to mutants with no net lumen charge (exhibiting only pairs of opposite lumen charges, which contribute to rectification but not to gating), and mutants with approximately symmetrical lumen charges of the same sign (causing gating in both polarities, but contributing only weakly to rectification). Overall, these results highlight how charge location can affect non-linear ion transport in biological nanopores across both fast and slow time scales.

We also perform several control experiments corroborating the hypothesis of voltage-induced structural deformations. When comparing cryo-EM structures of aerolysin wt and the strongly gating E254A-E258A mutant, we find that around 44 % of the E254A-E258A particles appear in dimeric pore stacks formed by one mature aerolysin pore within an aerolysin pre-pore (Figure 5e), similar to what previously found with a disulfide-cross-linked K246C-E258C mutant^58^. This indicates that, while aerolysin wt univocally forms a mature pore β-barrel in multiple membrane-mimicking environments^39^, the E254A-E258A mutant shows a much lower efficiency for β-barrel formation as roughly half of the particles observed are trapped in the pre-pore conformation where the β-barrel is not yet formed. This shows that these mutations impact mechanical stability of the pore. Another hint is the dependency of the gating with membrane thickness (Figure 5f & SI Figure 15). Membrane thickness and composition is known to influence the dynamics of membrane proteins^59^, and has been implicated in gating^60,61^. By changing the lipid tail length, we find that gating decreases with increasing membrane thickness. The change in thickness leads to a change in lipid protein contact area which leads to a shift in the forces acting on the protein through a difference in hydrophobic mismatch, pressure and tension, as well as potential steric hindrance with the above-membrane parts of the nanopore. The observation that gating is membrane-dependent points to the transmembrane region as the active site of the mechanism. To further verify the hypothesis of a conformational change, we cross-linked the highly-gating m2MspA pore with glutaral or methanal in order to increase its mechanical stability (Figure 5g & SI Figure 15). We find that such cross-linked pores do not gate while maintaining the same rectification and conductance (SI Table 2) as the untreated m2MspA.

Mutating charged residues in the pore lumen not only alters the electrostatic environment but also disrupts the hydrogen-bond network, salt bridges, and other intermolecular forces, leading to changes in the mechanical stability of the pore^18^. This intricate interplay between lumen charge and mechanical deformation within nanometric confinement helps explain why characterizing and molecularly understanding gating remains so challenging. Nevertheless, the overwhelming uniformity with which these β-barrel nanopores respond to voltage, frequency, temperature, ionic strength, pH, and lumen charge distribution strongly indicates a common underlying mechanism. As proposed previously^11^, this deformation is likely localized in the transmembrane region shared by all β-barrel pore-forming toxins. By combining time-varying forcing with a wide library of mutants, cross-linked pores, and membrane modifications, we achieved a comprehensive understanding of this mechanism. Importantly, while our work focused on multimeric β-barrel nanopores, all comparable results reported for monomeric β-barrel channels are consistent with our observations, supporting the conclusion that this phenomenon extends across both classes. More broadly, we expect it to be generally applicable to any pore with a radius around 1 nm, although other pore-specific mechanisms may also contribute to gating.

### Synaptic plasticity emulated with engineered biological nanopores

To demonstrate the implications of our findings in the context of ionic bio-computing, we implement a synaptic behavior using a carefully selected mutant of aerolysin (E254A). This mutant is chosen for its strong gating rate and on/off ratio, in agreement with our phenomenological model (SI Figure 8 and Figure 5b). The membrane’s conductance can be increased sequentially (potentiation) upon the application of positive voltage pulses (Figure 6a). The pores are thus “learning” upon repeated simulation like a biological synapse. By applying negative pulses, the conductance can be progressively rested (depression) to its initial level (Figure 6b). For each pulse, the conductance (defined as *G = ΔI/ΔV*) can be extracted. By plotting its relative variation in Figure 6c, we show how our membrane embedded biological nanopores can be programmed as volatile memory. Here the conductance plays the role of an analog variable that can be set and reset using voltage pulse stimuli. Our biological nanopore synapses exhibit an on/off ratio of 4 in pulse operation and can be programmed with short pulses of a few milliseconds. When using sinusoidal forcing of 0.3 Hz at an amplitude of 150 mV we were able to run two experiments for 5600 cycles (SI Figure 15). These performances are significantly better than several solid-state nanofluidic memristors^62–64^ which typically have at least one dimension on the micron scale.

To highlight the impact of the mutation on the observed memristive performances we report the synaptic dynamic of the aerolysin wt, which exhibits a much weaker effect (Figure 6c). By relying on protein mutation, it is thus possible to engineer nanofluidic synapses with tailored plasticity. This versatility is, to the best of our knowledge, a unique property of biological pores, and could be harnessed for designing nanofluidic neural networks with deterministic node properties.

## Conclusion and Outlook

β-barrel-forming nanopores, whether toxins or transporters, fulfill a variety of biological roles, yet they all mediate the passage of ions, water, and small solutes across membranes. A central feature of these pores is their ability to regulate molecular transport in response to voltage, a fundamental mechanism in many living systems. In this work, we demonstrate that key aspects of biological ion transport can be rationalized using a nanopore characterization approach based on alternating current, offering new insights into voltage-dependent transport processes. We show that the heterogeneous lumen charge controls two distinct nonlinearities: open-pore rectification and gating. Our work comprises a wide database of mutants elucidating this phenomenology beyond what has been shown in numerous previous studies ^11,20,22^. In light of these findings, we built detailed biophysical models of rectification and gating. Notably, we propose that gating results from ionic depletion interactions that lead to a partial deformation of the β-barrel. Then, we show that the collapsed state can be stabilized by the external voltage through a dissociation of lumen charge-counterion pairs. Overall, we provide a plausible and detailed explanation that accounts for the observed nanopore behavior across a broad range of experimental conditions and control settings. Direct experimental evidence for the physical origin of this extensively studied phenomenon remains challenging to obtain. Nonetheless, the conformational dynamics of the β-barrel we uncovered calls for confirmation using techniques such as single-molecule FRET, and our theoretical developments could be supplemented with enhanced sampling approaches to simulate the pore dynamics. Furthermore, our approach could be extended to the numerous different biological nanopores in order to deepen our understanding of Nature’s sophisticated ionic machinery.

Understanding nanopore physics is irremissible to tap into their full power, both for emerging techniques like protein sequencing as well as computing applications with diverging strategies. On the one hand, designing gating-free mutants as well as *de novo* designed nanopores^65^ tailored for sensing applications will remove limitations for this research field and nascent industry. On the other hand, rectification and gating can be employed to generate iontronic components like resistors, diodes or memristors with a tuning resolution reaching the physical limit of 1 single charge. While the construction of multi-component ionic computing systems is still at its infancy^33^, our synaptic plasticity results show that such networks could potentially use biological building blocks with dimensions similar to state-of-the-art electronic transistors.

## Acknowledgements

We are grateful to Dr. Michael Mayer and Dr. Gisou van der Goot for their insightful discussions and thoughtful feedback. We acknowledge funding from the European Research Council (grants 101020445—2D-LIQUID N.R. and A.R., MSCA No. 101034413 P.R.), the Swiss National Science Foundation (grants 205321_192371, and 200021L_212128 to M.D.P., TMPFP2-217134 to T.E, and IZSEZ0_183779 to J. H. G. and A. R.) and the Swiss National Supercomputing Centre (CSCS) for access to the HPC resources used to run MD simulations. We thank the staff members of the Dubochet Center for Imaging in Lausanne, in particular Dr. Emiko Uchikawa and Dr. Sergey Nazarov, for their assistance with cryo-EM sample preparation and data collection. We thank Prof. Aleksander Antanasijevic and Dr. Yoan Duhoo from EPFL Protein Production and Structure Core Facility for their support in cryo-EM data processing.

## Author contributions

S.F. Mayer, T. Emmerich, N. Ronceray, M. Dal Peraro and A. Radenovic conceived the project. S.F. Mayer and M.F. Mitsioni have conducted and analysed planar lipid bilayer experiments with help of S. Soussou, J. Jeanneret-Grosjean, L. van den Heuvel and A. Fulciniti. L. van den Heuvel has written the data processing code and processed the simulation data under the supervision of S.F. Mayer. P. Robin has built the models for rectification and gating. L.A. Abriata has run the simulations and helped with their processing. L.F. Krapp has analysed the single pore aerolysin gating data and simulation data. M.F. Mitsioni, with the help of S. Vacle, L. Feletti and A. Möller, produced nanopores under the supervision of M.J. Marcaida. J.S. Anton has done all cryo-EM work and conducted the blood assay. H.D. Brinkerhoff, A.H. Laszlo and J.H. Gundlach provided initial guidance setting up DC measurements and provided feedback on the project. S.F. Mayer, T. Emmerich, P. Robin, N. Ronceray and M.F. Mitsioni wrote the manuscript with the input from all authors. A. Radenovic and M. Dal Peraro jointly supervised this work.

## Competing interests

The authors declare that they have no conflict of interest.

## Methods

### Section 1. Material

We purchased 1,2-diphytanoyl-sn-glycero-3-phosphocholine (DPhPC), 1,2-di-O-phytanyl-sn-glycero-3-phosphocholine (DoPhPC), 1,2-dimyristoleoyl-sn-glycero-3-phosphocholine (14:1_Δ9-Cis_PC), 1,2-dipalmitoleoyl-sn-glycero-3-phosphocholine (16:1_Δ9-Cis_PC), 1,2-dioleoyl-sn-glycero-3-phosphocholine (18:1_Δ9-Cis_PC), 1,2-dieicosenoyl-sn-glycero-3-phosphocholine (20:1_cis_PC), 1,2-dierucoyl-sn-glycero-3-phosphocholine (22:1_cis_PC) dissolved in chloroform from Avanti Polar Lipids (Birmingham, AL, USA), n-Octyltetraoxyethylene (OTOE) from Bachem (Bubendorf, CH), Plasmids from Genscript (Piscataway, NJ, USA), Turbonuclease from Lucerna-Chem AG (Luzern, CH), HEPES from Chemie Brunschwig AG (Basel, CH), 0.45 µm Rotilabo PVDF syringe filter from Carl Roth GmbH & Co. KG (Karlsruhe, DE), isopropyl β-d-1-thiogalactopyranoside (IPTG) from Huberlab (Aesch, CH), defibrinated sheep blood from Rockland Immunochemicals, Inc. (Limerick, PA, USA), HiPrep 26/10 Desalting column, HisTrap HP column and PD-10 desalting columns packed with Sephadex G-25 resin from Cytiva (Marlborough, MA, USA), and all other chemicals from Merck (Darmstadt, DE). All electrolyte solutions used in bilayer experiments were buffered with 10 mM phosphate and filtered through 0.22 µm pore size PES vacuum filters from VWR International (Radnor, PA, USA). We used purified water (18.2 MΩ) from a milli-Q water purifying system. Ag/AgCl wire electrodes were made by immersing one side of high purity silver wire pieces in bleach for 1 h and subsequent washing with water. Lipids were aliquoted at 0.2 mg per glass vial, chloroform was evaporated under vacuum and vials subsequently closed with Polytetrafluoroethylene (PTFE) lined caps, in an argon glove box and stored at-20 °C to avoid oxidation. Lipids were dissolved freshly in octane or pentane at 10 mg/mL.

### Aerolysin purification

The clone of the full-length aerolysin wt protein (in the pET22b vector with a C-terminal hexa-histidine-tag (His-tag)) was kindly provided by the Van der Goot laboratory at EPFL. The mutants were generated by Genscript using the wt plasmid as a template. The aerolysin protein was produced as previously described by Cao et al.^14^. In brief, plasmids were transformed into BL21 (DE3 Plys) *E. coli* cells by heat shock. Cells were grown to an optical density of 0.6 to 0.7 at 600 nm in LB media. Protein expression was induced by the addition of 1 mM IPTG and subsequent growth overnight at 18 °C. Cell pellets were resuspended in aerolysin-lysis-buffer (500 mM NaCl, 50 mM tris, pH 7,4), mixed with cOmplete Protease Inhibitor Cocktail, and then lysed by sonication on ice. Turbonuclease solution was added to the resulting suspension which was then centrifuged (20’000 *g* for 30 minutes at 4 °C). After syringe filtration over a 0.45 µm filter, the supernatant was applied to a HisTrap HP column previously equilibrated with aerolysin-lysis-buffer. The protein was eluted as a monomer with a gradient of 30 column volumes of aerolysin-elution-buffer (500 mM NaCl, 50 mM tris, 500 mM imidazole, pH 7,4). aerolysin-containing fractions were then buffer-exchanged to aerolysin-final-buffer (20 mM tris, 150 mM NaCl, pH 7.4) using a HiPrep Desalting column. The purification of the selected fractions was confirmed by SDS-polyacrylamide gel electrophoresis. Aerolysin was concentrated to 0.5 mg/mL and stored at-20 °C. The activation of aerolysin to undergo conformational changes and enable heptamerization is achieved by incubating it with trypsin in agarose at a ratio of 1:4 (trypsin:pore, *v/v*).

### α-HL purification

The coding sequences for all α-HL mutants were custom synthesised, optimised for *E. coli* expression and inserted into the pET28-b vector between the NcoI and XhoI cleavage sites, such that the resulting proteins carry a C-terminal His-tag for later purification. Plasmids were transformed in BL21 (DE3) cells by heat shock transformation. The cells were grown in LB medium at 37 °C until optical density of 0.6 to 0.7 at 600 nm at which point protein expression was induced by the addition of 0.1 mM IPTG, followed by subsequent overnight growth at 18 °C. Cell pellets were resuspended in α-HL-lysis-buffer (500 mM NaCl, 20 mM HEPES, 1% Triton X-100 (*v/v)*, pH 8.0), mixed with cOmplete Protease Inhibitor Cocktail, and then lysed by using sonication on ice. Turbonuclease solution was added to the resulting suspension which was then centrifuged (20’000 *g* for 60 minutes at 4 °C). After syringe filtration over a 0.45 µm filter, the supernatant was applied to a HisTrap HP column previously equilibrated with α-HL-lysis-buffer. The column was then washed with 3 column volumes of α-HL-lysis-buffer followed by a wash with 10 column volumes of α-HL-wash-buffer (500 mM NaCl, 20 mM HEPES, 5 mM imidazole, pH 8.0). The α-HL heptamers were then eluted with a linear gradient of α-HL-elution-buffer (500 mM NaCl, 20 mM HEPES, 500 mM imidazole, pH 8.0). α-HL pores were not buffer-exchanged to a final buffer but rather stored in the ratio of α-HL-wash-buffer and α-HL-elution-buffer they eluted in. Peak fractions were analysed by SDS-polyacrylamide gel electrophoresis and only the fractions with the α-HL heptamer were retained for subsequent planar lipid bilayer experiments.

### MspA purification

The coding sequences for all MspA mutants was custom synthesised, optimised for *E. coli* expression and inserted into the pET28b vector, between the NcoI and XhoI cleavage sites, such that the resulting proteins have a C-terminal His-tag to aid purification. Plasmids were transformed in *E. coli* BL21 (DE3) cells. Protein expression was induced by the addition of 0.1 mM IPTG as soon as the cells reached an optical density of 0.6 and subsequent growth overnight at 18 °C. Cell pellets were resuspended in MspA-lysis-buffer (100 mM sodium phosphate, 0.1 mM EDTA, 150 mM NaCl, 0.5% (*w/v*) Genapol X-80, pH 6.5), mixed with cOmplete Protease Inhibitor Cocktail, and then lysed using sonication on ice, followed by boiling at 60 °C for 10 minutes. Turbonuclease solution was added to the resulting suspension which was then chilled in ice for 10 minutes. To remove intact cells and cell debris, the samples were centrifuged at 20’000 *g* for 60 minutes at 4 °C. After syringe filtration over a 0.45 µm filter, the supernatant was applied to a HisTrap HP column previously equilibrated with MspA-lysis-buffer. The column was then washed with 3 column volumes of MspA-lysis-buffer, followed by 10 column volumes of MspA-wash-buffer (500 mM NaCl, 20 mM HEPES, 5 mM imidazole, 0.5% (*w/v*) Genapol X-80, pH 8.0). The MspA octamers were then eluted with a linear gradient of MspA-elution-buffer (500 mM imidazole, 500 mM NaCl, 20 mM HEPES, 0.5% (*w/v*) Genapol X-80, pH 8.0). Peak fractions were analysed by SDS-polyacrylamide gel electrophoresis and only the fractions with the MspA octamer were retained. The protein was then buffer-exchanged to MspA-final-buffer (500 mM NaCl, 20 mM HEPES, pH 8.0, 0.6% OTOE) using PD-10 desalting columns and subsequently tested in planar lipid bilayer experiments.

The cross-linked pores were prepared as follows. 100 μL of MspA wt or m2MspA were mixed with 10 μL of 8% (*v/v*) glutaral in water to a final concentration of 0.8% glutaral and incubated for 10 minutes before being added to the experiments. For pores cross-linked with methanal, 100 μL of pore solution were mixed with 5 μL of 20% (*v/v*) methanal in water to a final concentration of 1% (*v/v*) methanal and incubated for 10 minutes before addition to the planar lipid bilayer experiment.

### Section 2. Experiments

#### Planar lipid bilayer experiments

Unless stated otherwise, an Orbit Mini (Nanion Technologies, Munich, DE) with 50 µm aperture MECA-4 chips (Ionera Technologies GmbH, Freiburg, DE) was used for all experiments. Bilayers were painted by dipping, pipette tips in 10 mg/mL DPhPC in octane, and subsequent dabbing on Kimwipes, so that all visible liquid was removed from the pipette tip. The pipette tip was then immersed in the electrolyte solution filled MECA-4 chip, placed over one of the four apertures and a bubble was pushed in and out of the pipette tip until a membrane was formed. This usually required several tries, exchanging lipid-primed pipette tips frequently. For all open-pore *IV* curve recordings, we pretreated PTFE films (Eastern Scientific LLC, Rockville, MD, USA) with a single aperture with a diameter of 50 μm by pipetting 1 μL of 1% (*v/v*) hexadecane in hexane onto both sides of the aperture of the PTFE film. We mounted the PTFE film in a PTFE chamber using high-vacuum grease (Dow Corning Corporation, Midland, MI, USA). The PTFE film separated two compartments containing aqueous electrolyte, which were only connected by the aperture in the PTFE film over which the bilayer was formed. We formed planar lipid bilayers across the aperture using a technique previously described^26,66^. Briefly, we added electrolyte solution to both compartments so that the level was below the aperture and pipetted 3 μL of DPhPC dissolved in pentane onto the surface of the electrolyte solution. After the pentane evaporated, a lipid monolayer formed at the air-aqueous interface. We raised and lowered the electrolyte solution until we measured a very small baseline current (-3 pA < *I* < 3 pA) indicating that a membrane had formed. All single pore DC experiments and the experiments with sine wave frequencies at and below 0.001 Hz were measured in painted bilayers formed on selfmade PTFE/FEP supports made from Dual-Shrink tube (Zeus, Orangeburg, SC, USA) using a tungsten needle, sharpened from a welding electrode. Supports were pretreated by pipetting 1 µL DoPhPC in hexane (2 mg/mL) onto the support, blowing through the support from the backside with an air filled syringe to clear it from the lipid solution. This pretreatment was repeated 2 times with short drying cycles in a desiccator under vacuum. The bilayer was painted from pure DoPhPC wetted with hexadecane with a two-haired fine paintbrush. We verified the stability (absence of leak currents; expected noise & capacitance level) of all bilayers by briefly applying transmembrane voltages of up to 200 mV. The current was measured with an axopatch 500B amplifier (Molecular Devices, LLC.,San Jose, CA, USA) through Ag/AgCl pellet electrodes (World Precision Instruments Inc., Sarasota, FL, USA) or Ag/AgCl wire electrodes. Ag/AgCl wire electrodes were chlorinated by immersing coiled silver wire in bleach for 1 h before rinsing thoroughly with demineralized water. After use electrodes were stripped from silver chloride with ammonium hydroxide and re-chlorinated. We assembled experimental setups inside Faraday cages. Voltages were applied externally through NI PXI-8336 & PXI-4461 controlled by in-house labview software, both applied voltage and resulting current were measured. We carried out all experiments at room temperature (23 to 25 °C) and tested all setups with different model cells to confirm proper functionality of the amplifiers. We cleaned the setups by rinsing all parts that came in contact with the electrolyte, nanopores and lipids with several cycles of demineralized water and isopropyl alcohol. We subjected PTFE cells and films to additional cleaning with chloroform to remove vacuum grease.

### Red blood cell assay

Defibrinated sheep blood was diluted in aerolysin-final-buffer until an optical density (OD) of 1 was reached. For each mL of blood, 1 μL of trypsin in agarose was added. The concentration of aerolysin wt and aerolysin mutants was determined by measuring the absorbance at 280 nm with a microvolume UV-Vis spectrophotometer, and all mutants were diluted to a concentration of 5 mM in aerolysin-final-buffer. The blood sample was combined with the pores in a 96-well plate with transparent bottom to a final sample volume of 100 μL per well with a final pore concentration of 0.5 μM or 1 μM. The samples were gently mixed using a pipette. Triplicates were prepared for each pore and pore concentration. Additionally, three control samples containing only blood and trypsin were included as controls. The OD of each sample was monitored over a period of 1 hour, with OD measurements taken every 15 seconds. To compare the activity of each mutant, the measurements were normalised by dividing by the initial absorbance value of each sample. The absorbance was then plotted over time to visualise the lysis activity.

### Cryo-EM

The aerolysin E254A-E258A sample in SMALP and cryo-EM grids were prepared as described previously^39^. The dataset was collected at the Dubochet Center for Imaging (Lausanne, CH) using the 300 kV TFS Titan Krios G4 equipped with a Cold-FEG and Falcon 4 detection. The dataset was collected in electron counting (EC) mode. The Falcon 4 gain references were measured before starting the data collection. The data collection was performed using the TFS EPU software packages. Movies were recorded at a nominal magnification of 120’000x, corresponding to 0.658 Å/pixel with defocus values ranging from 0.8 to 1.7. The exposure dose was set to 50 e/Å^2^. The datasets were processed in CryoSPARC^67^. The reported resolutions are based on the gold-standard Fourier shell correlation (FSC) = 0.143 criteria^68^ and local-resolution variations were estimated using CryoSPARC. For the model building, the aerolysin wt (9FM6) and the post-prepore (5JZW) structures were used as initial models which were fit into the cryo-EM map using Phenix^69^, manually adjusted using Coot^70^ and refined in Rosetta^69^ and Phenix. In the E254A-E258A pore structure (9GXJ) residues 14-23 & 246-251 and in the post-prepore (not quasi-pore) structure (9IGN), residues 197-308 were not built due to a lack of density in these areas. The details on the processing and refining process and the EMDB IDs can be found in SI Table 3.

### Section 3. Simulations

The systems for atomistic MD simulations were prepared with the CHARMM-GUI^70^ website, using the structure of heptameric aerolysin wt available in the PDB with ID 5JZT, onto which the desired mutations were crafted using PyMOL. The protein heptamers were placed in the DPhPC membranes (CHARMM’s lipid name PHPC) using PPM 2.0 through the CHARMM-GUI interface. The systems were further prepared (solvated and neutralised) with standard CHARMM-GUI procedures and parameters, in 1 M KCl. The systems were parameterized using CHARMM36m^71^ for the protein components and the corresponding version of TIP3P water, plus the standard CHARMM parameters available for DPhPC lipids. After standard minimization and equilibration procedures with standard parameters and restraint strengths using Gromacs 2022, we ran the production MD simulations applying the indicated voltages using the same MD engines applying semi-isotropic pressure coupling to 1 atm, a temperature of 298 K, 2 fs integration steps, LINCS-based restraints on hydrogens, and PME electrostatics with 12 Å cutoff. Restraints on the Cα atoms were also applied allowing sampling the effect of sidechain-contributed electrostatics on ion currents without interference from structural perturbations that might be unrealistic given the limited resolution of the starting PDB structure and the high applied voltages. Simulations were extended for around 400 ns; and for all subsequent analysis only the last 280 ns of the production phase were used. These analyses were carried out using standard VMD commands and custom scripts as detailed below.

***The ionic flux*** at a given voltage bias was extracted from whole-atom simulations using a previously published method^72^. In brief, the instantaneous ionic current at time *t* was

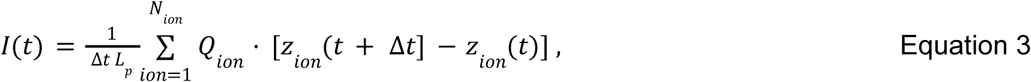

where *Δt* = 0.2 ns is the time gap between consecutive pairs of simulation snapshots written during production, *L_p_* is the length of the pore β-barrel (9 nm for aerolysin), *N_ion_* is the number of ions inside the pore lumen and *Q_ion_*and *z_ion_* are the charge and z-coordinate of ion, respectively. The average current at a given voltage bias was computed by applying a linear regression to the cumulative sum of *I(t)*.

### Section 4. Data Analysis

#### Two-state description

We simplified the gating process of nanopores as a stochastic switching between two discrete conductance levels, the open-pore state *G_O_(V)*, and the closed-pore state *G_x_(V)*. In accordance with the gating measurements on single pores (Figure 1), we set *G_x_(V) =* ɛ · *G_O_(V)* with ɛ = 0.14 which is the average gating level of 4 different recorded pores. In this simplified 2-state system, the probability of a pore to be closed, *p(t)*, is governed by Equation 2. To experimentally measure the mean voltage-dependent opening and closing rates, *k_O_* and *k_X_*, we applied a step-current with bias voltage *V(t) = V_app._* to an ensemble of pores. The fraction of open pores then decays exponentially as

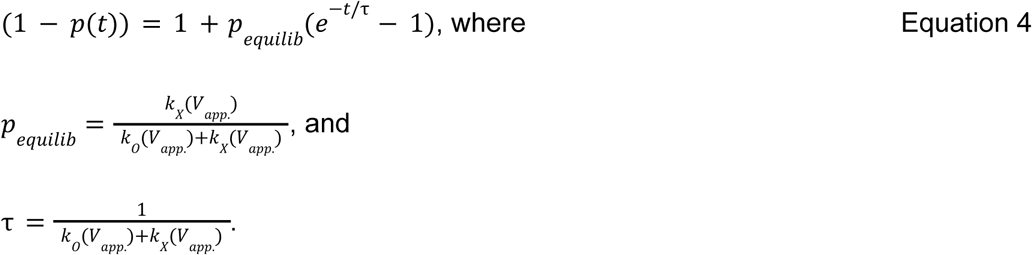

Thus, by applying voltage steps of various bias voltages to a pore ensemble and extracting *p_equilib_*and *τ* from the measured current response, the opening and closing rates were estimated. Exponential fitting was implemented as a weighted linear regression of

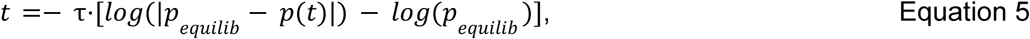

where the optimal *τ* and *p_equilib_* were those that minimised the weighted residual sum of squares. The weights decayed logarithmically from 1 (at *t =* 0) to 10^-15^ (at *t = t_end_ =* 35 s) to ensure a good estimation of *τ*.

At low bias voltages (*V_app._ <* 60 mV), where the closing rate is almost zero so that an accurate exponential fit becomes problematic, pores were first closed by applying a high bias voltage of 200 mV for 35 seconds, after which the voltage was reduced to *V_app._* and the opening rate was estimated as *k_O_(V_app._) = τ^-1^*.

At each measured bias voltage, estimates of *k_X_(V_app._)* and *k_O_(V_app._)* were averaged over three current responses to voltage steps recorded sequentially in the same membrane system.

### Alternating voltage traces

In the general case, assuming only 2 conductance states, the expected value for the conductance of an ensemble of gating pores *G(t)* is given by Equation 1. Given the 2-state assumption *G_x_(V) =* ɛ · *G_O_(V)*, the expected ensemble current, *I(t)*, is

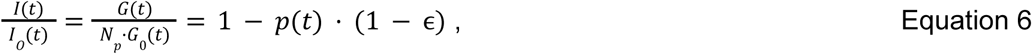

where *I_O_(t) = N_p_*⋅*G_O_(t)*⋅*V(t)* is the open-pore current obtained from AC measurements at high frequency. Thus, from a recording of the ensemble current response to an alternating voltage, the frequency-dependent closed-pore probability could be estimated as

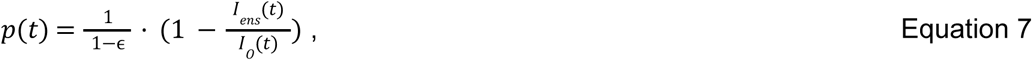

Where *I_ens_ (t)* is the average measured ensemble current of multiple AC cycles in the same ensemble. Note that for each cycle, the ensemble current was normalised by the estimated number of pores in that cycle, *N_p_*. To find the number of pores, a voltage window where no gating took place was predefined, and the current in this window was fitted to the *IV* curve in the same window using a linear regression, the slope of which gave *N_p_*. Throughout a cycle, the closing rate *k_X_* in Hz was estimated as the maximum of 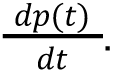

## Notes

### Competing Interest Statement

The authors have declared no competing interest.

### Summary of Updates

New theoretical model for gating have been added following reviewer's feedback

## References

1. Margheritis, E., Kappelhoff, S. & Cosentino, K. Pore-Forming Proteins: From Pore Assembly to Structure by Quantitative Single-Molecule Imaging. Int. J. Mol. Sci. 24, 4528 (2023).

2. Capone, R., Blake, S., Rincon Restrepo, M., Yang, J. & Mayer, M. Designing Nanosensors Based on Charged Derivatives of Gramicidin A. J. Am. Chem. Soc. 129, 9737–9745 (2007).

3. Butler, T. Z., Pavlenok, M., Derrington, I. M., Niederweis, M. & Gundlach, J. H. Single-molecule DNA detection with an engineered MspA protein nanopore. Proc. Natl. Acad. Sci. 105, 20647–20652 (2008).

4. Derrington, I. M. et al. Nanopore DNA sequencing with MspA. Proc. Natl. Acad. Sci. 107, 16060–16065 (2010).

5. Manrao, E. A. et al. Reading DNA at single-nucleotide resolution with a mutant MspA nanopore and phi29 DNA polymerase. Nat. Biotechnol. 30, 349–353 (2012).

6. Deamer, D., Akeson, M. & Branton, D. Three decades of nanopore sequencing. Nat. Biotechnol. 34, 518–524 (2016).

7. Brinkerhoff, H., Kang, A. S. W., Liu, J., Aksimentiev, A. & Dekker, C. Multiple rereads of single proteins at single–amino acid resolution using nanopores. Science 374, 1509–1513 (2021).

8. Wang, Y., Zhao, Y., Bollas, A., Wang, Y. & Au, K. F. Nanopore sequencing technology, bioinformatics and applications. Nat. Biotechnol. 39, 1348–1365 (2021).

9. Mayer, S. F., Cao, C. & Dal Peraro, M. Biological nanopores for single-molecule sensing. iScience 25, 104145 (2022).

10. Bertil Hille. Ion Channels of Excitable Membranes. (Sinauer Associates, Inc. Publishers, Sunderland, Massachusetts USA, 2001).

11. Bainbridge, G., Gokce, I. & Lakey, J. H. Voltage gating is a fundamental feature of porin and toxin β-barrel membrane channels. FEBS Lett. 431, 305–308 (1998).

12. Dessaux, D., Mathé, J., Ramirez, R. & Basdevant, N. Current Rectification and Ionic Selectivity of α-Hemolysin: Coarse-Grained Molecular Dynamics Simulations. J. Phys. Chem. B 126, 4189–4199 (2022).

13. Miedema, H. et al. A Biological Porin Engineered into a Molecular, Nanofluidic Diode. Nano Lett. 7, 2886–2891 (2007).

14. Cao, C. et al. Single-molecule sensing of peptides and nucleic acids by engineered aerolysin nanopores. Nat. Commun. 10, 4918 (2019).

15. Mueller, P. & Rudin, D. O. Induced excitability in reconstituted cell membrane structure. J. Theor. Biol. 4, 268–280 (1963).

16. Schindler, H. & Rosenbusch, J. P. Matrix protein from Escherichia coli outer membranes forms voltage-controlled channels in lipid bilayers. Proc. Natl. Acad. Sci. U. S. A. 75, 3751–3755 (1978).

17. Menestrina, G. Ionic channels formed byStaphylococcus aureus alpha-toxin: Voltage-dependent inhibition by divalent and trivalent cations. J. Membr. Biol. 90, 177–190 (1986).

18. Kasianowicz, J. J. & Bezrukov, S. M. Protonation dynamics of the alpha-toxin ion channel from spectral analysis of pH-dependent current fluctuations. Biophys. J. 69, 94–105 (1995).

19. Mohammad, M. M. & Movileanu, L. Impact of Distant Charge Reversals within a Robust β-Barrel Protein Pore. J. Phys. Chem. B 114, 8750–8759 (2010).

20. Nestorovich, E. M. & Bezrukov, S. M. Beta-Barrel Channel Response to High Electric Fields: Functional Gating or Reversible Denaturation? Int. J. Mol. Sci. 24, 16655 (2023).

21. Paulo, G. et al. Hydrophobically gated memristive nanopores for neuromorphic applications. Nat. Commun. 14, 8390 (2023).

22. Mayse, L. A. & Movileanu, L. Gating of β-Barrel Protein Pores, Porins, and Channels: An Old Problem with New Facets. Int. J. Mol. Sci. 24, 12095 (2023).

23. Colombini, M. Voltage gating in the mitochondrial channel, VDAC. J. Membr. Biol. 111, 103–111 (1989).

24. Bocquet, L. & Charlaix, E. Nanofluidics, from bulk to interfaces. Chem. Soc. Rev. 39, 1073–1095 (2010).

25. Maglia, G., Restrepo, M. R., Mikhailova, E. & Bayley, H. Enhanced translocation of single DNA molecules through α-hemolysin nanopores by manipulation of internal charge. Proc. Natl. Acad. Sci. 105, 19720–19725 (2008).

26. Hanke, W. & Schlue, W.-R. Planar Lipid Bilayers Methods and Applications. (Academic Press, 1993).

27. Bayley, H. & Martin, C. R. Resistive-Pulse SensingFrom Microbes to Molecules. Chem. Rev. 100, 2575–2594 (2000).

28. Strukov, D. B., Snider, G. S., Stewart, D. R. & Williams, R. S. The missing memristor found. Nature 453, 80–83 (2008).

29. Sung, C., Hwang, H. & Yoo, I. K. Perspective: A review on memristive hardware for neuromorphic computation. J. Appl. Phys. 124, 151903 (2018).

30. Kumar, S., Wang, X., Strachan, J. P., Yang, Y. & Lu, W. D. Dynamical memristors for higher-complexity neuromorphic computing. Nat. Rev. Mater. 7, 575–591 (2022).

31. Robin, P. et al. Long-term memory and synapse-like dynamics in two-dimensional nanofluidic channels. Science 379, 161–167 (2023).

32. Xiong, T. et al. Neuromorphic functions with a polyelectrolyte-confined fluidic memristor. Science 379, 156–161 (2023).

33. Emmerich, T. et al. Nanofluidic logic with mechano-ionic memristive switches. *Nat*. Electron. 1–8 (2024) doi:10.1038/s41928-024-01137-9.

34. Emmerich, T. et al. Nanofluidics. Nat. Rev. Methods Primer 4, 1–18 (2024).

35. Schindler, H. & Rosenbusch, J. P. Matrix protein in planar membranes: clusters of channels in a native environment and their functional reassembly. Proc. Natl. Acad. Sci. U. S. A. 78, 2302–2306 (1981).

36. Rappaport, S. M. et al. Conductance hysteresis in the voltage-dependent anion channel. Eur. Biophys. J. 44, 465–472 (2015).

37. Anton, J. S. et al. Aerolysin Nanopore Structures Revealed at High Resolution in a Lipid Environment. J. Am. Chem. Soc. 147, 4984–4992 (2025).

38. Cao, C. et al. Mapping the sensing spots of aerolysin for single oligonucleotides analysis. Nat. Commun. 9, 2823 (2018).

39. Anton, J. S. et al. Aerolysin nanopore structure revealed at high resolution in lipid environment. 2024.08.12.607338 Preprint at 10.1101/2024.08.12.607338 (2024).

40. Noskov, S. Y., Im, W. & Roux, B. Ion Permeation through the α-Hemolysin Channel: Theoretical Studies Based on Brownian Dynamics and Poisson-Nernst-Plank Electrodiffusion Theory. Biophys. J. 87, 2299–2309 (2004).

41. Maffeo, C., Bhattacharya, S., Yoo, J., Wells, D. & Aksimentiev, A. Modeling and Simulation of Ion Channels. Chem. Rev. 112, 6250–6284 (2012).

42. Vlassiouk, I. & Siwy, Z. S. Nanofluidic Diode. Nano Lett. 7, 552–556 (2007).

43. Karnik, R., Duan, C., Castelino, K., Daiguji, H. & Majumdar, A. Rectification of Ionic Current in a Nanofluidic Diode. Nano Lett. 7, 547–551 (2007).

44. Poggioli, A. R., Siria, A. & Bocquet, L. Beyond the Tradeoff: Dynamic Selectivity in Ionic Transport and Current Rectification. J. Phys. Chem. B 123, 1171–1185 (2019).

45. You, Y. et al. Angstrofluidics: Walking to the Limit. Annu. Rev. Mater. Res. 52, 189–218 (2022).

46. Zamyatnin, A. A. Protein volume in solution. Prog. Biophys. Mol. Biol. 24, 107–123 (1972).

47. Liu, N., Samartzidou, H., Lee, K. W., Briggs, J. M. & Delcour, A. H. Effects of pore mutations and permeant ion concentration on the spontaneous gating activity of OmpC porin. Protein Eng. Des. Sel. 13, 491–500 (2000).

48. Delcour, A. H., Adler, J. & Kung, C. A single amino acid substitution alters conductance and gating of OmpC porin ofEscherichia coli. J. Membr. Biol. 119, 267–275 (1991).

49. Ngo, V. A. et al. The Single Residue K12 Governs the Exceptional Voltage Sensitivity of Mitochondrial Voltage-Dependent Anion Channel Gating. J. Am. Chem. Soc. 144, 14564–14577 (2022).

50. Fologea, D. et al. Controlled Gating of Lysenin Pores. Biophys. Chem. 146, 25–29 (2010).

51. Krasilnikov, O. V., Muratkhodjaev, J. N. & Zitzer, A. O. The mode of action of *Vibrio cholerae* cytolysin. The influences on both erythrocytes and planar lipid bilayers. Biochim. Biophys. Acta BBA - Biomembr. 1111, 7–16 (1992).

52. Duret, G., Simonet, V. & Delcour, A. H. Modulation of Vibrio cholerae Porin Function by Acidic pH. Channels 1, 70–79 (2007).

53. Chimerel, C., Movileanu, L., Pezeshki, S., Winterhalter, M. & Kleinekathöfer, U. Transport at the nanoscale: temperature dependence of ion conductance. Eur. Biophys. J. 38, 121–125 (2008).

54. Queralt-Martín, M. et al. VDAC Gating Thermodynamics, but Not Gating Kinetics, Are Virtually Temperature Independent. Biophys. J. 119, 2584–2592 (2020).

55. Aidley, D. J. & Stanfield, P. R. Ion Channels: Molecules in Action. (Cambridge University Press, 1996).

56. Grosse, W. et al. Structure-Based Engineering of a Minimal Porin Reveals Loop-Independent Channel Closure. Biochemistry 53, 4826–4838 (2014).

57. Kaufman, I. K., McClintock, P. V. E. & Eisenberg, R. S. Coulomb blockade model of permeation and selectivity in biological ion channels. New J. Phys. 17, 083021 (2015).

58. Iacovache, I. et al. Cryo-EM structure of aerolysin variants reveals a novel protein fold and the pore-formation process. Nat. Commun. 7, 12062 (2016).

59. Andersen, O. S. & Koeppe, R. E. Bilayer Thickness and Membrane Protein Function: An Energetic Perspective. Annu. Rev. Biophys. Biomol. Struct. 36, 107–130 (2007).

60. Wiese, A. et al. Influence of the lipid matrix on incorporation and function of LPS-free porin from *Paracoccus denitrificans*. Biochim. Biophys. Acta BBA - Biomembr. 1190, 231–242 (1994).

61. Lakey, J. H. & Pattus, F. The voltage-dependent activity of Escherichia coli porins in different planar bilayer reconstitutions. Eur. J. Biochem. 186, 303–308 (1989).

62. Robin, P. et al. Long-term memory and synapse-like dynamics in two-dimensional nanofluidic channels. Science 379, 161–167 (2023).

63. Ramirez, P., Gómez, V., Cervera, J., Mafe, S. & Bisquert, J. Synaptical Tunability of Multipore Nanofluidic Memristors. J. Phys. Chem. Lett. 14, 10930–10934 (2023).

64. Kamsma, T. M. et al. Brain-inspired computing with fluidic iontronic nanochannels. Proc. Natl. Acad. Sci. 121, e2320242121 (2024).

65. Berhanu, S. et al. Sculpting conducting nanopore size and shape through de novo protein design. Science 385, 282–288 (2024).

66. Montal, M. & Mueller, P. Formation of Bimolecular Membranes from Lipid Monolayers and a Study of Their Electrical Properties. Proc. Natl. Acad. Sci. 69, 3561–3566 (1972).

67. Punjani, A., Rubinstein, J. L., Fleet, D. J. & Brubaker, M. A. cryoSPARC: algorithms for rapid unsupervised cryo-EM structure determination. Nat. Methods 14, 290–296 (2017).

68. Scheres, S. H. W. & Chen, S. Prevention of overfitting in cryo-EM structure determination. Nat. Methods 9, 853–854 (2012).

69. Wang, R. Y.-R. et al. Automated structure refinement of macromolecular assemblies from cryo-EM maps using Rosetta. eLife 5, e17219 (2016).

70. Lee, J. et al. CHARMM-GUI Input Generator for NAMD, GROMACS, AMBER, OpenMM, and CHARMM/OpenMM Simulations Using the CHARMM36 Additive Force Field. J. Chem. Theory Comput. 12, 405–413 (2016).

71. Huang, J. & MacKerell, A. D. CHARMM36 all-atom additive protein force field: validation based on comparison to NMR data. J. Comput. Chem. 34, 2135–2145 (2013).

72. Aksimentiev, A. & Schulten, K. Imaging *α*-Hemolysin with Molecular Dynamics: Ionic Conductance, Osmotic Permeability, and the Electrostatic Potential Map. Biophys. J. 88, 3745–3761 (2005).

73. Kavokine, N., Marbach, S., Siria, A. & Bocquet, L. Ionic Coulomb blockade as a fractional Wien effect. Nat. Nanotechnol. 14, 573–578 (2019).

74. Parsegian, A. Energy of an Ion crossing a Low Dielectric Membrane: Solutions to Four Relevant Electrostatic Problems. Nature 221, 844–846 (1969).

75. Long, D., Viovy, J.-L. & Ajdari, A. Simultaneous Action of Electric Fields and Nonelectric Forces on a Polyelectrolyte: Motion and Deformation. Phys. Rev. Lett. 76, 3858–3861 (1996).

76. van Dorp, S., Keyser, U. F., Dekker, N. H., Dekker, C. & Lemay, S. G. Origin of the electrophoretic force on DNA in solid-state nanopores. Nat. Phys. 5, 347–351 (2009).

77. de Souza, J. P., Levy, A. & Bazant, M. Z. Electroneutrality breakdown in nanopore arrays. *Phys*. Rev. E 104, 044803 (2021).

78. Laszlo, A. H., Derrington, I. M. & Gundlach, J. H. MspA nanopore as a single-molecule tool: From sequencing to SPRNT. Methods 105, 75–89 (2016).

79. Wilmsen, H. U., Pattus, F. & Buckley, J. T. Aerolysin, a hemolysin fromAeromonas hydrophila, forms voltage-gated channels in planar lipid bilayers. J. Membr. Biol. 115, 71–81 (1990).

80. Wurl, A. et al. Filling the Gap with Long n-Alkanes: Incorporation of C20 and C30 into Phospholipid Membranes. Langmuir 38, 8595–8606 (2022).

81. Zoni, V., Campomanes, P. & Vanni, S. Investigating the structural properties of hydrophobic solvent-rich lipid bilayers. Soft Matter 17, 5329–5335 (2021).

82. Yu, L. et al. Stable polymer bilayers for protein channel recordings at high guanidinium chloride concentrations. Biophys. J. 120, 1537–1541 (2021).

83. Vreeker, E. et al. Nanopore-Functionalized Hybrid Lipid-Block Copolymer Membranes Allow Efficient Single-Molecule Sampling and Stable Sensing of Human Serum. Adv. Mater. 37, 2418462 (2025).

84. Degiacomi, M. T. et al. Molecular assembly of the aerolysin pore reveals a swirling membrane-insertion mechanism. Nat. Chem. Biol. 9, 623–629 (2013).

85. Kavokine, N., Netz, R. R. & Bocquet, L. Fluids at the Nanoscale: From Continuum to Subcontinuum Transport. Annu. Rev. Fluid Mech. 53, 377–410 (2021).

