## Supplementary material for "Lumen charge governs gated ion transport in β-barrel nanopores": SI_text

**Theory notes**

**SI Section 1 Analytical theory of rectification in aerolysin**

The effect of ionic rectification, also known as nanofluidic diode, has been the study of many experimental and theoretical studies; it was also recently shown to be linked to gating and memory effects in solid-state nanopores[31]. However, these focused on relatively large channels (nm in diameter) compared to ours, with spatially-extended surface charges and geometrical asymmetries. Here, we develop a new theoretical framework to tackle the effect of individual, localised charged residues in a molecular pore.

We model aerolysin as a cylindrical pore of length $L_p$ = 9 nm and diameter $2r_p$ = 1.3 nm, with a distribution of surface charges corresponding to charged residues (Figure 3a). These charges act as the potential energy landscape to diffusing ions. When an electric field is applied, ions accumulate downstream of potential wells and upstream of potential barriers.

In the limit where the energy landscape is small enough, compared to thermal energy, so that it can be treated perturbatively, so that the overall current through the pores reads:

$$I \simeq G_0\left[1 + \beta(V_{app.})\right]V_{app.},$$ SI Equation 1

where the open pore is the limit and where we defined the rectification factor. In what follows, we show that, for a mutant where residues with charge at position in the wt are replaced with neutral residues, the rectification factor is given by:

$$\beta(V_{app}) \simeq \beta_{wt} - Q\frac{eV_{app}}{k_BT}\frac{L_p\lambda_B}{2r_p^2}\sum_{n=1}^{\infty}\frac{(-1)^n}{(\kappa_D^2L_p^2/4+\pi^2n^2)\pi n}sin\frac{2\pi n z_0}{L_p}$$ SI Equation 2

Where $\beta_{wt}$ is the rectification factor of the wt, $\kappa_D$ is the inverse Debye length and $\lambda_B$ the Bjerrum length (under typical conditions, $\kappa_D^{-1}$ = 0.3 nm and $\lambda_B$ = 0.7 nm). In what follows, we derive this expression.

We consider that the channel connects two macroscopic reservoirs filled with KCl with concentration $c_0$ = 1 M. The central axis of the channel corresponds to the z axis, with the channel going from $z = -L_p/2$ to $z = +L_p/2$. We assume that all quantities (potential, concentration, etc.) are uniform across the cross-section of the channel, and therefore use a strictly 1D model. We denote by $c_\pm(z)$ the local concentrations in cations and anions, and $V(z)$ the local electrostatic potential.

The boundary conditions are such that $V(z = \pm L_p/2) = \mp V_{app}$. The reservoirs also impose boundary conditions for the ionic concentrations: $c_\pm(z = L_p/2) = c_\pm(z = -L_p/2) = c_0$. To lighten up the notation, we temporarily work in such units that $r_p = \varepsilon = e = D = k_B T = 1$, with $D$ the diffusion coefficient of ions.

We first consider the case $V_{app} = 0$. In this case, the system is at thermal equilibrium, and the electrostatic potential $V$ solves the Poisson-Boltzmann equation:

$$\partial_{zz} V - \kappa_D^2 sinhV = -\rho,\ c_\pm = c_0 e^{\mp V_0}, \qquad \text{SI Equation 3}$$

where we recall that ρ is the distribution of fixed lumen charges. In what follows, we denote by $c_\pm^{(0)}$ and $V^{(0)}$ the solutions of these equations.

Assume now that we apply a non-zero voltage drop. The stationary Poisson-Nernst-Planck (PNP) equations become:

$$j_\pm = -\partial_z c_\pm \mp c_\pm \partial_z V$$

$$\partial_{zz} V = -\rho - c_+ + c_- \qquad \text{SI Equation 4}$$

With $j_\pm$ the (conserved) flux of cations/anions. This set of equations can be solved by the following change of variables, introducing the fugacities $f_\pm$:

$$c_\pm = f_\pm(z) e^{\mp V(z)}. \qquad \text{SI Equation 5}$$

Injecting this ansatz into the PNP equation yields:

$$f_\pm(z) = c_0 e^{\pm V_{app}} - j_\pm \int_{-L_p/2}^{z} e^{\pm V(y)}\, dy, \qquad \text{SI Equation 6}$$

from where we can extract the value of $j_\pm$ by enforcing the boundary condition at $z = L_p$:

$$j_\pm = \pm c_0 \frac{2 sinhV_{app}}{\int_{-L_p/2}^{L_p/2} e^{\pm V(y)}\, dy} \qquad \text{SI Equation 7}$$

We now assume that we are sufficient close to equilibrium, such that:

$$V(z) \simeq -V_{app}\frac{z}{L_p} + V^{(0)}(z). \qquad \text{SI Equation 8}$$

Note that this is a rather uncontrolled approximation, but by solving the PNP equations numerically we find that it is a good approximation even for $V_{app}$ = 100 mV even in presence of strong and localised surface charges. Then, we have:

$$\int_{-L_p/2}^{L_p/2} e^{\pm V(y)}\,dy = \int_{-L_p/2}^{L_p/2} e^{\mp V_{app} y/L_p} e^{\pm V^{(0)}(y)}\,dy, \qquad \text{SI Equation 9}$$

Defining $\Gamma^{\pm}(y) = e^{\pm V^{(0)}(y)}$, we obtain:

$$j_{\pm} = \pm\frac{c_0 V_{app}}{L_p} \times \left[\frac{V_{app}}{L_p \sinh V_{app}} \int_{-L_p/2}^{L_p/2} e^{\mp V_{app} y/L_p} \Gamma^{\pm}(y)\,dy\right]^{-1} \qquad \text{SI Equation 10}$$

We can simplify this expression by introducing the Fourier series of *Γ*:

$$\Gamma^{\pm}(z) = \sum_{n=-\infty}^{\infty} \Gamma^{\pm}_n e^{i\pi n z/L_p}, \qquad \text{SI Equation 11}$$

with the coefficient given by:

$$\Gamma^{\pm}_n = \frac{1}{2L_p} \int_{-L_p/2}^{L_p/2} \Gamma^{\pm}(z) e^{-2i\pi n z/L_p}\,dz. \qquad \text{SI Equation 12}$$

We obtain:

$$j_{\pm} = \pm\frac{c_0 V_{app}}{L_p} \times \left[\Gamma^{\pm}_0 + \sum_{n\neq 0} \frac{\mp V_{app}(-1)^n}{i\pi n \mp V_{app}} \Gamma^{\pm}_n\right]^{-1} \qquad \text{SI Equation 13}$$

Since we only work at order 2 in $V_{app}$ and that $\Gamma_{-n} = \Gamma_n^*$, we obtain:

$$j_{\pm} \simeq \pm\frac{c_0 V_{app}}{L_p} \times \left[\Gamma^{\pm}_0 \mp V_{app} \sum_{n=1}^{\infty} (-1)^n \frac{2}{\pi n} Im\Gamma^{\pm}_n\right] \qquad \text{SI Equation 14}$$

The second term on the right-hand side describes rectification. Since $Im\Gamma_n$ describes the asymmetry of the potential created by fixed lumen charges, rectification is hence directly linked to the shape of the lumen charge distribution.

This expression of $j_\pm$ is not extremely helpful, so now we try to derive a more effective form.

By solving numerically the full PNP equations, we find that the Debye-Hückel limit is sufficient to capture all the relevant physics considering our parameters, despite the fact that we work with high concentrations and localised surface charges. We hypothesised that the reason is the strong screening of lumen charges, so that the energy landscape of the pore remains not too large with respect to $k_BT$. In this limit, it is easy to express the potential inside the pore $V^{(0)}$ as function of the lumen charge distribution ρ in Fourier space:

$$V_n^{(0)} = \frac{\rho_n}{k_n^2+\kappa_D^2}, \qquad \text{SI Equation 15}$$

With $k_n = 2\pi n/L_p$. For a given mutant, we can analyse ρ as the charge distribution $\rho_{wt}$ plus a contribution $-Q\delta(z - z_0)$ from the mutation, so that

$$\rho_n = \rho_{wt,n} - \frac{Q}{L_p}\left(1 - (-1)^n e^{-\kappa_D L_p/2}\right)e^{+2i\pi n z_0/L_p} \qquad \text{SI Equation 16}$$

Besides, the $\Gamma$'s can be expanded as:

$$\Gamma_\pm \simeq 1 \pm V^{(0)}. \qquad \text{SI Equation 17}$$

The total ionic current is $j = j_+ - j_-$ , so the rectification factor is given by:

$$\beta = \frac{j(V_{app})+j(-V_{app})}{j(V_{app})-j(-V_{app})}. \qquad \text{SI Equation 18}$$

We finally obtain:

$$\beta = \beta_{wt} - V_{app}\sum_n \frac{QL_p}{2\pi n}\frac{(-1)^n-e^{-\kappa_D L_p/2}}{\kappa_D^2 L_p^2/4+\pi^2 n^2} sin\frac{2\pi n z_0}{L_p} \qquad \text{SI Equation 19}$$

This derivation was performed using dimensionless units. To recover a physically meaningful result we must replace $V_{app}$ in the above equation by $eV_{app}/k_BT$ and $Q$ by $Q\lambda_B/r_p^2$; upon doing so, we obtain the announced result.

The main limitation of this model is that it assumes that the potential landscape within the pore is small compared to $k_BT$, such that we can treat its effects perturbatively. This approach could be criticised in several ways. First, we use the linearized Poisson-Boltzmann equation despite using highly concentrated solutions ($c_0$ = 1 M) in experiments. In addition, since aerolysin is a heptamer, any charged residue corresponds to a surface charge of $Q = \pm 7$. However, these high surface charges are balanced by the strong ionic screening due to the high salt concentration. By solving the full Poisson-Boltzmann equation numerically, we find that in practice we can account for all nonlinear effects by replacing the charge $Q$ induced by mutated residues by $Q^* \simeq 0.85 \cdot Q$.

Lastly, one may criticise our choice of a single, average value $r_p$ for the pore's radius, rather than considering variations of the channel's diameter. This approximation is valid as rectification is the result of accumulation of ions across the entire length of the pore upstream or downstream of lumen charges; the effect is therefore sensitive to the entire volume of the pore rather than to its radius at a specific point along the channel.

**SI Section 2. Gating model**

In this Section, we build an analytical model for gating in aerolysin. We start by defining the geometry and notations in Section 5.1, then we review experimental findings to constrain the model in Section 5.2. We use these observations to build a statistical mechanics model of gating in Section 5.3. We argue that gating itself is caused by depletion interactions between ions and the pore, resulting in a structural bistability. We then show that the closed state can be stabilized by an external electric field due to a local breakdown of electroneutrality in the closed state. In Section 5.4, we analyze how these results compare to experiments and propose reasonable values of model parameters.

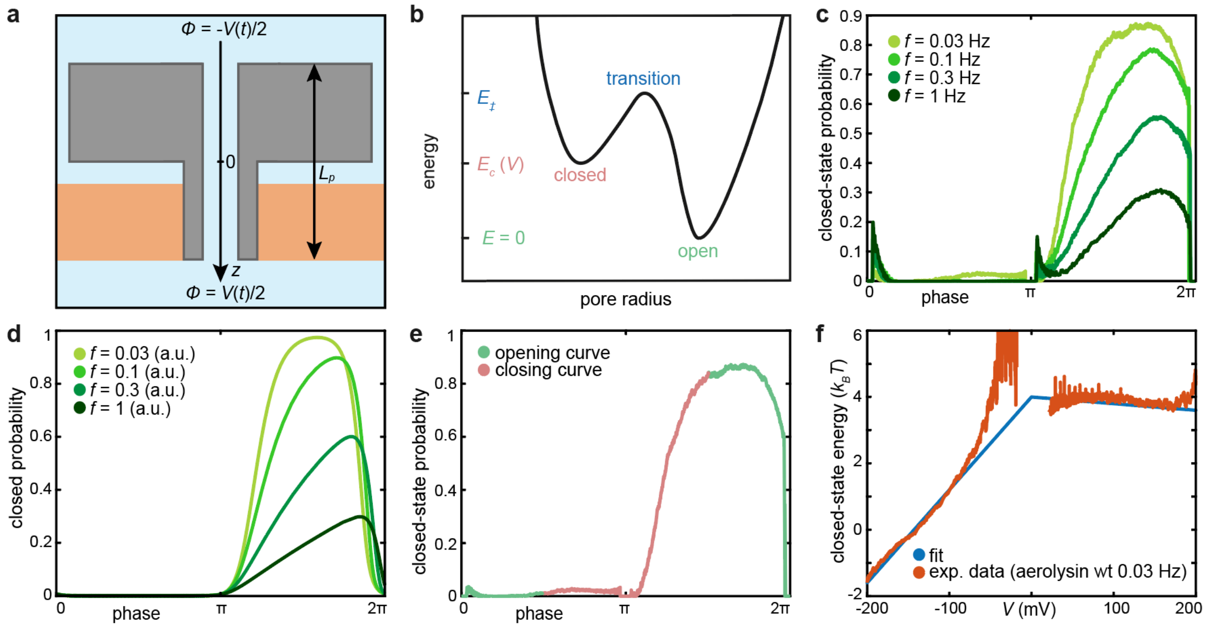

**SI Figure 1 | Constraining the biophysical model. a.** Geometry of the system. **b.** Two-state toy model of gating: the system is transitioning between states corresponding to different values of the pore radius, with transition rates given by the states' energies. **c.** Evolution of closed state probability with frequency. Experiments performed on aerolysin wt, amplitude 200 mV, 25 °C, 1 M KCl, pH 6.2 buffered with 10 mM phosphate. **d.** Effect of frequency in the two-state toy model, suggesting that only the energy of the closed state varies with the external voltage. **e, f.** Determination of the energy of the closed state, from the opening branch of the probability-voltage curve.

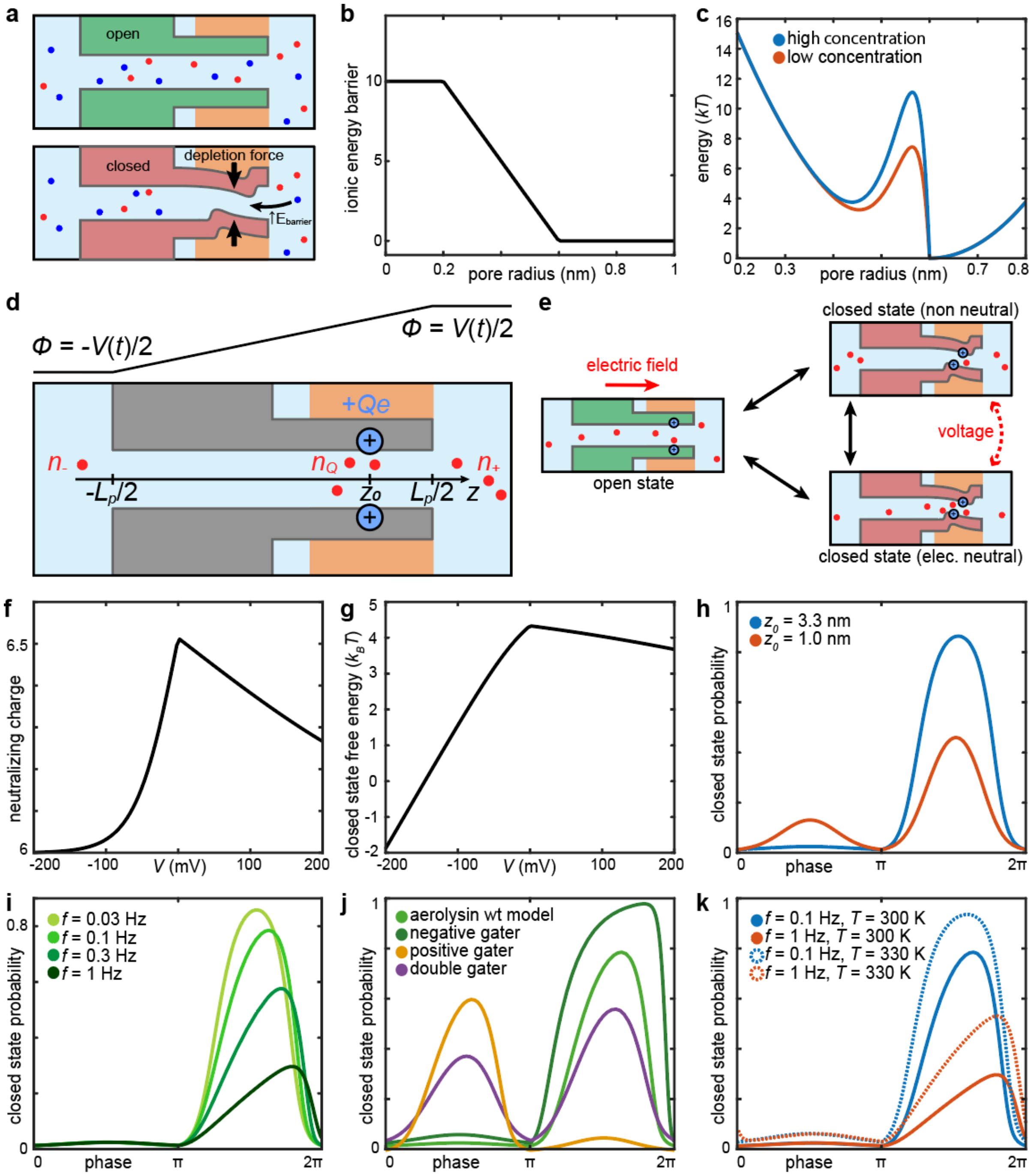

**SI Figure 2 | Model of ionic gating in aerolysin. a.** Schematic representation of depletion interactions. In the open state, the pore is relatively wide and permeable to ions. In the closed state, ions face an energy barrier due to the smaller radius, resulting in depletion interactions. **b.** Simplified energy profile for the ionic energy barrier $\mu^{ex}$. **c.** Energy landscape of the pore under zero voltage, due to depletion interactions (orange curve: concentration 1 M; blue curve: concentration 1.2 M). **d.** Parametrization of the gating model **e.** Schematic representation of the model's derivation. **f-k.** Results of the model, showing the number of counterions neutralizing the fixed lumen charge (**f**), the free energy of the closed state (**g**) and the closed state probability for various lumen charge positions (**h**, **j**), voltage frequency (**i**) and temperature (**k**). All parameters are detailed in the text.

#### 2.1 *Geometry and notations*

In all that follows, we consider a cylindrical pore representing aerolysin embedded in a lipid membrane. We use the geometry represented in SI Figure 1a: the pore is vertical, aligned with a $z$ axis pointing downward. The origin $z = 0$ is located in the middle of the pore's length (so that the bottom half of the pore corresponds to $z > 0$, and the top to $z < 0$ – note the sign convention here). The total length of the pore is $L_p = 9$ nm, so that $z$ runs from $- L_p/2$ to $+ L_p/2$. It also has a radius $R$, that can change due to conformational changes. In the open state, $R = r_p \sim 0.6$ nm. When we apply an external voltage $V(t)$, it is applied from an electrode at $z < 0$ to one at $z > 0$. Through the polarization of the reservoirs, this is equivalent of setting an external electrostatic potential $\phi(z)$ such that $\phi = V(t)/2$ for $z > L_p/2$, $\phi =- V(t)/2$ for $z <- L_p/2$, and $\phi = V(t)z/L_p$ in between.

#### 2.2 *Review of experimental findings and constraining the model*

<u>Effect of frequency: Only the closed state depends on applied voltage</u>

To characterize how the model should behave, we first consider a generic toy model of gating. Consider a pore that can exist in two states, closed and open, with respective probabilities $p$ and $1 - p$. We assume that the pore performs some Markovian dynamics between the two states, and write:

$$\frac{dp}{dt} = \frac{1-p}{\tau_1} - \frac{p}{\tau_2}, \qquad \text{SI Equation 20}$$

where $\tau_1$ and $\tau_2$ are the closing and opening times, respectively. We assume that the system is under an external voltage $V(t) = V_0 sin\omega t$, and that $\tau_1(V)$ and $\tau_2(V)$ depend only on the instantaneous value of voltage. We write:

$$\tau_2 = \tau e^{E_{\ddagger}-E_c(V)} \quad ; \quad \tau_1 = \tau e^{E_{\ddagger}-E_o(V)}, \qquad \text{SI Equation 21}$$

with $E_c$ and $E_o$ the energies (in $k_B T$ – note that these quantities can be negative) of the closed and open states, $E_‡$ is the energy of the transition state (see SI Figure 1b), and $\tau$ a typical molecular switching attempt timescale (Note that the opening time depends on the energy of the closed state, and vice-versa). As energies are defined up to an additive constant, we choose the open state under zero voltage as the reference $E_o(V = 0) = 0$. We test several possibilities for the shapes of these energies with voltage.

First of all, we look at the experimental curves for $p(t)$ for different values of the voltage frequency, see SI Figure 1c. We see that the closing portion of the curve ($p$ growing from 0 to 1) actually depends on frequency, while the opening portion does not. This shows that there is a rather large energy barrier to go from the open state to the closed state, and if $f > 1/\tau_1$, we see a depency in frequency. Even at large voltages (in absolute value), the system never suddenly jumps from the open state to the closed state. This means that $E_o$ remains very negative. On the other hand, the opening curve is very abrupt and does not depend on frequency. This is a sign that $E_c$ may increase for certain values of applied voltage, to the point where there is almost no energy barrier to switch from the closed state to the open one.

In other words, gating occurs because the energy of the closed state changes with voltage. Starting in the open state, the system faces a great and constant energy barrier to close. Whenever the energy of the closed state becomes less than that of the open state, the system closes, over a timescale that is limited by the energy of the transition state. Starting in the closed state, the system opens suddenly when the closed state becomes destabilized by the applied voltage.

We confirm this by simulating the above toy model for a constant $E_‡ = 10$, $E_c = 5 + V$, $V_0 = 9$, $\tau = 3 \times 10^{-5}$, and $\omega = 0.03, 0.1, 0.3, 1$, see SI Figure 1d.

Consequently, in all that follows, we assume that $E_o(V) = 0$, that $E_‡$ is a constant independent of voltage and that only the energy of the closed state, $E_c$, varies with voltage.

Effect of voltage: The closed state is only stabilized by the applied voltage

Next, we use the experimental data to observe the approximate shape of $E_c$ with applied voltage. Sadly, the dynamics of the pore is very slow, so we do not have access to the quasi-static limit $p_\infty(V)$, which would be the closed state probability under a constant voltage $V$. Instead, we make the following approximation:

$$p_\infty(V) \simeq p(V(t)), \qquad \text{SI Equation 22}$$

which is valid at low frequency ($f < 0.1\,\text{Hz}$), and only for the beginning of the closing curve (SI Figure 1e). We extract the closed state energy by:

$$E_c(V) = log\left[\frac{1}{p(V)} - 1\right] + constant \qquad \text{SI Equation 23}$$

We choose to study the aerolysin wt at $f = 0.03\,Hz$. In SI Figure 1f, we plot the closing probability $p(t)$ (obtained by an ensemble average over many experiments), then $E_c$ estimated from the closing curve. We see that the energy decreases with voltage at negative voltages (resulting in a stabilization of the closed state), but is roughly constant (actually, slightly decreasing) at positive voltage.

A good phenomenological fit (shown in SI Figure 1f, blue line):

$$E_c(V) \simeq 4 + 28\,min(V, 0) - 2\,max(V, 0). \qquad \text{SI Equation 24}$$

This is actually quite surprising. If this energy was interpreted as the electrostatic potential energy of a fixed charge on the pore's structure, one would expect $E_c \propto V$: an external field would stabilize the charge at one polarity, and destabilize it under the opposite polarity. An interesting remark is that the branch of $E_c(V)$ for $V > 0$ is not entirely flat: it has a weak negative slope, that is visible the the graph of the closed state probability, in the form of a secondary bump (see Figure 4, for $0 < \omega t < \pi$). This means that a channel that mainly gates at negative voltage (like aerolysine wt) will eventually gate under a strong enough positive voltage. This is counterintuitive, as it shows that the closed state is maximally unstable at $V = 0$. This suggests that the closed state is in fact stabilized by some charge dissociation effect, as this kind of phenomenon would also be highly unlikely under zero voltage.

Lastly, this asymmetric $E_c(V)$ also qualitatively explains why double gaters can exist. Let us assume that gating in aerolysin wt is caused by a charged group at the bottom of the pore. Consider now a mutant that introduces a similar group, symmetrically at the top of the channel. If the two groups are independent, the total energy profile of this mutant must be $E_t \sim E_c(V) + E_c(-V)$, from symmetry considerations. If we had $E_c(V) \propto V$, then the energy increase of the group at the top under a negative voltage would cancel out the energy decrease of the group at the bottom, and vice-versa under a positive voltage, resulting in no gating at all.

Instead, since the branch $E_c(V > 0)$ is quite flat, the gating effects of two opposite groups should be able to coexist in an additive manner, rather than cancelling out.

Constraints to be satisfied by the model

Overall, from the above discussion, we conclude that a model of gating should account for the following observations:

1. The channel is inherently bistable.
2. The closed state is 'disordered', in the sense that many closed states coexist.
3. Only the energy of the closed state is modified by the external voltage.
4. The closed state is stabilized by a voltage of a certain polarity, but flipping the polarity does not destabilize the closed state.
5. Charged groups on opposite ends of the pore should contribute more or less additively to gating, resulting in double gaters.
6. Groups near the mouth of the pore have the most impact on gating.
7. The closed state is stabilized for voltages that also correspond to high open pore conductance.
8. Even negative gaters (e.g. aerolysin wt, see Figure 4 of main text) show some weak positive gating. However, the system seems to "transition" between different closed states rather than staying in a single closed state.

**2.3** ***The model: Depletion interactions, electro-osmosis and charge separation***

We now derive a model of nanopore gating. First, we show how depletion interactions (SI Figure 2a) can induce a mechanical bistability, even in a “simple” cylindrical pore with no movable parts. Then, we discuss the impact of lumen charges on the stability of the open and closed states under an external voltage. We argue that electro-osmotic forces cancel out electrostatic forces in the open state, but that a local breakdown of electroneutrality can occur in the closed state, lowering its electrostatic energy under an applied voltage.

Mechanical bistability at zero voltage: Depletion interactions

As a first step, we consider a cylindrical pore of length $L_p$ and radius $R$, without any surface charge and at zero voltage.

Ions typically face high energy barriers when entering nanometer-scale pores. In the limit $L_p \gg R$, one can compute analytically the electrostatic contribution to this energy barrier, which amounts to an excess chemical potential for ions inside the pore[73,74]:

$$\mu^{ex} \simeq 2\alpha \frac{\ell_B}{R} k_B T, \qquad \text{SI Equation 25}$$

where $\ell_B = 0.6$ nm is the Bjerrum length of water and $\alpha$ is a numerical coefficient that depends on the dielectric contrast $\varepsilon_m/\varepsilon_w$ between the membrane and water. Typically, $\alpha$ is in the range $1 - 10$. We know that the open state has $R = 0.65$ nm; the structure of the closed state is not known but we can assume that it corresponds to a reduction of $R$ by, for example, 30%, to $R = 0.4$ nm. This corresponds to an increase of $5k_BT$ of the energy barrier. Note that this energy barrier stems from purely electrostatic considerations, i.e. the fact that lipid membranes have a low dielectric constant. Other factors need to be considered, such as van der Waals interactions, deformations of the solvation shell, the pore’s surface charge etc. Another important caveat is that this energy is a “bare” self-energy faced by ions entering the pore because of their charge. When many ions enter the pore, screening will tend to greatly counteract this energy barrier.

To simplify the discussion, we assume that the energy barrier faced by an ion entering the pore varies linearly with $R$, interpolating between $\mu^{ex} = 0$ at $R > R_1 = 0.6$ nm and $\mu^{ex} = 10\, k_B T$ at $R < R_0 = 0.2$ nm, as depicted in SI Figure 2b. This is of course an oversimplification, but we expect that the consequences of choosing a different energy profile will be minor on the overall results.

Next, we assume that, for a given value of $R$, the pore spontaneously fills with ions with concentration $c_0 e^{-\mu^{ex}(R)}$, where $c_0 = 2\,\mathrm{M}$ is the imposed ion concentration (in cations and anions, hence a factor 2) in the reservoir. We then write the total free energy of the system (pore structure and ions):

$$E_t(R) = \pi R^2 L c_0 e^{-\mu^{ex}(R)} \mu^{ex}(R) + \pi K (R - r_p)^2, \qquad \text{SI Equation 26}$$

where the open pore radius $r_p$ is assumed to be the equilibrium radius of the pore, and $K$ is some elastic modulus. In what follows, we will use the value $K \sim 30\, k_B T/nm^2$, but this quantity is difficult to estimate (and likely depends on the mechanics of the β-barrel). This system shows some bistability, as depicted in SI Figure 2c: if the pore is fully open, the free energy barrier vanishes and ions fill the pore. If it starts closing, however, ions face an unfavourable energy barrier, and it becomes favorable to expel all ions and close the pore. This results in a secondary energy minimum at $R \sim 0.4\,\mathrm{nm}$, corresponding to the closed state. With the radius reduction and the increased energy barrier, this closed state has a very low conductance compared to the open state. This simplified model also predicts that, in absence of voltage (which we will discuss now), the energy difference between the open and the closed state is about $\Delta E \sim 5\, k_B T$ ($p_{closed} \sim 0.5\%$, which is reasonable), and a transition state energy of $E_{\ddagger} = 8\, k_B T$, suggesting that transitions between the two states are very slow: for a molecular attempt time $\tau \sim 1\,\mathrm{ms}$, the pore switches from the open to the close state in an average of $\tau e^{E_{\ddagger}} \sim 3\mathrm{s}$.

The model is schematically depicted in SI Figure 2a.

We now discuss how this picture is modified in presence of an external voltage $V$. Intuitively, any effect of the external voltage that is asymmetric (in the sense that reversing the voltage bias changes the measurement) has to be linked to some charge distribution along the pore. We therefore consider how the energy of those charges is affected by an external field.

Absence of energy variation in the open state: Electro-osmosis

Experimentally, we see that the open state does not seem to depend on voltage. This seems counterintuitive as one would expect that lumen charges acquire an energy $qVz/L_p$ (with $z$ the position of the charge relative to the middle of the pore's length, see SI Figure 1a). However, this effect is fully cancelled out by counterions. An external field $E$ will exert a force $qE$ on surface charges, and an opposite force $-qE$ on the fluid inside the pore due to the presence of counterions. In the steady state, this force drives an electro-osmotic flow through the pore, which entirely dissipates this force through viscous stress on the pore's structure. If one then very slowly turns on the voltage $V$, no net force will be exerted on the pore's structure (although the flow might induce some elastic deformations). At first order, one therefore expects that the free energy remains unchanged. At best, the open state could be slightly destabilized by elastic deformations, but that effect would persist regardless of the sign of applied voltage, as the electro-osmotic flow exists in both cases.

This argument can be put on rigorous theoretical grounds[75,76]. The general result is that a charged but non-flexible structure immersed in water does not feel a net force when subjected to an external field (in contrast with, say, pores with a "mobile" charged arm like in the ball-and-chain model).

Closed state: voltage-dependent electroneutrality breakdown

The argument above suggests that an electroneutrality breakdown could result in a change in free energy with voltage (since without electroneutrality, counterions no longer compensate the force of the external field ). Such a possibility has already been discussed in great detail by Bazant and coworkers in several works[77]. Overall, one can reasonably expect a partial breakdown of electroneutrality in pores of sizes similar to aerolysin ($2L_p \sim 10$ nm and $R \lesssim 1$ nm). The computations by Bazant and coworkers amounts to solving the Poisson-Boltzmann equation in a cylindrical pore with a continuous surface charge. They find that, for $R$ small enough, a charged pore can break electroneutrality, with some of the counterions remaining outside the pore (near its mouths) rather than inside. It does not apply directly here, since we consider a system out of equilibrium under an external field, and because we consider strongly localized lumen charges.

However, in the closed state, the ionic current through the pore almost vanishes, so the system remains "almost" at equilibrium, and the system is so small that one can essentially reduce it to a 1D model. As a first step, we consider a pore of length $L_p$ with a single charged group of $Q = 7$ positive charges at position $z_0 \in [-L_p/2, L_p/2]$. We will discuss how this connects to experimental systems in Section 5.4.

The external voltage creates a potential field $\phi(z) = zV/L_p$. We consider the distribution of negative counterions around the fixed charge $Q$. We assume that the system can partially break electroneutrality, in which case some of the counterions may localize away from the fixed charge; however, the system must be electroneutral at large scales, and so the number of counterions is fixed to $Q = 7$ overall (plus a fluctuating background of positive and negative ions, that we do not consider here). This idea lets us treat the system (pore + counterions) like a closed thermodynamic system, even though it is not fully at equilibrium: there is still a non-zero conduction of ions through the closed pore, and we "label" counterions even though they can be replaced by another ion of the same sign.

We assume that counterions can either localize exactly around the charge, or escape the pore and break electroneutrality (by localizing at $z = \pm L_p/2$). We note $n_Q$, $n_-$ and $n_+$ the number of counterions at $z = z_0$, $z = -L_p/2$ and $z = +L_p/2$, respectively. In other words, the closed pore corresponds to a set of microstates, indexed by $(n_Q, n_+, n_-)$, with different energy levels. When the voltage $V$ is applied, some of these states may become more stable, increasing the overall probability to be in one of the many closed states. We represent this toy model in SI Figure 2d-e. The total energy of one of these "microstates" is then:

$$E = - s_Q e \frac{V}{2}(n_+ - n_-) - s_Q \frac{eVz_0}{L_p}(n_Q - Q) + E_0 (n_Q - Q)^2 - \mu^{ex}(Q - n_Q), \quad \text{SI Equation 27}$$

with $s_Q = +1$ the sign of the surface charge. $E_0$ corresponds to a penalty for breaking electroneutrality; it can be interpreted as the self-energy of a bare, uncompensated charge within the pore, as discussed above. This term scales like the square of the uncompensated charge, in the limit where the surface charges are close to each other. The last term corresponds to the gain in free energy for removing one ion from the channel, denoted $\mu^{ex}$ as discussed for the depletion interactions. This term does not impact the results of the model, but is included here for the sake of internal consistency with our model of depletion interactions. One has $n_+ + n_- + n_Q = Q$ (as we are only considering counterions). For

$V > 0$, a non-zero value of $n_-$ is strictly unfavourable, so to simplify the discussion we assume that $n_- = 0$ for $V > 0$ and $n_+ = 0$ for $V < 0$. We obtain:

$$E(n_Q) = - e\frac{|V|}{2}(Q - n_Q) - s_Q e\frac{Vz_0}{L_p}(n_Q - Q) + E_0(n_Q - Q)^2 - \mu^{ex}(Q - n_Q).$$ SI Equation 28

Note that this expression is valid regardless of the sign of the surface charge or of the applied voltage. The probability of observing $n_Q$ counterions neutralizing the surface charge $Qe$ is then obtained by computing the Boltzmann weight $p = \binom{Q}{n_Q} e^{-E(n_Q)}/Z$, with $Z$ the partition function:

$$Z = \sum_{n_Q} \binom{Q}{n_Q} e^{-E(n_Q)},$$ SI Equation 29

and the free energy:

$$F(V) = - k_B T log Z.$$ SI Equation 30

This free energy should be interpreted as an energy difference between the closed state at $V = 0$ and at $V(t) = V_0 sin 2\pi f t$. Note that we treated counterions as differentiable particles. This is reasonable as the fixed charge $Q$ is actually made of 7 individual amino acids at slightly different positions; we can identify each bound counterion by the fixed charge it is "attached to." Note that treating the ions as non-differentiable particles would have almost no effect on the overall results.

We show the results of the model in SI Figure 2f-g. For this figure, the parameters are set to $Q = 7$, $L_p = 9$ nm, $z_0 = 3.3$ nm, $V_0 = 200$ mV, $E_0 = 7k_B T$ and $\mu^{ex} = 5\,k_B T$, which are meant to reproduce the aerolysin wt pore. We show the average neutralizing charge as function of applied voltage, as well as the free energy of the closed state (relative to the open state), defined as:

$$E_c(V) = E_{c,0} + F(V),$$ SI Equation 31

where $E_{c,0} = 5\,k_B T$ has been estimated using the model of depletion interactions. We find that the breakdown of electroneutrality is only favoured when the electric field within the pore tends to drag counterions across a large portion of the pore. This polarity of the external field matches the one associated with the high conductance state (caused by ionic rectification), in perfect accordance with experimental observations.

Gating

One may now gather the different results obtained so far. From the depletion interaction model, we know that the system is inherently bistable. Taking the open state as the energy reference, the closed state under zero voltage corresponds to an energy level $E_{c,0} \sim 5k_BT$. Under an applied voltage, the energy of the closed state is shifted by $F(V)$, which we can compute analytically, while the energy of the open state remains unchanged. The transition state between the two has an energy $E_{\ddagger} \sim 8\,k_BT$, as computed from the depletion interaction model (see SI Figure 2c). We assume that the typical molecular attempt time is $\tau$. We can now solve:

$$\frac{dp}{dt} = \frac{1-p}{\tau_1} - \frac{p}{\tau_2}, \quad \text{SI Equation 32}$$

with

$$\tau_1 = \tau e^{E_{\ddagger}} \quad \text{SI Equation 33}$$

and

$$\tau_2 = \tau e^{E_{\ddagger} - E_{c,0} - F(V(t))}. \quad \text{SI Equation 34}$$

We solve this model for the set of parameters given in the previous sections, and $\tau = 0.3$ ms. We first solve this equation for $f \rightarrow 0$, and represent the closed state probability $p$ as function of the voltage phase $\varphi = 2\pi f t$ in SI Figure 2h, for $V_0 = 200$ mV (blue curve). We recover in particular the existence of a "secondary peak" of gating under high positive voltage ($\varphi \simeq \pi/2$), that is linked to the weak negative slope of $F(V)$ at $V > 0$, as seen in experimental data. This secondary peak becomes even more visible under a stronger voltage, or for a surface charge placed closer to the center of the channel (e.g. $z_0 = 1.0$ nm, SI Figure 2h, orange curve). Qualitatively, breaking electroneutrality is only slightly favorable under high positive voltage: one allows negative counterions to move to $z = L_p/2$, where $\phi = V_0/2$ (favorable), but we expose a positive surface charge at $z = z_0 \simeq L_p/2$

(unfavourable) in the process. Because $z_0 < L_p/2$, the outcome is still favorable, but barely. This effect also becomes stronger for $z_0 \simeq 0$, as the charge distribution of the pore becomes more symmetrical.

For $z_0 < 0$, e.g. $z_0 =- 3.3$ nm, the effect is reversed and the channel instead gates under positive voltage. The same situation is observed for $z_0 > 0$ and a negative surface charge. Lastly, in SI Figure 2i, we show the effect of voltage frequency: our model recapitulates the phenomenological features found in experiments.

#### 2.4 *Comparison with experimental data*

The previous section proposed a set of parameters that aimed at representing the aerolysin wt pore. We now discuss how the model can be extended to account for the various mutants considered in the study.

Pores with multiple surface charges

So far, we considered the effect of a single, positive, 7-valent surface charge located at $z_0$. We found that setting $z_0 \simeq 3.3$ nm allowed to recover most of the phenomenology observed in aerolysin wt. We should stress that our model involves a few free parameters, but we were able to obtain quantitatively reasonable results by estimating them in orders of magnitude.

One may then use this model to describe the various mutants of aerolysin. In particular, single mutants, which replace a charged group by a neutral one, are equivalent to adding another 7-valent charge (of sign opposite to the one being deleted) along the channel. We will consider the following examples:

1. D222N, which amounts to adding a positive charge at the top of the channel,
2. E258A, which amounts to adding a second positive charge at the bottom (close to the effective aerolysin wt charge),
3. K242A and K238A, which add a negative charge at the bottom,
4. The double mutant K242A-K238A, which add two positive surface charges at the bottom.

Note that each time, we mean a 7-valent surface charge.

In the first two cases, as a first approximation, we can treat the aerolysin wt charge and the newly added charge as more or less independent, as they have the same sign and their effect on the overall free energy of the pore can reasonably be added (neglecting an unfavourable interaction between the two charged groups if they both break electroneutrality at the same time). Therefore, we could sum the result previously obtained for $F(V)$, the free energy contribution of the aerolysin wt charge, with a similar contribution from a charge located at $z_1$. The value of $z_1$ can be obtained from the pore's structure, with $z_1 =- 1.7$ for D222N and $z_1 = 4.0$ for E258A. Such a procedure yields a good qualitative agreement with the experimental data, albeit not a quantitative one. We hypothesize that the main issue lies in the fact that we neglected interactions between surface charges of similar signs, which are unlikely to break electroneutrality at the same time in practice, as that would be electrostatically unfavourable.

The situations where this additive procedure would include a positive and a negative charge (e.g. K242A and K238A, which have the aerolysin wt positive charge, and a negative charge due to a deleted positive charge) are even trickier: by neglecting interactions, we disregard the fact that those two charges actually cancel each other out. This means that the system (pore's structure + counterions) can no longer be treated as a closed thermodynamic system, as two counterions of opposite charge (one coming from the aerolysin wt charge, one from the mutated group) could effectively 'annihilate' each other (i.e. 'disappear into the ion background concentration'), leaving a pair of opposite surface charge groups. A detailed model of this effect is outside the scope of the present work. As a first approximation, we will consider that these pores do not gate, as their surface charges cancel out and they therefore cannot break electroneutrality.

One may include the double mutant in our model, however. By deleting two 7-valent positive surface charges, this mutant effectively flips the sign of the aerolysin wt charge. We therefore assume it can be modelled by placing a negative charge at the bottom of the channel, leading to positive gating (instead of negative gating in the aerolysin wt).

In light of the above remarks, and to simplify the discussion, we assume that all mutants can be modelled, in a first approximation, as a single effective charge located at some position $z_0$ which we leave as a free parameter to parametrize the different possible mutations. Pores that are overall neutral, like K242A and K238A, are assumed to have a zero effective charge. We show the graphs of the closed state probability as function of voltage phase for different values of $z_0$ in SI Figure 2j, reproducing the experimental phenomenology.

Correlation with rectification

One key aspect is how this model of gating connects with the one we derived for rectification, as both effects are caused by the charge distribution within the pore. Again, to keep the discussion simple, we model all mutants by a single charged group located at some position $z_0$, that contributes to both gating and rectification, plus a background of pairs of surface charges of opposite sign, that contribute to rectification through a constant rectification factor $\beta_{background}$, but not to gating. Gathering all results so far, the total rectification factor can be expressed as:

$$\beta(z_0) \simeq \beta_{background} - Q^* \frac{eV_0}{k_B T} \frac{L_p \ell_B}{2r_p^2} \sum_{n=1}^{\infty} \frac{(-1)^n}{(\kappa_D^2 L_p^2/4+\pi^2 n^2)\pi n} sin\frac{2\pi n z_0}{L_p}, \qquad \text{SI Equation 35}$$

as derived in a previous section (note that here we used the background, rather than aerolysin wt, as reference). The maximum closing rate is computed from:

$$\frac{dp}{dt} = \frac{1-p}{\tau_1} - \frac{p}{\tau_2}, \qquad \text{SI Equation 36}$$

with

$$\tau_1 = \tau e^{E_\ddagger} \qquad \text{SI Equation 37}$$

and

$$\tau_2 = \tau e^{E_\ddagger - E_{c,0} - F(V(t))}. \qquad \text{SI Equation 38}$$

The electrostatic free energy $F(V)$ itself is computed from its definition:

$$F(V) =- k_B T log\left[\sum_{n_Q=0}^{7} \binom{Q}{n_Q} e^{-E(n_Q,V)}\right], \qquad \text{SI Equation 39}$$

with

$$E(n_Q, V) =- e\frac{|V|}{2}(Q - n_Q) - e\frac{Vz_0}{L_p}(n_Q - Q) + E_0(n_Q - Q)^2 - \mu^{ex}(Q - n_Q). \qquad \text{SI Equation 40}$$

For each value of $z_0$, we compute $\beta(z_0)$ and $k_X(z_0) = \frac{dp}{dt}_{|max}$ assuming that the surface charge is positive (a negative surface charge at $z_0$ is equivalent to a positive charge at $-z_0$). For negative gating (defined as $\frac{dp}{dt}$ reaching its maximum under negative voltage), we replace $k_X$ (which is always positive by construction) by $-k_X$.

The model parameters are determined as follows. For rectification, we use $\beta_{background} = 0.12$ (estimated from several mutants that rectify but do not gate) – the other parameters are set by the pore's structure, or are physical constants. For gating, most parameters can be estimated from electrostatic considerations as detailed above: $E_0 = 7\,k_BT$, $\mu^{ex} = 5\,k_BT$, $E_{\ddagger} = 8\,k_BT$. The main unknowns are $E_{c,0}$ (energy of the closed state under 0 voltage; this value could be mutant-dependent), and the molecular attempt time $\tau$. The latter is particularly difficult to estimate. One solution is to consider the open-state conductance fluctuations across a single pore. The noise power spectrum becomes flat at frequencies $f > 1$ kHz, suggesting $\tau \lesssim 1$ ms, but it is unclear if that threshold should be attributed to pore fluctuations, intrinsic ion dynamics, or hardware noise. This timescale could also be mutant-dependent (pores with many mutations could become more unstable, leading to a smaller $\tau$).

In lack of a better choice, we used the values $\tau = 0.3$ ms and $E_{c,0} = 3\,k_BT$, which provided the best agreement with experimental data.

In Figure 4d of the main text, we represent our prediction for $k_X$ as function of $\beta$, and compare it to experimental data. The discontinuity around $\beta = \beta_{background}$ corresponds to pores that have a more or less symmetrical net charge distribution (which can be modelled by $z_0 = 0$, and gate both positively and negatively), or only present pairs of oppositely charged groups (which correspond to no gating at all).

As expected, not all mutants can be captured with this simplified model (where the charge distribution is essentially represented by a single point charge and a background of dipoles). In particular, the model underestimates gating in mutants with several charges of the same sign at similar locations (E258A, E254A, E258A254A). Similarly, the model fails to accurately describe pores with many mutations – the accumulated mutations potentially result in a complex charge landscape, and may affect the dynamics of the pore's structure.

Effect of other parameters

To assess the validity of our model, we investigate the effect of ionic concentration and temperature. Experimentally, we observed that the gating rate is slightly decreased at higher ionic strength, and increased at higher temperature.

In our model of gating, the ionic concentration mainly enters through depletion forces. In SI Figure 2c, we compare the energy landscape of the pore for a low (orange curve) and high ionic concentration (blue curve). While the energies of the open and the closed states are essentially unchanged, the energy of the transition state is greatly increased at higher concentration, as it reinforces depletion forces. The energy of the transition state sets the timescale of gating dynamics: our model therefore predicts that the gating rate decreases at high concentration. This result is in agreement with experimental findings (SI Figure 15), but the effect seems to be overestimated. A likely explanation is that we neglected ionic screening, which would result in lowering all electrostatic energies at high ionic strength, and therefore speed up gating dynamics.

In SI Figure 2k, we represent the closed state probability as a function of voltage phase at 300K as considered so far (continuous curves) and 330K (dashed curves). Counterintuitively, the model predicts that gating increases at high temperature, despite the relative energy difference (and therefore the associated Boltzmann weights) between the closed and open states being lower. This fact can be explained as follows. While increasing the temperature from 300K to 330K decreases the magnitude of the aforementioned energy difference (about $\pm\ 3\,k_BT$) by 10%, it also lowers the energy of the transition state (which is much higher, about $8\,k_BT$): gating is therefore slightly weaker, but also much faster. Overall, this results in an increase of the gating rate. This finding is in full accordance with experimental observations (SI Figure 15).

Lastly, our model does not account for ion specificity, and therefore does not reproduce the effect of ion species observed in experiments, where gating is weaker in aerolysin wt when using multivalent cations (SI Figure 15). Qualitatively, two effects could be distinguished. First, the valence of counterions could modify how easy it is to break lumen charge/counterion pairs. However, we observed a dependency of gating with the co-ion species (e.g. aerolysin wt, which has a positive charge, with either NaCl or MgCl2). This could stem from a second effect: the affinity of ions with the pore could modulate depletion forces, strengthening or weakening gating. These effects could be distinguished, for example, by investigating if a given ion species modifies gating similarly in all mutants (i.e. not dependent on whether that species is a co- or counterion for the lumen charge of a particular mutant) or not. However, we leave this discussion for future works.

### 2.5 ***Qualitative summary***

Gating phenomenology

We can now summarize the phenomenology that our gating model captures, and the underlying assumptions:

- Ions entering nanometric pores face an energy barrier that increases when the radius of the pore decreases.
- This barrier is at the source of depletion interactions: below a certain radius, maintaining the pore open is unfavorable and it is more favorable to collapse its structure and expel all ions.
- This effect induces an inherent bistability of the β-barrel, irrespective of any “movable” parts within the pore, like certain helices discussed in the gating of VDAC for example. A “simple” cylindrical pore is bistable in itself.
- The open state is unaffected by external electric fields, as these do not exert a net force on the pore (due to compensating electro-osmotic forces).
- In the closed state, however, a breakdown of electroneutrality can happen, breaking that compensation.
- Intuitively, if the external voltage increases beyond a certain value, it becomes favorable to “tear” a counterion from the surface charge, and put it at the mouth of the pore where it sees a favorable voltage. This process is very reminiscent of the Wien effect/ionic Coulomb blockade.
- This effect strongly depends on the sign of applied voltage. Let us consider the following: one applies a field such that the potential is positive at $z > 0$ and negative at $z < 0$, and that there is a negative surface charge surrounded by positive counterions at $z = z_0 > 0$. If a positive counterion is displaced from $z = z_0$ to $z = - L_p/2$, it will see a negative potential and the stripped negative surface charge

will see a positive voltage: both have a low energy. If now we reverse the direction of voltage, displacing a counterion from $z = z_0$ to $z =+ L_p/2$ is favorable, but this exposes a negative surface charge under a negative potential: it is overall still favorable, but barely.

- Breaking electroneutrality has an inherent cost (due to creating “uncompensated charges”), which increases rapidly with the number of counterions that “escape” the pore.
- Taking all of this together, a surface charge at one of the mouths of the pore would result in a strong gating in only one direction. A surface charge in the middle of the pore would result in weak gating in both polarities (the free energy gain by displacing counterions still exists, but is partially cancelled by the cost of breaking electroneutrality). A surface charge at an intermediate position would result in gating for mainly one voltage polarity, with a “secondary gating” in the other one.
- Gating is stronger for the voltage polarity that also corresponds to a conductance increase of the open state caused by ionic rectification.
- There are many different closed states, corresponding to different ways of breaking electroneutrality when the pore has a complex charge distribution.

Overall, this model is able to faithfully capture most of the phenomenology observed in experiments.

Limitations

Our model relies on a set of assumptions that we will now discuss. First, we used equilibrium statistical mechanics to study a system under an external driving force. This is partially justified by the fact that the ionic current through the closed pore is very low (albeit non zero), and the fact that counterions of the surface charges are likely localized, so that they can be identified to define the system’s microstates. Of course, these ions are constantly being replaced by new ones, so this picture should be taken with a grain of salt.

Another limitation is that we did not consider the effect of electrostatic interactions (beyond a binding energy between the surface charges and counterions, and an electrostatic energy barrier faced by charges entering the pore). In particular, we did not consider the effect of screening, which would likely amount to lowering the various electrostatic energy barriers considered here. While the concentration in the fluidic reservoirs, $c_0 = 1$ M, is very high, the average number of ions in the closed state is likely quite low, as ions have to pay a large free energy cost to enter the pore. This energy barrier is typically in the range $5 - 10\, k_B T$, meaning that only a couple ions probably permeate the closed pore (in absence of any surface charge), justifying that we neglect screening as a first approximation.

We also assumed we could model the surface charge distribution of each mutant by a single effective surface charge at some (mutant-dependent) location $z_0$, plus a constant background of electrostatic dipoles (corresponding to pairs of oppositely charged groups). This is clearly an oversimplification. However, accounting for more complex distribution would likely require better treatment of the electrostatic interactions between surface charges, or between ions. As our simplified picture already captures the essential experimental features, we leave this prospect for future works.

Lastly, one may ask if our model of gating could be extended to other pores than aerolysin. Here, we modelled cylindrical pores with no “movable parts” (such as flexible helices) bearing localized charge groups. One important assumption, both for gating and rectification, was that the pore’s radius ($r_p = 0.65$ nm in the open state) was small enough so that we can use a 1D model. This picture likely breaks down in bigger pores like MspA or α-HL, as the electrostatic screening length becomes comparable to the radius of the pore. These two pores are also not fully cylindrical, as they present a geometrical asymmetry. Such asymmetries generally have an important impact on rectification, which can therefore not be directly related to the charge distribution and gating.

Overall, however, this model sheds light on how gating can occur in β-barrel nanopores through a mechanical bistability, and how lumen charges can control this effect.

### Additional information

#### SI Section 3 Notes on working with aerolysin, MspA and α-HL

Mutating proteins is not an easy task and we have encountered limits by mutating several combinations of residues of aerolysin, generally all lower residues can be mutated even in combinations without impairing the function of the protein. Other residues can lead to pores with low membrane incorporation rates. Taking out all lumen charges in aerolysin has only worked so far when we do the additional mutations D209N and R288A. MspA has also been easy to mutate but the pores are harder to work with in experiments, as they need to be added together with detergents[78] which destabilises the membrane. Further, they tend to insert as dimers of octamers, which is likely to be tunable with detergent type and concentration. They are also much less inclined to insert into a reformed membrane with the same orientation. α-HL, which we mutated much less extensively, has not caused any troubles. We also need to mention that cross-linking cannot be readily done with aerolysin and α-HL since unlike MspA both pores do not form their multimeric full pore state in solution but undergo a major conformational change on the membrane to enter the pore which is prevented in aerolysin and α-HL pores treated with cross-linker. Likely there exists a pH optimum for each pore that will limit gating. For aerolysin wt that optimum should be slightly above the pH of 6.2 used in most of our measurements.

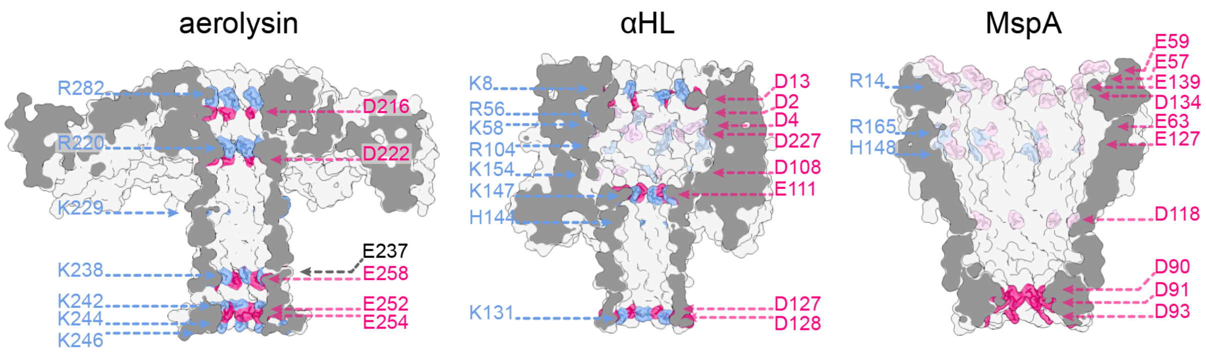

**SI Figure 3 | β-barrel pores and their lumen charges.** From left to right the wt of aerolysin, α-HL and MspA. All charges in the lumen are coloured in the structures with positive residues in blue and negative residues in red. Amino acid type and position in the peptide chain are indicated. Charges in regions that exhibit diameters larger than 2.5 nm carry a lighter shade of their respective colour. Charges outside of the lumen or stem regions are not shown.

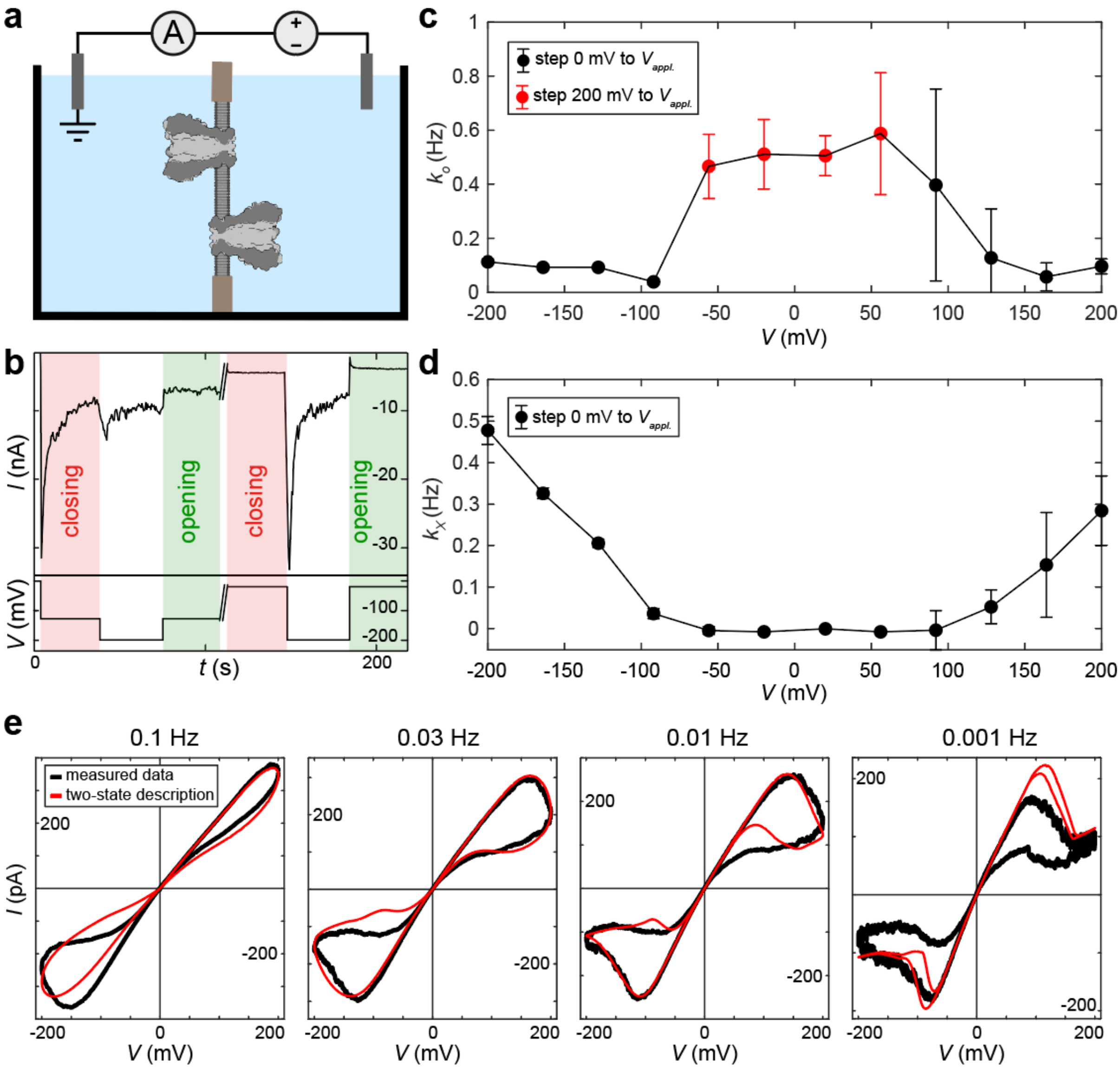

**SI Figure 4 | The memristive behaviour of an ensemble of pores can be described by individual pores that switch between two conductance states. a.** Schematic representation of the experiment with m2MspA pores inserted in both directions into the membrane. **b.** Example current recording used to extract opening and closing rates $k_O$ and $k_X$. When a bias voltage step of –128 mV was applied (first closing), the conductance of the ensemble decayed exponentially, and $k_O$ and $k_X$ were estimated from the parameters of this decay (see methods Section 4). In contrast, when a bias voltage step of –20mV is applied, pores hardly close (second closing), which is why the bias voltage was dropped to –200 mV for 35 seconds to close all pores, after which the bias was brought back to –20mV. From the resulting exponential increase in conductance (second opening), $k_O$ was estimated at bias voltages close to 0 (|V|<60mV). **c.** The resulting estimate of the mean opening rate $k_O$ at different bias voltages. Data points in red were estimated using a voltage step from ±200mV to the bias voltage (opening), whereas black points were estimated with a voltage step from 0mV to the bias voltage (closing). **d.** Estimate of the mean closing rate $k_X$ at different bias voltages. Error bars in **c** and **d** show twice the standard deviation of 3 repeat measurements done in the same pore ensemble. **e.** Linear interpolation of estimated rates of a single pore results in an accurate description of the ensemble gating behaviour, with $N_p$ as the only fitting parameter.

### SI Section 4 Two-state gating description

In order to extract voltage-dependent closing rates we used ensemble measurements at DC. Specifically, m2MspA inserted in both directions (SI Figure 4a). Applying a voltage step from $V$ = 0 to $V = V_{app.}$ results in an exponential decay in the ionic flux as the individual pores are closing (SI Figure 4b). From the decay parameter $\tau$, and the current at equilibrium A, the opening and closing rates $k_O$ and $k_X$ at that bias voltage are estimated (SI Figure 4c-d) (see methods Section 4). Since the formulation for $k_O$ diverged at bias voltages under 60 mV, a downward voltage step was used (SI Figure 4b), where pores are first closed at $V$ = ±200 mV and their opening (exponential increase in current) is observed at $V = V_{app.}$. We used linear interpolation of the extracted rates to make a continuous description of the switching rates of a single pore throughout one sinusoidal voltage cycle. In SI Figure 4e we then fit this continuous description to the experimental data at different frequencies, with the number of pores in the ensemble as the only fitting parameter. The description is accurate from 100 mHz down to 10 mHz below that the inaccuracy of the extracted closing rates becomes visible. The main problem is that the exponential decay is measured for only 35 s, after the elapse of which gating equilibrium is not reached for all voltages.

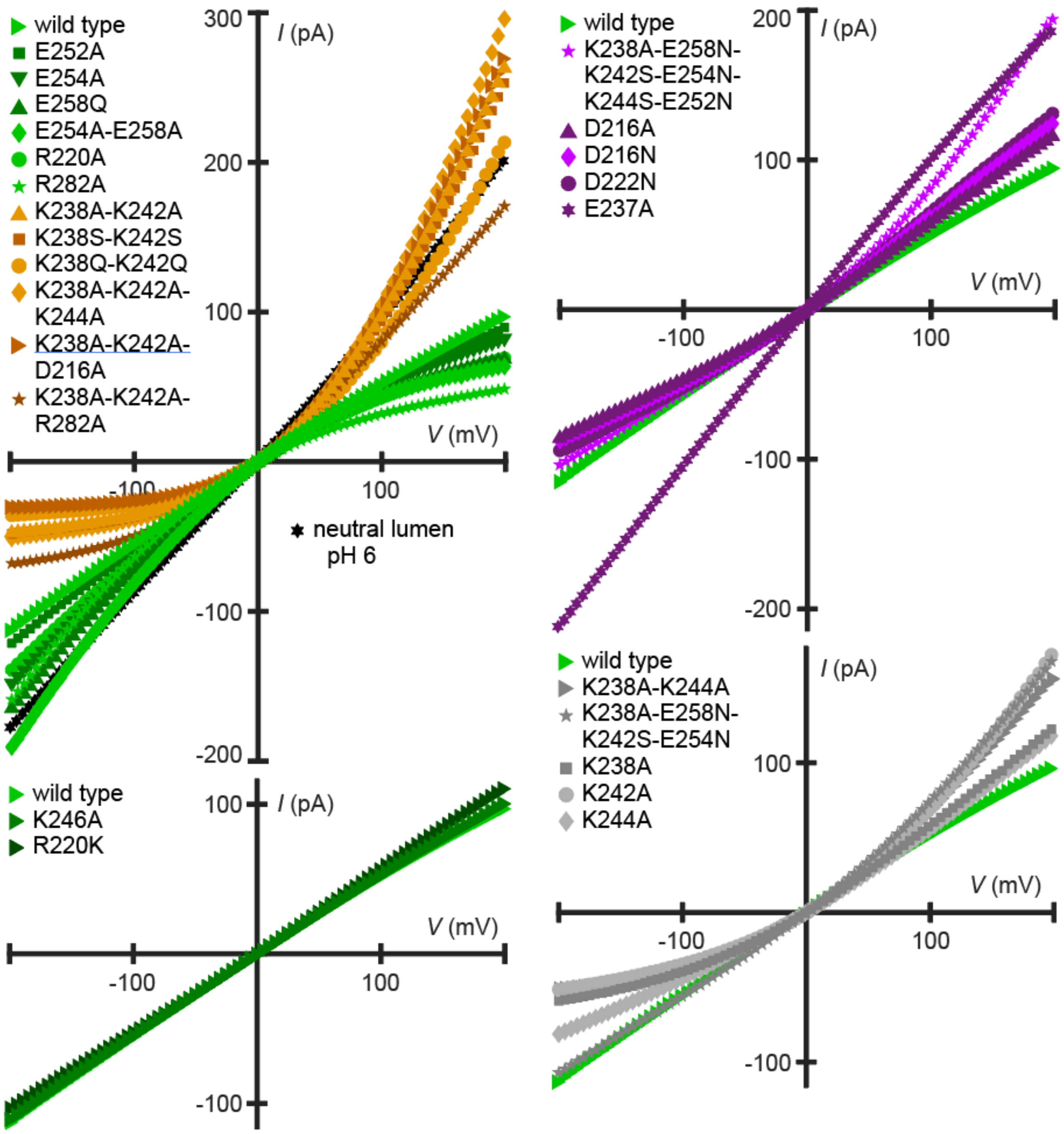

**SI Figure 5 | open-pore *IV* curves of mutants of aerolysin. top-left** open-pore *IV* curves for pores gating at only positive and only negative applied polarities. **top-right** open-pore *IV* curves for pores gating at both polarities. **bottom-left** open-pore *IV* curves for wt and wt-similar nanopores. **bottom-right** open-pore *IV* curves for pores exhibiting low gating. All plots also show aerolysin wt as a reference.

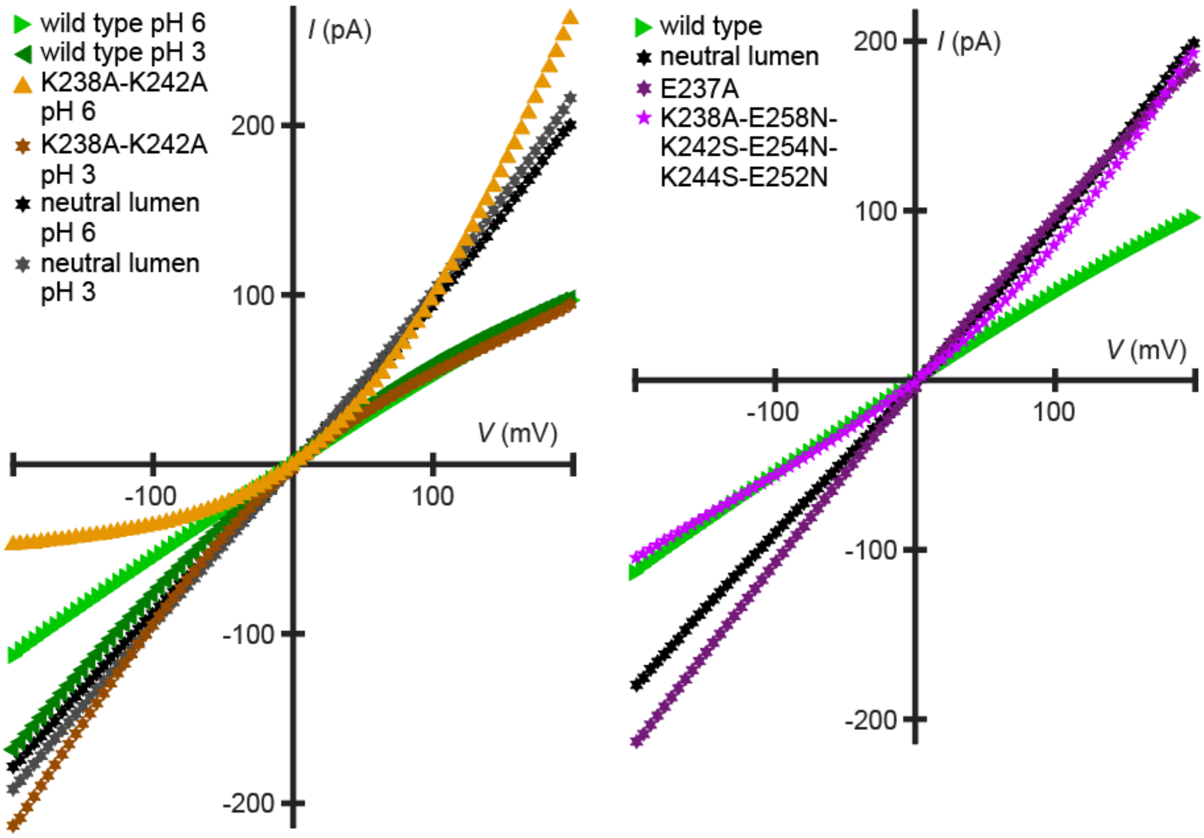

**SI Figure 6 | Highlighted open-pore *IV* curves of mutants of aerolysin. (Left)** open-pore *IV* curves of two mutants and wt for which the gating behaviour as a function of pH was investigated. **Right** open-pore *IV* curves of E237A, neutral lumen mutant, a mutant with the lower β-barrel charges removed and wt. The neutral lumen mutant exhibits the following mutations: K238A-E258N-K242S-E254N-K244S-E252N-R282A-D216N-R220A-D222N-K246A-D209N-R288A.

**SI Section 5 Rectification and conductivity of aerolysin, MspA and α-HL**

As we have shown that rectification of aerolysin is mainly modulated by its lumen charges this is also visible in the pH dependency of rectification where a mutant with titratable residues, which are not in a salt bridge where they are shielded, completely changes its rectification form concave to convex (SI Figure 6). One interesting observation is that E237A has the largest influence on the current through aerolysin (SI Figure 6 & SI Table 1), a finding that we attribute to the breaking of the hydrogen bonding network of E237 and Q263 but do not understand so far. The currents flowing through α-HL and its mutants are larger than the ones of aerolysin and the rectification is less strong (SI Figure 7 & SI Table 2). This can be attributed to the geometry factor that plays much more of a role than in aerolysin. The same is true for MspA where the rectification is even weaker due to its geometry (SI Figure 7 & SI Table 2).

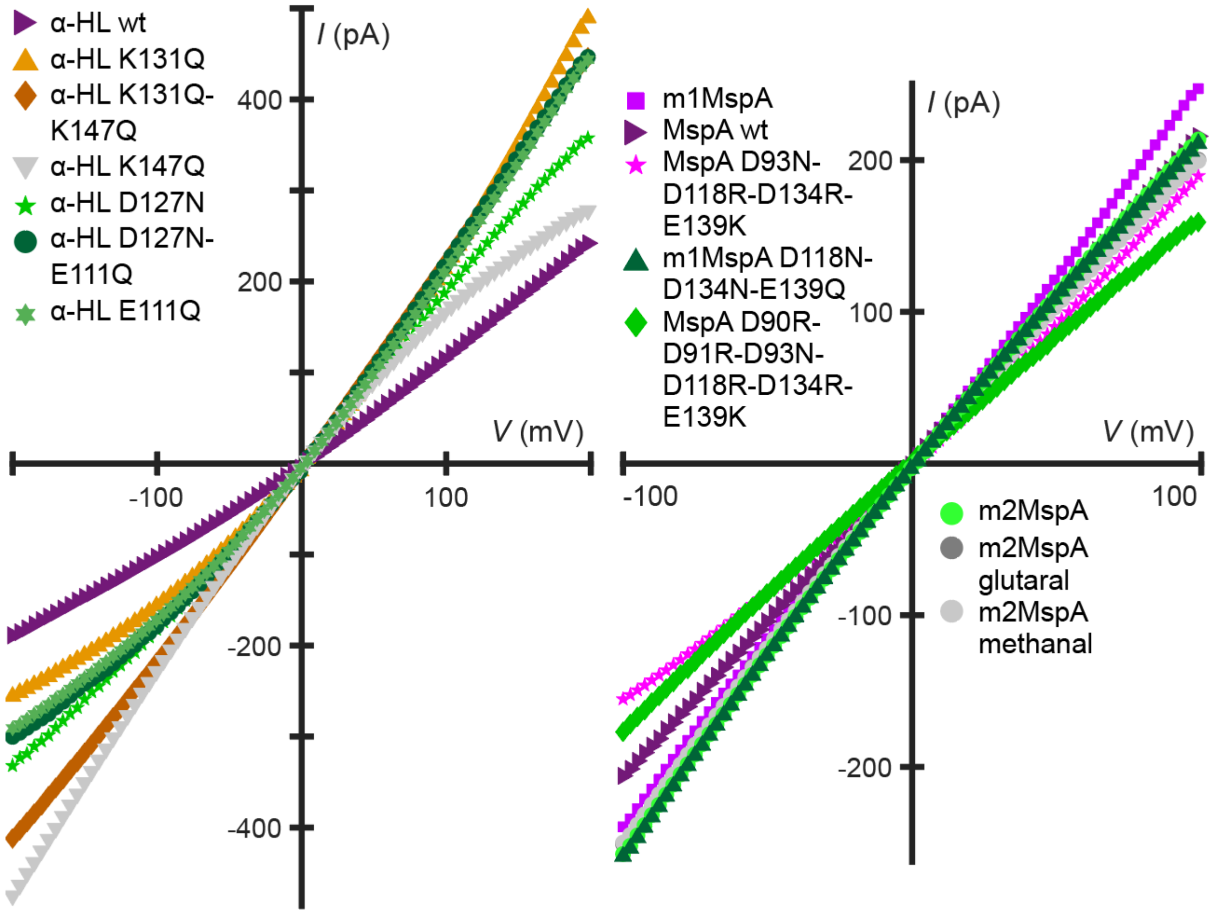

**SI Figure 7 | open-pore *IV* curves of MspA and α-HL and their mutants. left** MspA mutants **right** α-HL mutants.

**SI Table 1 |** Conductance at ±0.1 V as well as *β* and closing rates at both polarities for all mutants of aerolysin. Ordered by the conductivity at positive polarity.The neutral lumen mutant exhibits the following mutations: K238A-E258N-K242S-E254N-K244S-E252N-R282A-D216N-R220A-D222N-K246A-D209N-R288A.

| | K238A-K242A-K244A | K238A-K242A-D216A | neutral lumen mutant pH3 | K238A-K242A | E237A | K238S-K242S | neutral lumen mutant | K238A-E258N-K242S-E254N-K244S-E252N | K238A-K242A-R282A | K238Q-K242Q | K238A-E258N-K242S-E254N | K242A | K238A-K244A | D222N |
|---|---|---|---|---|---|---|---|---|---|---|---|---|---|---|
| $G_{0.1}$ (nS) | 1.08 | 1.01 | 1.01 | 0.99 | 0.99 | 0.96 | 0.94 | 0.82 | 0.82 | 0.81 | 0.75 | 0.70 | 0.70 | 0.66 |
| $G_{-0.1}$ (nS) | 0.35 | 0.24 | 0.88 | 0.33 | 0.97 | 0.33 | 0.82 | 0.50 | 0.45 | 0.26 | 0.51 | 0.32 | 0.35 | 0.47 |
| $\beta$ | 0.51 | 0.62 | 0.07 | 0.50 | 0.01 | 0.49 | 0.07 | 0.24 | 0.29 | 0.51 | 0.19 | 0.38 | 0.34 | 0.16 |
| $k_{X+}$ ($s^{-1}$) | 0.28 | 0.47 | 0.36 | 0.39 | 0.17 | 0.25 | 0.35 | 0.11 | 0.53 | 0.21 | 0.08 | 0.05 | 0.08 | 0.28 |
| $k_{X-}$ ($s^{-1}$) | | | | | -0.47 | | | -0.36 | | | -0.13 | -0.03 | -0.13 | -0.33 |

| | D216N | aerolysin wt pH3 | D216A | K238A | R220K | K244A | K238A-K242A pH3 | K246A | aerolysin wt | E254A | E252A | E258Q | E254A-E258A | R220A | R282A |
|---|---|---|---|---|---|---|---|---|---|---|---|---|---|---|---|
| $G_{0.1}$ (nS) | 0.63 | 0.59 | 0.59 | 0.59 | 0.57 | 0.56 | 0.54 | 0.53 | 0.52 | 0.51 | 0.51 | 0.45 | 0.44 | 0.43 | 0.32 |
| $G_{-0.1}$ (nS) | 0.45 | 0.70 | 0.43 | 0.36 | 0.48 | 0.40 | 0.86 | 0.50 | 0.50 | 0.62 | 0.53 | 0.66 | 0.73 | 0.59 | 0.58 |
| $\beta$ | 0.17 | -0.08 | 0.16 | 0.24 | 0.09 | 0.17 | -0.23 | 0.03 | 0.02 | -0.10 | -0.02 | -0.18 | -0.25 | -0.15 | -0.28 |
| $k_{X+}$ ($s^{-1}$) | 0.30 | | 0.33 | 0.02 | | 0.12 | | | | | | | | | |
| $k_{X-}$ ($s^{-1}$) | -0.78 | -1.00 | -0.57 | -0.03 | -0.45 | -0.13 | -0.50 | -0.34 | -0.43 | -1.29 | -0.64 | -0.71 | -2.19 | -0.55 | -0.45 |

**SI Table 2 |** Conductance at ±0.1 V as well as *β* and closing rates at both polarities for all mutants of MspA and α-HL. Ordered per pore by the conductivity at positive polarity.

| | m1MspA | MspA wt | m2MspA | m1MspA D118N-D134N-E139Q | m2MspA glutaral | m2MspA methanal | MspA D93N-D118R-D134R-E139K | MspA D90R-D91R-D93N-D118R-D134R-E139K | α-HL K131Q-K147Q | α-HL D127N-E111Q | α-HL K131Q | α-HL E111Q | α-HL D127N | α-HL K147Q | α-HL wt |
|---|---|---|---|---|---|---|---|---|---|---|---|---|---|---|---|
| $G_{0.1}$ (nS) | 2.47 | 2.14 | 2.12 | 2.09 | 2.00 | 1.99 | 1.88 | 1.59 | 2.23 | 2.20 | 2.19 | 2.14 | 1.91 | 1.68 | 1.18 |
| $G_{-0.1}$ (nS) | 2.36 | 2.03 | 2.54 | 2.56 | 2.49 | 2.47 | 1.54 | 1.74 | 2.19 | 1.79 | 1.57 | 1.72 | 1.82 | 2.28 | 1.03 |
| $\beta$ | 0.02 | 0.03 | -0.09 | -0.10 | -0.11 | -0.11 | 0.10 | -0.04 | 0.01 | 0.10 | 0.17 | 0.11 | 0.02 | -0.15 | 0.07 |
| $k_{X+}$ ($s^{-1}$) | 0.34 | 0.78 | 0.07 | 0.06 | 0.07 | 0.08 | 0.60 | 0.08 | 0.93 | 0.12 | 0.42 | 0.05 | 0.08 | 0.10 | 0.18 |
| $k_{X-}$ ($s^{-1}$) | -1.05 | -0.56 | -0.57 | -0.63 | -0.14 | -0.22 | -0.55 | -0.58 | -0.07 | -0.56 | -0.24 | -0.38 | -0.48 | -0.15 | -0.81 |

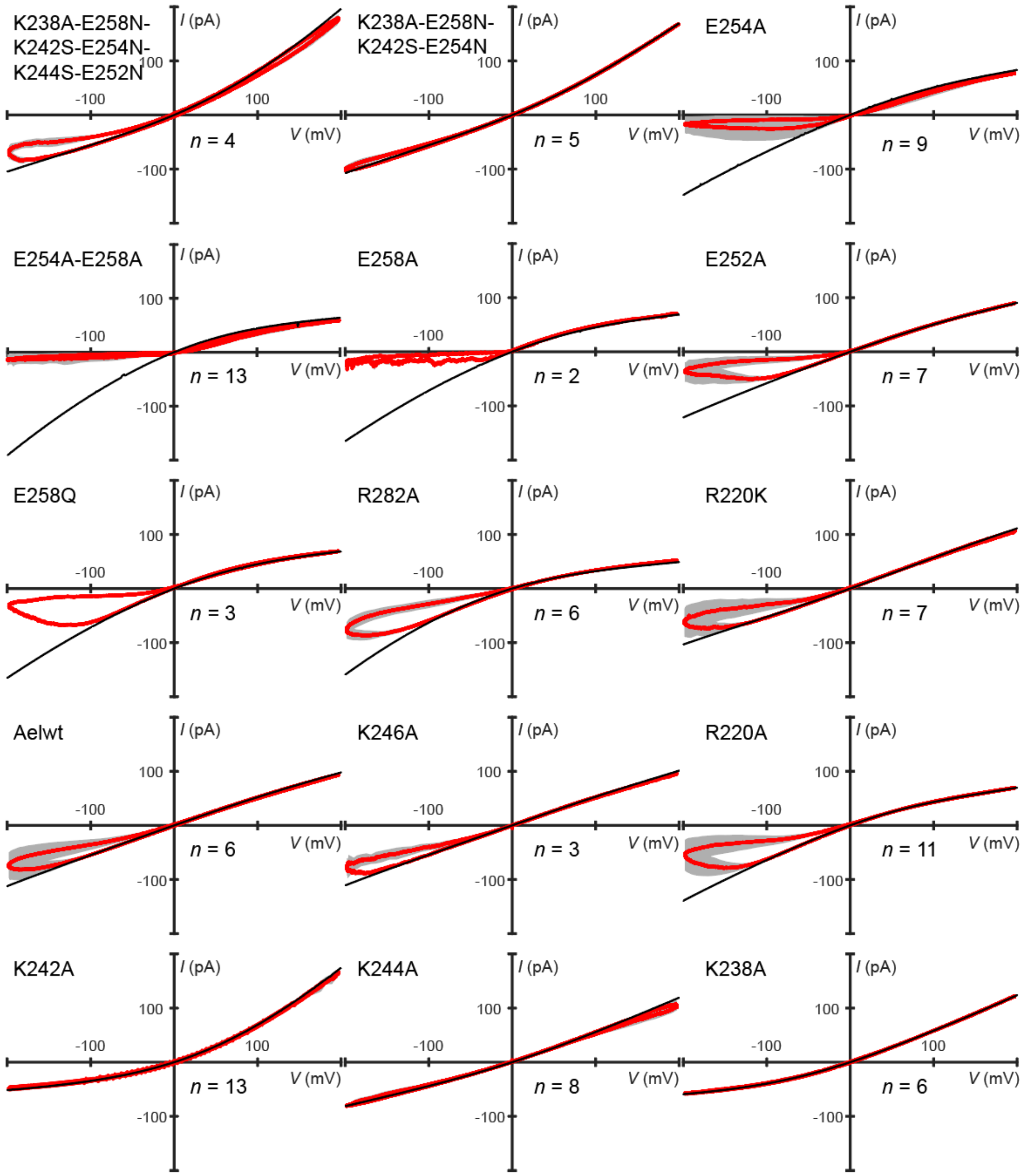

**SI Figure 8 | Aerolysin wt and mutants and their gating behaviour** at 0.1 Hz 200 mV amplitude. All curves are recorded at 25 °C, 1 M KCl, pH 6.2 buffered with 10 mM phosphate. Number of repeats *n* is indicated for each plot. Each repeat differs in the numbers of pores in the membrane and the amount of cycles recorded. In red is the average of all repeats. The open-pore *IV* curve is plotted in black. The grey area represents 2 standard deviations of different repeat experiments.

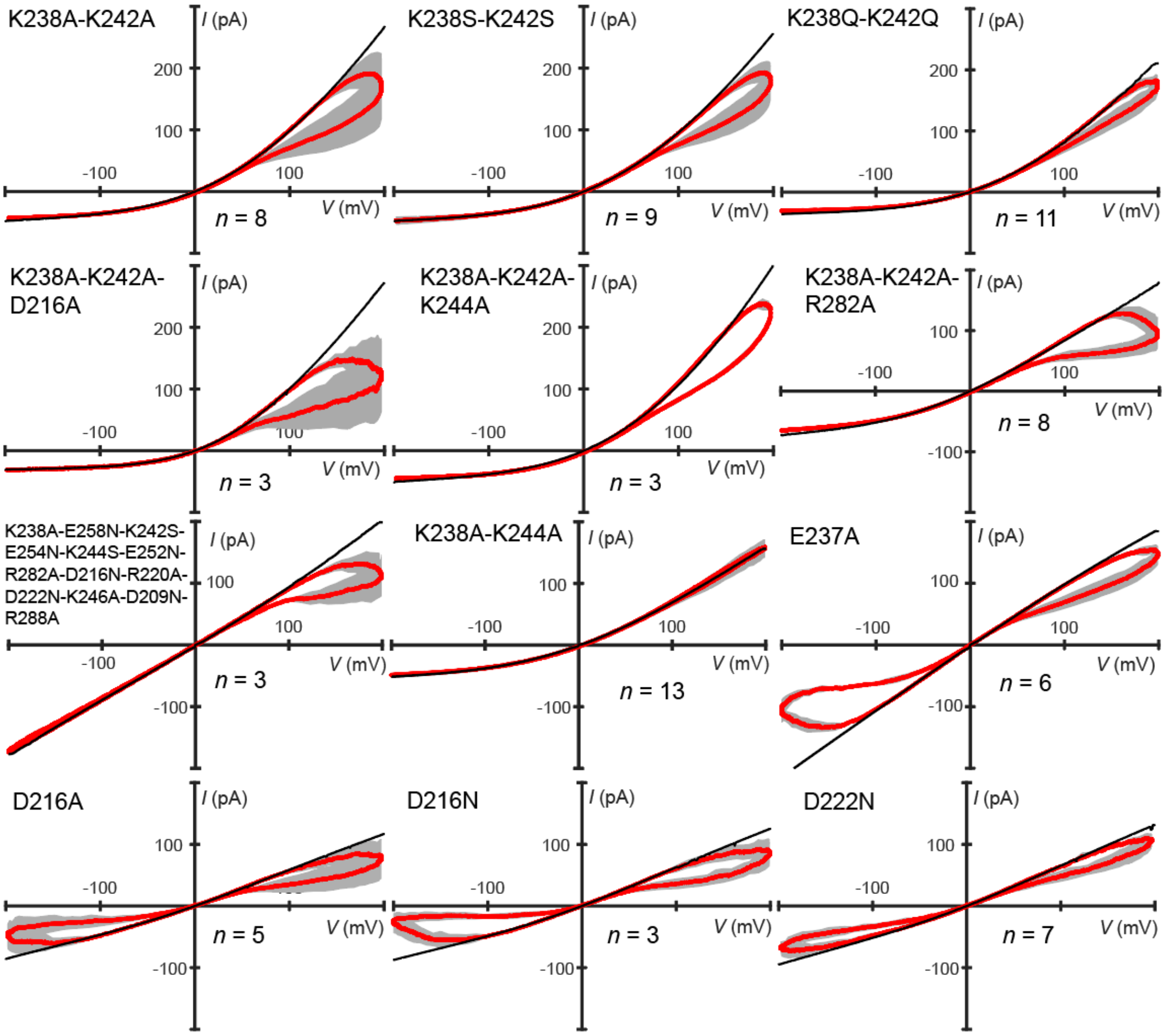

**SI Figure 9 | More aerolysin mutants and their gating behaviour** at 0.1 Hz 200 mV amplitude. All curves are recorded at 25 °C, 1 M KCl, pH 6.2 buffered with 10 mM phosphate. Number of repeats *n* is indicated for each plot. Each repeat differs in the numbers of pores in the membrane and the amount of cycles recorded. In red is the average of all repeats. The open-pore *IV* curve is plotted in black. The grey area represents 2 standard deviations of different repeat experiments.

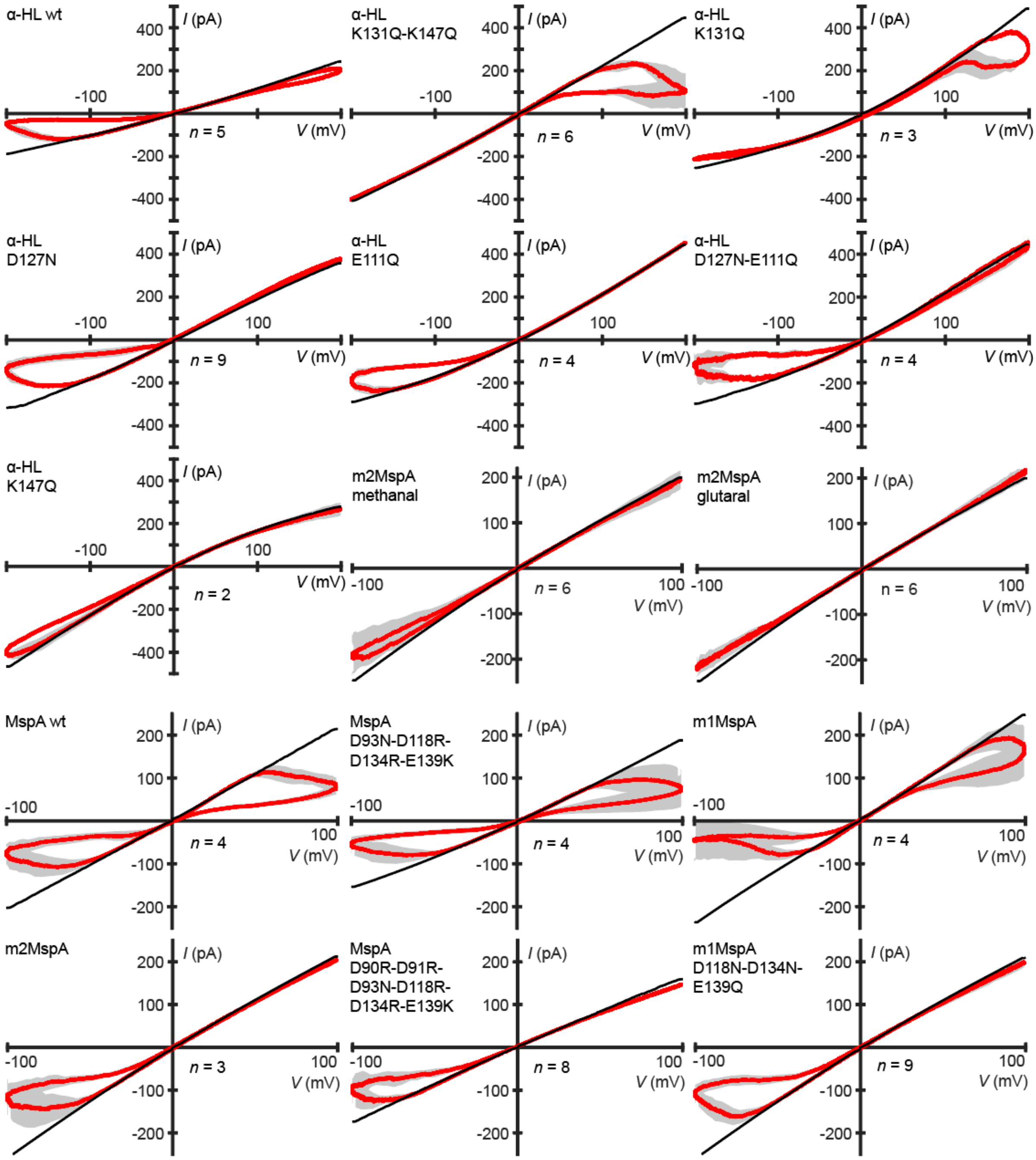

**SI Figure 10 | α-HL and MspA pores and their gating behaviour.** α-HL wt and mutants recorded at 0.1 Hz 200 mV amplitude. MspA wt and mutants recorded at 0.1 Hz and 100 mV amplitude. All *IV* curves are recorded at 25 °C, 1 M KCl, pH 6.2 buffered with 10 mM phosphate. The number of repeats *n* is indicated for each plot. The open-pore *IV* curve is plotted in black. The grey area represents 2 standard deviations of different repeat experiments.

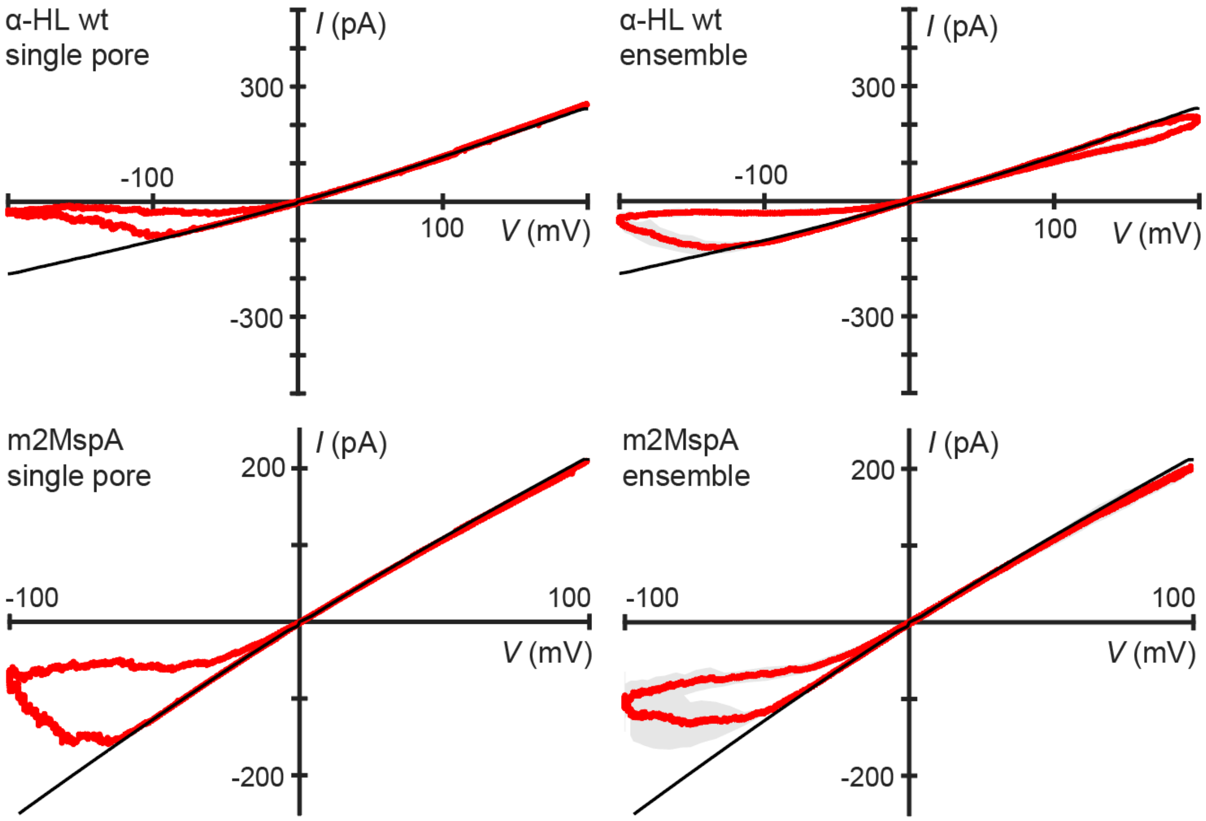

**SI Figure 11 | Ergodicity of α-HL wt and m2MspA.** The gating behaviour of 16 consecutive cycles of a single α-HL wt and of 66 consecutive cycles of a single m2MspA pore compared to their respective ensemble gating behaviour. α-HL wt recorded at 0.1 Hz 200 mV amplitude. m2MspA recorded at 0.1 Hz and 100 mV amplitude. All *IV* curves are recorded at 25 °C, 1 M KCl, pH 6.2 buffered with 10 mM phosphate. The open-pore *IV* curve is plotted in black. The grey area of ensemble experiments represents 2 standard deviations of different repeat experiments.

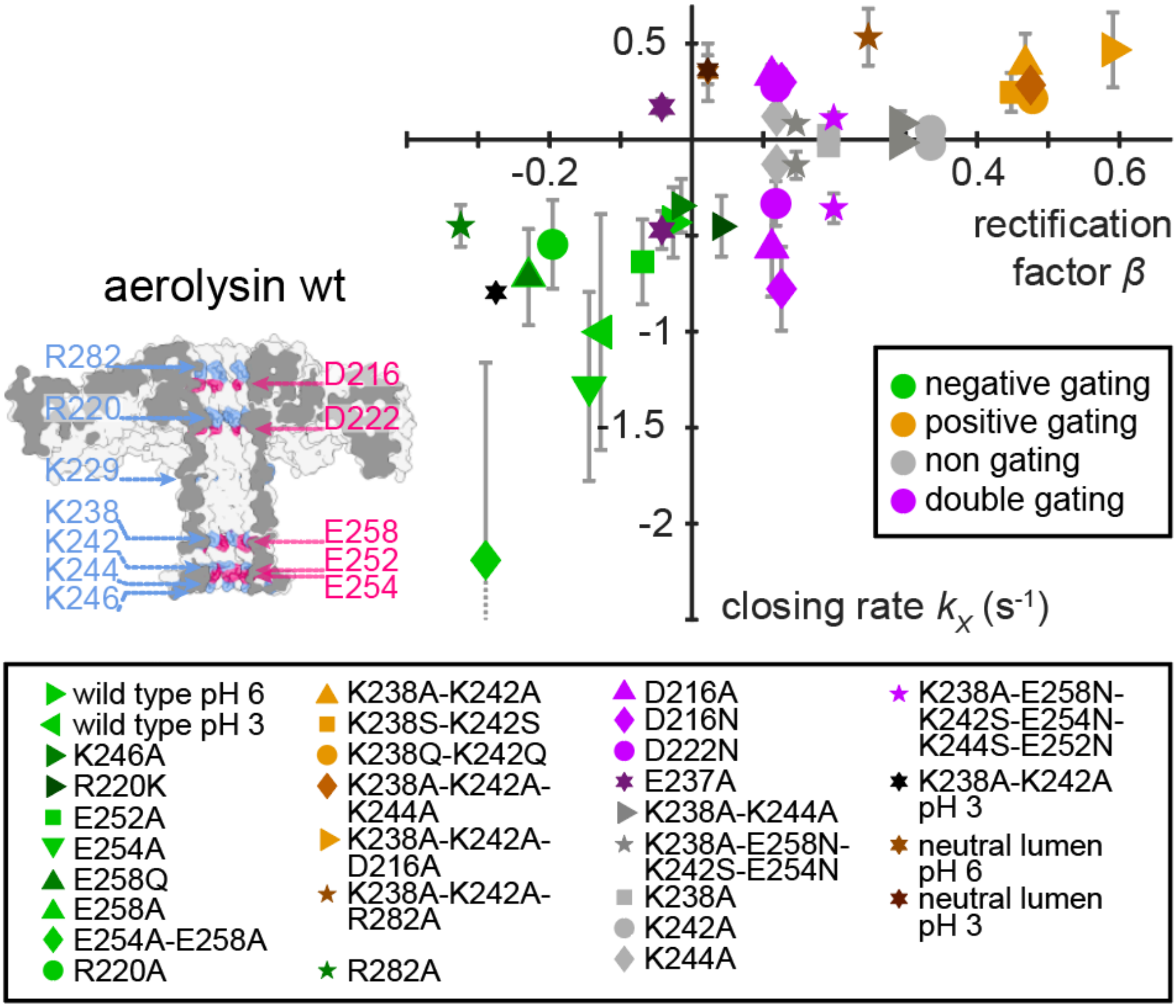

**SI Figure 12 |** Scatter plot of the pH response and all 26 aerolysin mutants' closing rates *versus* rectification factor *β*. Pores that show low gating or gating at both polarities are represented twice. For pores that show gating only in the negative or positive quadrant only one value is shown. The error bars show the standard deviation of closing rates of different repeat experiments. Each symbol represents a different mutant. The neutral lumen mutant exhibits the following mutations: K238A-E258N-K242S-E254N-K244S-E252N-R282A-D216N-R220A-D222N-K246A-D209N-R288A.

### SI Section 6 Gating dependencies

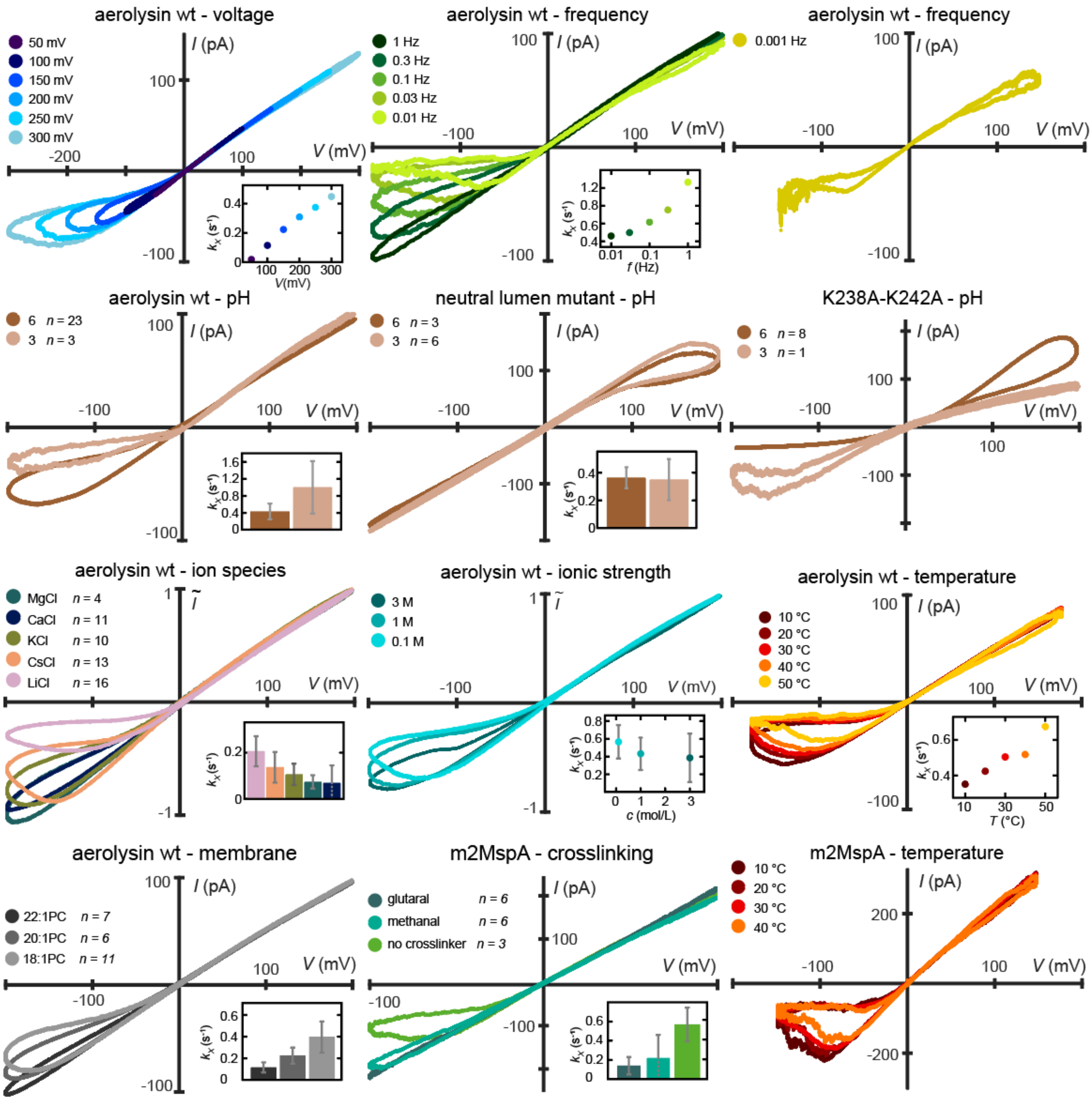

**SI Figure 13 | Gating as a function of voltage, frequency, pH, ion species, ionic strength and temperature.** Voltage, frequency and temperature dependencies are measured in the same experimental conditions and display of the number of experiments, *n*. The ionic species, ionic strength are recorded with 150 µm, aperture MECA-chips and the ionic species measurements are buffered with 10 mM MES instead of phosphate to a pH of 6.35 ±0.15 and are recorded at 0.03 Hz. Unless otherwise specified all recordings are done at 0.1 Hz, 1 M KCl, pH 6.2, 25 °C. In the inset the computed closing rate is shown. The neutral lumen mutant exhibits the following mutations: K238A-E258N-K242S-E254N-K244S-E252N-R282A-D216N-R220A-D222N-K246A-D209N-R288A.

We characterised the dependency of gating on frequency and amplitude. When changing the amplitude at a constant frequency, it becomes apparent that the closed-state probability increases with voltage. Further, when changing the frequency at a constant amplitude we can see that as expected the closing rate increases with frequency. Importantly, the lack of gating for aerolysin at positive voltages does not mean that aerolysin never gates at positive polarities, but rather that the closed-state probability is too low for the phenomenon to occur at the chosen conditions. At lower frequencies, the gating becomes visible also at positive polarities, this suggests a fundamentally symmetrical behaviour modulated by other factors[11]. We also measured the dependence of ionic strength, ion species and temperature on the gating behaviour. We found that temperature also influences gating, with an increase in gating at higher temperatures as previously shown for VDAC[54] and OmpF[53] This could come from the increased flexibility of proteins at higher temperature and would point towards a conformational change. As previously shown for α-HL[19] and lysenin[50], the gating becomes stronger with increasing Dukhin number[24],

$$Du = \frac{\Sigma}{r_p c F} \quad \text{(SI Equation 41)}$$

where $\Sigma$ is the pore net surface charge, $r_p$ its radius, $c$ the salt concentration and $F$ the Faraday constant. Such behaviour indicates that surface contribution plays a role in the gating process[24]. This observation is in line with the often charged pore lumina of biological nanopores (SI Figure 3). Since a change in pH should alter that charge, we also measured the pH dependency of gating in aerolysin and found that like in α-HL[18], lysenin[50], HlyA[50] OmpU and OmpT[52] a low pH indeed leads to stronger gating in aerolysin wt, K238A-K242A but not in the neutral lumen mutant which lacks de-/protonatable lumen charges. This clearly points to the lumen charge being the only influencing factor in the pH dependence of gating. We chose phosphate buffer to maintain a consistent buffer system across all tested pH values, as it provides three relevant buffering ranges. It is also apparent that the type of cation plays a role in the gating behaviour with lithium showing the strongest gating, followed by caesium, potassium, calcium and magnesium. Unlike previously shown for OmpF[20], in aerolysin gating does not follow the Hofmeister series. A valency dependence should not be concluded since previous experiments with zinc chloride show an increase in gating activity[79] but are otherwise consistent with our results.

**SI Section 7 The lipid membrane as another parameter influencing gating**

We realised that gating is dependent on the individual planar lipid membrane, we have observed that the magnitude of gating can vary between membranes. An extreme example of this is shown in SI Figure 14 where two membranes formed shortly after each other on one MECA-4 chip can show different degrees of gating. This has to do with the inherent diversity of painted and folded membranes. The oil that is used to anchor the membrane to its support influences membrane thickness, where shorter alkanes make for thicker membranes[80,81]. This thickness is hard to monitor accurately since membrane capacitance is the quotient of membrane area and thickness. However the variability can be decreased by increasing the size of the aperture over which the membrane is formed. By increasing the membrane area 9-fold by switching the aperture diameter of our membrane supports from 50 µm to 150 µm we minimise the variability and obtain consistent results between experiments. This was done for the membrane thickness, ion species, ionic strength and the pH dependency measurements of the neutral lumen and K238A-K242A mutant. For temperature, voltage and frequency dependencies this is not necessary since they are measured on one single membrane. For mutant measurements this was not done since the changes in gating behaviour are large and above the experimental variability.

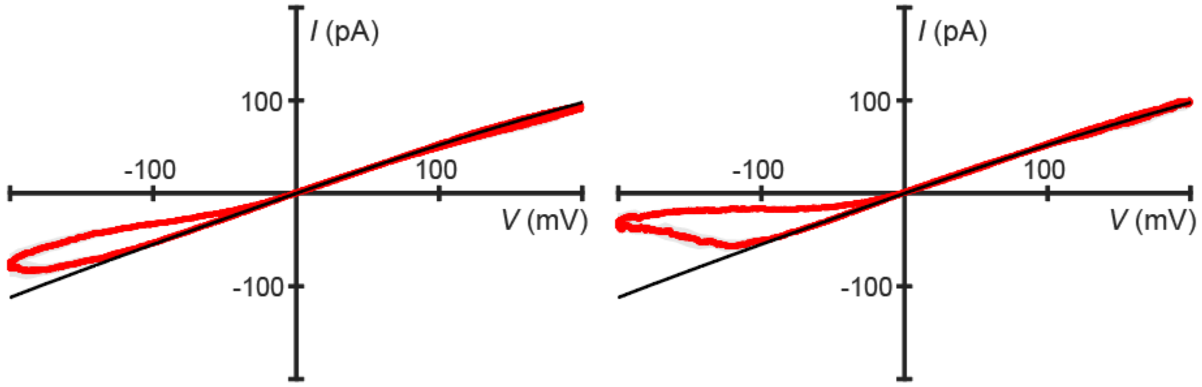

**SI Figure 14 | Extreme example of gating variability** on the same MECA-4 chip with 50 µm apertures, two membranes formed shortly after each other. The left experiment shows a lower gating than the one on the right. We avoid this through experimental repetition, averaging, as well as using larger membrane supports.

**SI Section 8 Long term memristor stability**

We ran experiments to assess the long term stability of our experiments. We were able to effortlessly run experiments for over 5 hours which were ended by a software issue. Despite the fact that the bilayer membrane is fragile, when voltage is limited to around 150 mV and the time that the membrane is exposed to higher potentials is optimized, these experiments can run for several hours. Further membranes can be stabilized mainly through the use of polymers[82] or polymer-lipid hybrid bilayers[83] or directly embedded complete with pores on commercially available chips as demonstrated by the flow cells sold by Oxford Nanopore Technologies.

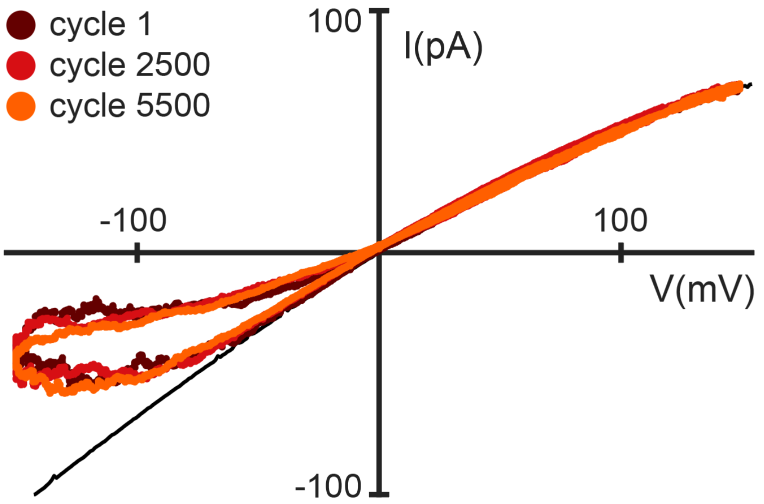

**SI Figure 15 | Long term memristor stability** E254A ensemble measurement run for over 5 hours at 0.3 Hz and 150 mV amplitude. Over 5600 cycles were measured. Here we show 3 of the cycles in the experiment.

### SI Section 9 Cryo-EM measurements

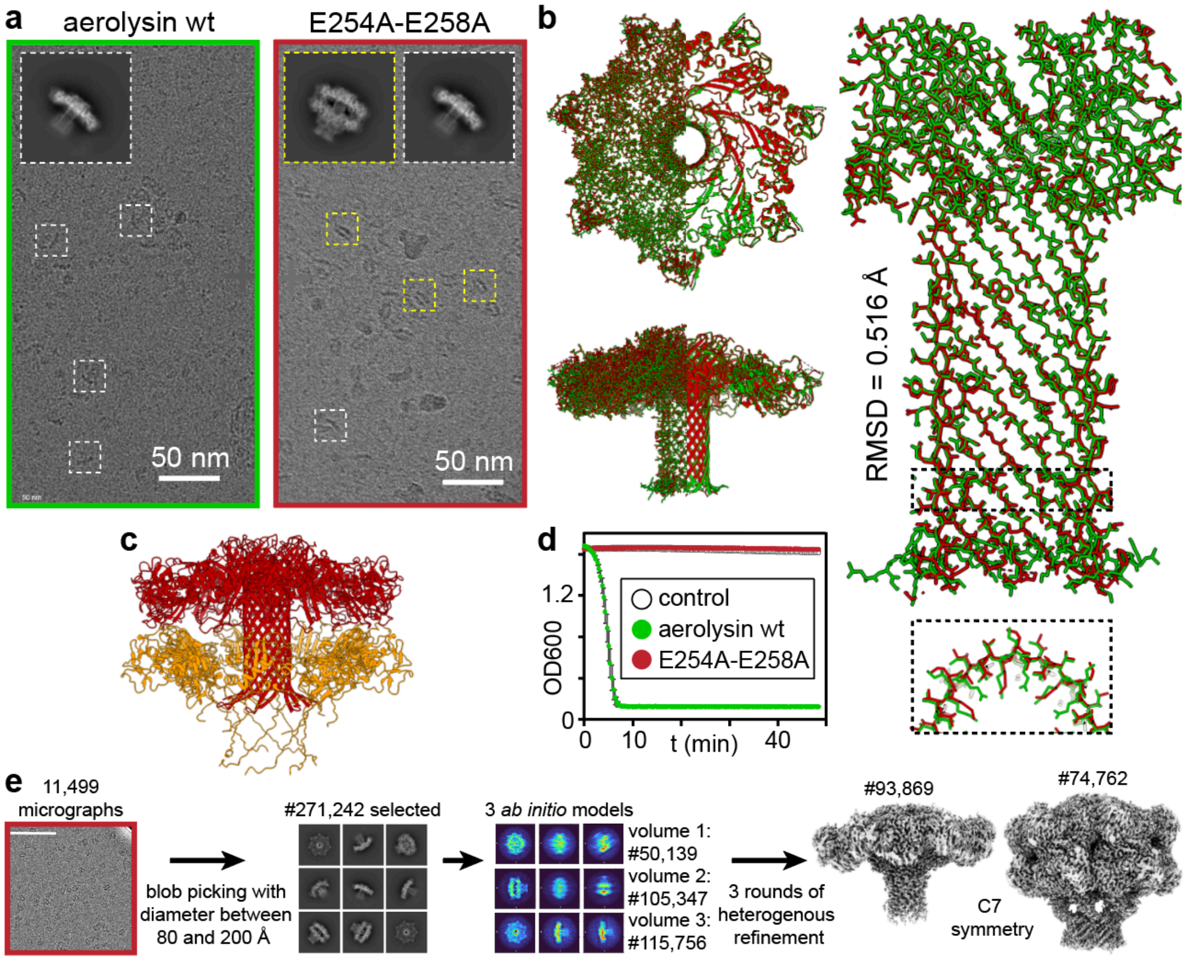

**SI Figure 16 | Cryo-EM structures of aerolysin wt and mutant E254A-E258A. a.** Representative cryo-EM grids of aerolysin wt and E254A-E258A (9GXJ) showing formed aerolysin pore particles as well as a double stack of prepore and pore (9IGN) as previously observed by Iacovache et al. for K246C-E258C[58]. **b.** Alignment of the structures of wt (green) and the K246C-E258C mutant (red, reconstructed from the population showing only one pore particle) showing no apparent difference in the overall structure RMSD = 0.52 Å. **c.** Refined structure of the double stacked particle visible in the K246C-E258C mutant with the prepore in orange and the formed pore in red. **d**. Blood assay showing the inability of the K246C-E258C mutant to lyse red blood cells. **e.** Flow chart of the data processing in cryoSPARC with the number of particles (#) indicated.

The strongest gating aerolysin nanopore that we have measured is the mutant E254A-E258A. By solving the cryo-EM structure of E254A-E258A and comparing it to the structure of aerolysin wt[39], we found that they are essentially the same (SI Figure 16b). Importantly, we further found that the mutant, unlike aerolysin wt, shows additional stacked particles formed by aerolysin pore within an aerolysin pre-pore, something that was already observed in the E258C-K246C mutant, where however the β-barrel instability were ad hoc engineered with the introduction of a disulfide bond[58]. This indicates that, while aerolysin wt univocally forms a mature β-barrel in multiple membrane-mimicking environments[39], the E254A-E258A mutant shows a much lower efficiency for β-barrel formation as roughly half of the particles observed are in the pre-pore conformation. As pore formation in aerolysin is thought to happen in the presence of a membrane via a two-step mechanism (monomers to pre-pore to mature pore[84]) which essentially goes downhill in free energy, this may imply that the free energy landscape of pore formation of E254A-E258A is perturbed and its pores are likely to be less stable than wt. This can have implications for the enhanced observed gating behaviour of E254A-E258A and draws a link between mutations changing the electrostatics of the transmembrane portion of the β-barrel, the ability to form stable pores and therefore the appearance of gating.

**SI Table 3 | CryoEM maps processing and model refinement statistics**

| Protein<br>**Acess codes** | **E254A-E258A pore** | **E254A-E258A quasipore & post-prepore** |
|---|---|---|
| PDB | 9GXJ | 9IGN |
| EMDB | EMD-51664 | EMD-52853 |
| **Data collection and processing** | | |
| Microscope | TFS Titan Krios G4 | TFS Titan Krios G4 |
| Detector | Falcon IV | Falcon IV |
| Recording mode | EC | EC |
| Magnification | 120'000 | 120'000 |
| Voltage (kV) | 300 | 300 |
| Total dose ($e^-/Å^2$) | 50 | 50 |
| Nominal under focus range (µm) | 0.8 - 1.7 | 0.8 - 1.7 |
| Pixel size (Å) | 0.658 | 0.658 |
| # movie micrographs | 11'499 | 11'499 |
| # molecular projection images in map | 95'488 | 115'756 |
| Symmetry | C7 | C7 |
| Map resolution (Å) | 2.30 | 2.25 |
| Map sharpening B-factor | 66.2 | 63.6 |
| **Model refinement and validation** | | |
| Residues | 2'842 | 4'970 |
| RMSD Bond Length (4σ) | 0.004 | 0.005 |
| RMSD Bond Angles (4σ) | 0.607 | 0.707 |
| Ramachandran Outliers (%) | 0 | 0 |
| Ramachandran Allowed (%) | 3.24 | 2.98 |
| Ramachandran Favored (%) | 96.75 | 97.02 |
| Rotamer outliers (%) | 2.73 | 2.23 |
| Clash score | 3.95 | 3.21 |
| MolProbity score | 1.7 | 1.55 |
| EMRinger score | 6.06 | 6.43 |

**SI Section 10 Gating is unlikely to stem from contaminations**

Contaminations have been mentioned as a possible cause of gating. However, it would be highly unlikely that an effect caused by contaminations is observed universally and consistently across different laboratories and times. Further, since gating duration increases with voltage the contaminant would need to be trapped in the pore, unable to translocate; this by itself is only explainable with a highly charged contaminant which should block the pore independently of polarity. Since gating is highly dependent on polarity and can be tuned with lumen charges, it is reasonable to exclude contaminants as a cause of gating.

### SI Section 11 Molecular Dynamics Simulations

Observing gating in MD simulations is hard since the effect occurs on the order of seconds, a timescale unreachable in MD simulations to this day. We set out to try with higher voltages but were unable to observe gating. Ultimately we used lower voltage simulations (±50 and ±200 mV) to probe the effects of pore lumen charges on the ion distributions in the pore (SI Figure 18). We checked the validity of our simulations by calculating the currents and found that they are in good agreement with experimental data (SI Figure 17). The ion distributions can be seen to be polarity-dependent. Ions move through the pore biassed by the external and local charge induced electric fields (SI Figure 18).

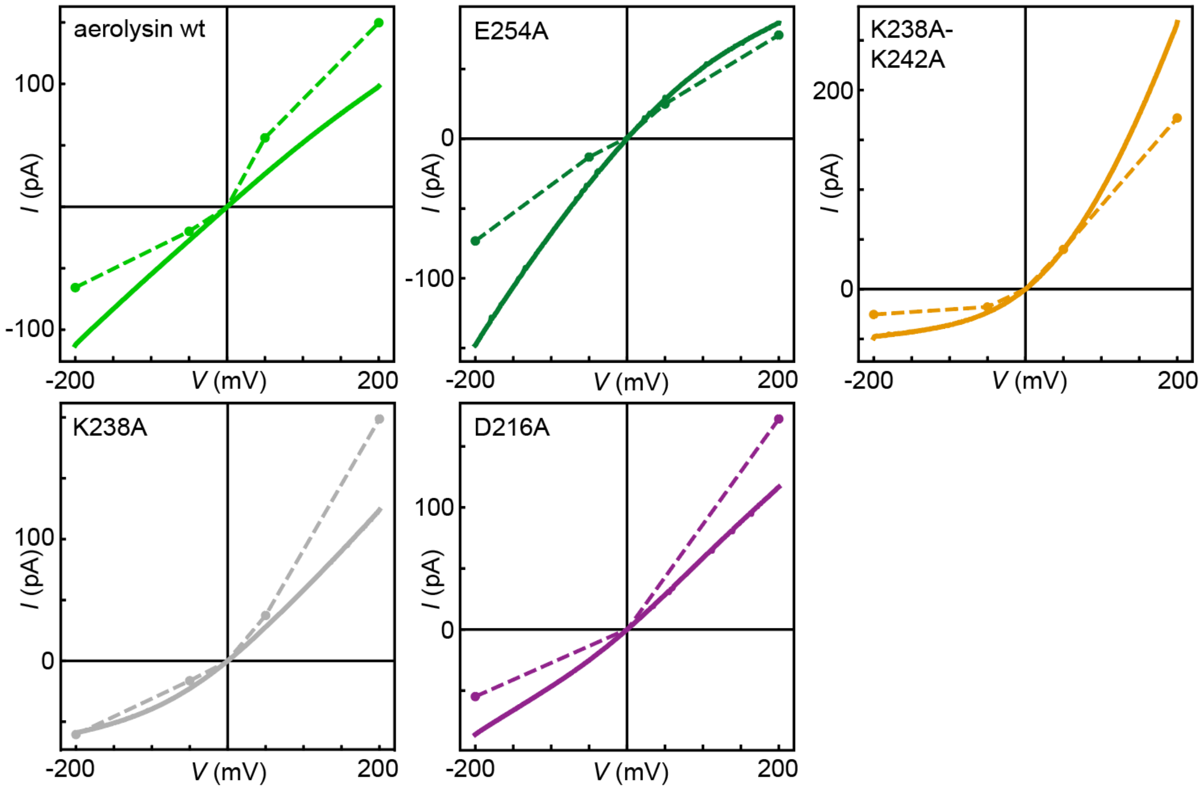

**SI Figure 17 | Simulated *versus* experimental results**. Comparison between measured open-pore *IV* curves (solid lines) with *IV* curves calculated from simulations (dashed lines) The simulation data show good agreement with the measured values, demonstrating the accuracy of the simulations.

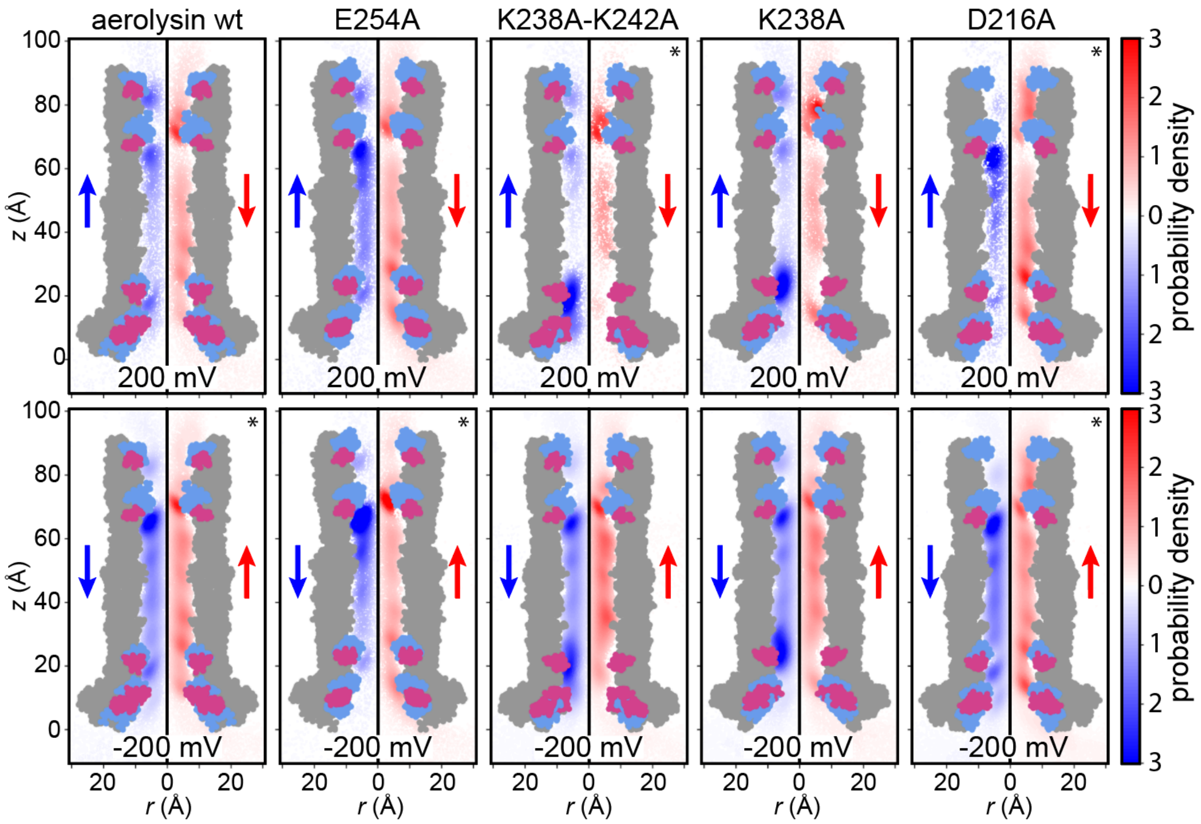

**SI Figure 18 | Ion radial probability densities measured on the MD simulations of aerolysin wt and selected mutants inserted in DPhPC bilayers under the indicated voltages.** Chloride ions in red, potassium in blue, and their respective directions of flow are indicated with arrows. The charged residues inside of the pore (grey) are coloured by their respective charge, negative (red) positive (blue). The upper row of plots represents the simulation at 200 mV the lower row the simulation at -200 mV. Only the translocating ions are identified and plotted. The polarity and mutant-dependent changes in ion distributions and densities is subtle but visible.

### SI Section 12 Conductance at different salt concentration

We report in SI Figure 19 the conductance of aerolysin wt against the salt concentration. We observe a saturation toward small concentrations pointing to a transition from bulk to surface regime. Consequently we express the conductance as:

$$G_0 = K \cdot \frac{\pi r_p^2}{L_p} \cdot 2\mu_e F c \cdot \sqrt{1 + Du^2} \qquad \text{SI Equation 42}$$

Where $\mu_e$ is the electrophoretic mobility, $c$ is the salt concentration, $r_p$ is the pore radius and $L_p$ its length and $K$ is a unitless coefficient taking into account the energy barrier. In the square root, the first term represents bulk contribution and the second one is the surface contribution[85]. By fitting the conductance against concentration, and solving SI Equation 42 we obtain $\Sigma$ = 9±5 mC/m$^2$ and a coefficient $K$ = 0.21 ±0.01. Assuming that charge is uniformly distributed, this fitted surface charge value translates into 2.9 ±1.5 charges.

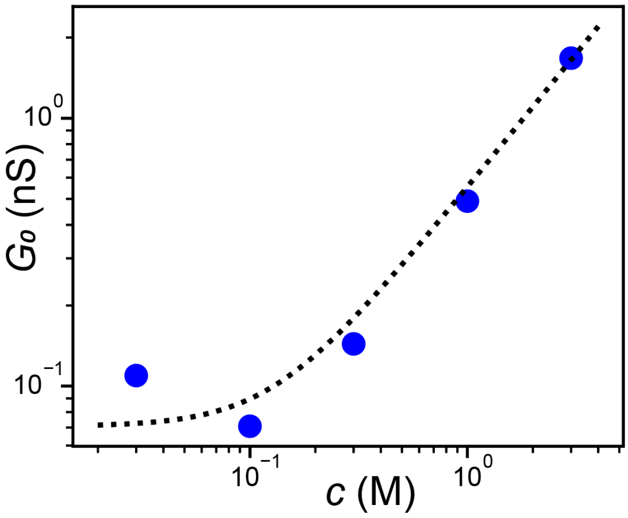

**SI Figure 19 | Conductance vs salt concentration (KCl) of aerolysin wt.** The data (blue dots) is fitted to SI Equation 42, the fit (dotted line) is shown. Assuming $L_p$ = 9 nm and $r_p$ = 0.65 nm, we extract a charge of 4 mC/m$^2$ and a $K$ coefficient of 0.5. The conductance presented is an average of the conductance at 200 mV and -200 mV.
